# Interaction of high-fat diet and brain trauma alters adipose tissue macrophages and brain microglia associated with exacerbated cognitive dysfunction

**DOI:** 10.1101/2023.07.28.550986

**Authors:** Rebecca J. Henry, James P. Barrett, Maria Vaida, Niaz Z. Khan, Oleg Makarevich, Rodney M. Ritzel, Alan I. Faden, Bogdan A. Stoica

**Author notes:** **Corresponding Author:** Rebecca J. Henry: Department of Anatomy and Neuroscience, University College Cork, Ireland. equal contribution (James P. Barrett, Maria Vaida). equal contribution (Alan I. Faden, Bogdan A. Stoica).

## Abstract

Obesity increases the morbidity and mortality of traumatic brain injury (TBI). We performed a detailed analysis of transcriptomic changes in the brain and adipose tissue to examine the interactive effects between high-fat diet-induced obesity (DIO) and TBI in relation to central and peripheral inflammatory pathways, as well as neurological function. Adult male mice were fed a high-fat diet (HFD) for 12 weeks prior to experimental TBI and continuing after injury. Combined TBI and HFD resulted in additive dysfunction in the Y-Maze, novel object recognition (NOR), and Morris water maze (MWM) cognitive function tests. We also performed high-throughput transcriptomic analysis using Nanostring panels of cellular compartments in the brain and total visceral adipose tissue (VAT), followed by unsupervised clustering, principal component analysis, and IPA pathway analysis to determine shifts in gene expression programs and molecular pathway activity. Analysis of cellular populations in the cortex and hippocampus as well as in visceral adipose tissue during the chronic phase after combined TBI-HFD showed amplification of central and peripheral microglia/macrophage responses, including superadditive changes in select gene expression signatures and pathways. These data suggest that HFD-induced obesity and TBI can independently prime and support the development of altered states in brain microglia and visceral adipose tissue macrophages, including the disease-associated microglia/macrophage (DAM) phenotype observed in neurodegenerative disorders. The interaction between HFD and TBI promotes a shift toward chronic reactive microglia/macrophage transcriptomic signatures and associated pro-inflammatory disease-altered states that may, in part, underlie the exacerbation of cognitive deficits. Targeting of HFD-induced reactive cellular phenotypes, including in peripheral adipose tissue macrophages, may serve to reduce microglial maladaptive states after TBI, attenuating post-traumatic neurodegeneration and neurological dysfunction.

## 1. INTRODUCTION

Traumatic brain injury (TBI) is a debilitating disorder that affects more than 1.5 million individuals annually in the United States [1]. Obesity is among the most prevalent pre-existing conditions that can negatively impact TBI outcomes. Currently, approximately 13% of the world population is considered clinically obese [body mass index BMI greater than 30kg/m^2^] [2] placing a large burden on healthcare systems [3]. Emerging evidence indicates that obese patients have increased TBI complications and higher mortality rates (36%) [4–7].

Activation of brain and peripheral inflammation pathways is a key pathophysiological feature in both TBI [8–12] and obesity [12–20] and has been implicated in the development of associated neurological dysfunction [8, 21, 22]. Notably, evidence suggests that pre-existing diet-induced obesity is associated with an amplification of post-TBI pro-inflammatory responses and increased microglia-altered states in several brain regions [23–25]. Additional studies report diet-induced exacerbations in TBI-induced cognitive decline [24, 25] and central (brain) insulin resistance [26].

Microglia play a key role in mediating the inflammatory responses after TBI. In clinical and experimental TBI, microglia have been shown to undergo chronic reactive changes that may contribute to long-term neurodegenerative processes and cognitive decline [10, 11]. In animal models, pharmacological depletion of microglia or other interventions that reduce chronic microglial activation decrease TBI-induced neuroinflammation and associated neurological deficits [8, 27, 28]. Microglia also mediate hypothalamic inflammation and neuronal stress responses in diet-induced obesity [19]. Selective pharmacological inhibition of microglial phagocytosis limits diet-induced obesity hyperphagia and weight gain, decreases dendritic spine loss, and reduces cognitive dysfunction [21].

Obesity triggers peripheral inflammation that includes reactive changes in adipose tissue macrophages (ATMs), which account for >50% of the cells in the increased visceral adipose tissue (VAT) [29]. VAT is an immunogenic tissue and the resulting state of obesity-induced chronic low-grade inflammation and increased secretion of pro-inflammatory factors into the circulation contribute to brain neuroinflammatory responses [29, 30]. Obesity is also associated with increased cognitive decline and dementia [31–33]. Inflammation may play a role in driving the negative pathological consequences of diet-induced obesity, including behavioral deficits [34–37]. NLR family, pyrin domain-containing 3 (NLRP3) contributes to obesity-induced inflammation [36, 38] through NLRP3-induced activation of microglial IL-1 receptor 1 (IL-1R1) [36]. Targeted knockout of NLRP3 at the level of the VAT attenuates diet-induced obesity cognitive deficits; thus, implicating a potential role for NLRP3/IL-1β signaling in brain-visceral adipose tissue interactions related to obesity [36]. Furthermore, the depletion of myeloid NOX-2, a key pro-inflammatory molecule, decreases VAT inflammation, VAT macrophage infiltration, and cognitive dysfunction [37].

HFD-induced obesity may promote the development of VAT macrophage reactive states that prime the development of deleterious brain microglia phenotypes, increasing vulnerability to secondary insults including TBI. Thus, the amplification of chronic microglial maladaptive states in combined TBI-HFD may contribute to the observed exacerbation of cognitive dysfunction [13, 23]. The goal of the present study was to use transcriptomic analysis to examine whether diet-induced obesity and TBI interact both centrally (brain) and systemically (VAT) to promote altered disease-driving microglia/macrophage states that may negatively impact neurological function.

## 2. MATERIALS & METHODS

### 2.1 Animals

Studies were performed using adult male C57Bl/6J mice (8-10 weeks old; Jackson Laboratories). Mice were housed in shoebox cages (5 mice in each cage) at least 1 week prior to any procedures in a room (22–23°C) with a 12-h/12-h light-dark cycle. Food and water were provided *ad libitum*. The mice were handled briefly before use. Procedures were conducted from 10:00 to 17:00 in a quiet room. All protocols involving the use of animals complied with the Guide for the Care and Use of Laboratory Animals published by the National Institutes of Health (NIH) (DHEW publication NIH 85-23-2985) and were approved by the University of Maryland Animal Use Committee.

### 2.2 Experimental Design

To induce our model of DIO, C57Bl/6 mice (n=8-12/group) were placed on either a high-fat diet (HFD) (60 kcal% Fat; D12492, Research Diets) or a standard diet (SD) (10 kcal% Fat; D12450B, Research Diets) for a period of 12 weeks prior to exposure to controlled cortical impact (CCI) or Sham surgery.

#### Cohort 1

Following exposure to CCI/Sham surgery, mice were anesthetized (100 mg/kg sodium pentobarbital, I.P.) and transcardially perfused with ice-cold 0.9% saline (100ml) at 28 days post-injury (dpi). VAT, brain tissue, and blood samples were collected and stored at −80°C until processed for RNA/protein analysis.

#### Cohort 2

To assess more chronic effects of diet and/or TBI, on neurobehavioral, neuroinflammatory and neurodegenerative outcomes, a second cohort of mice was fed a SD or HFD diet for 12 weeks prior to induction of CCI/Sham surgery. Mice underwent a battery of neurobehavioral tasks to assess cognitive behaviors throughout 90 dpi. Mice were anesthetized (100 mg/kg sodium pentobarbital, I.P.) and transcardially perfused with ice-cold 0.9% saline (100ml) at 90 dpi. Cluster of differentiation 11b (CD11b) positively selected cells were isolated from hippocampal and cortical tissue using Miltenyi MACS Cell separation. RNA was extracted from both the CD11b positively selected cells (primarily microglia but also resident/infiltrating macrophages) as well as flowthrough (the remaining brain cells, including neurons, astrocytes, and oligodendrocytes) and analyzed using a Nanostring Glial Panel. RNA was also extracted from the VAT and analyzed using a Nanostring Neuroinflammation panel.

### 2.3 Controlled Cortical Impact

Our custom-designed controlled cortical impact (CCI) injury device consists of a microprocessor-controlled pneumatic impactor with a 3.5 mm diameter tip. Mice were anesthetized with isoflurane evaporated in a gas mixture containing 70% N_2_O and 30% O_2_ and administered through a nose mask (induction at 4% and maintenance at 2%). The depth of anesthesia was assessed by monitoring respiration rate and pedal withdrawal reflexes. Mice were placed on a heated pad, and core body temperature was maintained at 37°C. The head was mounted on a stereotaxic frame, and the surgical site was clipped and cleaned with Nolvasan and ethanol scrubs. A 10-mm midline incision was made over the skull, the skin and fascia were reflected, and a 5-mm craniotomy was made on the central aspect of the left parietal bone. The impounder tip of the injury device was then extended to its full stroke distance (44 mm), positioned to the surface of the exposed dura, and reset to impact the cortical surface. Moderate-level CCI (n=12) was induced using an impactor velocity of 6 m/s and deformation depth of 2 mm as previously described [8, 39–42]. After injury, the incision was closed with interrupted 6-0 silk sutures, anesthesia was terminated, and the animal was placed into a heated cage to maintain normal core temperature for 45 minutes post-injury. Sham animals underwent the same procedure as TBI mice except for the craniotomy and impact. All animals were monitored daily post-injury.

### 2.4 Neurobehavioral testing

#### 2.4.1 Y-maze spontaneous alternation

The Y-maze was carried out at 70 days dpi to access spatial working memory and was essentially performed as previously described [8, 41]. Briefly, the Y-maze (Stoelting Co., Wood Dale, IL) consisted of three identical arms, each arm 35 cm long, 5 cm wide, and 10 cm high, at an angle of 120° with respect to the other arms. One arm was randomly selected as the “start” arm, and the mouse was placed within and allowed to explore the maze freely for 5 mins. Arm entries (arms A–C) were recorded by analyzing mouse activity using ANY-maze software (Stoelting Co., Wood Dale, IL). An arm entry was attributed when all four paws of the mouse entered the arm, and an alternation was designated when the mouse entered three different arms consecutively. The percentage of alternation was calculated as follows: total alternations x 100/ (total arm entries − 2). If a mouse scored >50% alternations (the chance level for choosing the unfamiliar arm), this was indicative of proper spatial working memory.

#### 2.4.2 Novel Object Recognition task

The Novel object recognition (NOR) was carried out to assess non-spatial hippocampal-mediated memory as previously described with slight modifications, on 77-78 dpi [8]. Mice were placed in the NOR chamber where two identical objects were placed near the left and right corners of the open field for training (familiar phase) and allowed to freely explore until they spent a total of 20LJsec exploring the identical objects (exploration was recorded when the front paws or nose contacted the object). After 24 hours, object recognition was tested by substituting a novel object for a familiar training object (the novel object location was counterbalanced across mice). The time spent with each object was recorded using the computed Any-Maze automated software. Because mice inherently prefer to explore novel objects, a preference for the novel object (more time than chance [15LJsec] spent with the novel object) indicates intact memory for the familiar object. Mice that failed to explore and remained immobile throughout the NOR test were excluded from the analysis.

#### 2.4.3 Morris Water Maze (MWM)

Spatial learning and memory were assessed using the Morris Water Maze (MWM) task carried out at 80-84 dpi, as previously described [8]. The MWM protocol included two phases: (1) standard hidden platform training (acquisition) and (2) the twenty-four-hour probe test. Briefly, a circular tank (100 cm in diameter) was filled with water (23±2°C) and was surrounded by various extra-maze cues on the wall of the testing area. A transparent platform (10 cm in diameter) was submerged 0.5 cm below the surface of the water. Starting at 80 dpi, the mice were trained to find the hidden submerged platform located in the northeast (NE) quadrant of the tank for 4 consecutive days (80-83dpi). The mice underwent four trials per day, starting from a randomly selected release point (east, south, west, and north). Each mouse was allowed a maximum of 90LJsec to find the hidden submerged platform. The latency to the platform was recorded by the Any-Maze automated video tracking system. Reference memory was assessed by a probe test carried out at 24 hours following the last acquisition day, on 84 dpi. The platform was removed, and the mice were released from the southwest (SW) position, and the time in the target quadrant was recorded. Additionally, search strategy analysis was performed as previously described [8]. Mice that failed to swim and remained immobile throughout the 90-sec trial in the MWM test were excluded from the analysis.

### 2.5 Isolation of CD11b-Positive Cells

A magnetic bead-conjugated anti-cluster of differentiation 11b (CD11b) was used to isolate microglia/macrophages from ipsilateral (injured hemisphere) cortical and hippocampal tissue using Miltenyi MACS Separation Technology (Miltenyi Biotec, Auburn, CA) as per manufacturers instructions. Briefly, ipsilateral perilesional cortex and hippocampus from Sham and CCI mice were rapidly microdissected and a single cell suspension was prepared from the combined tissues (pooled tissue from two mice) using enzymatic digestion (Neural Tissue Dissociation Kit; Miltenyi Biotec) in combination with a gentleMACS Dissociator. Myelin was removed using Debris Remval Solution step (Miltenyi Biotec) and cells were incubated with anti-CD11b MicroBeads (Miltenyi Biotec) and loaded onto an LS column (Miltenyi Biotec) placed in the magnetic field of a MACS separator. The negative fraction (flow-through) was collected, and the column was washed 3 times with MACS buffer (Miltenyi Biotech). CD11b-positive cells were eluted by removing the magnetic field, resulting in the isolation of viable CD11b-positve cells (microglia/macrophages) from Sham and CCI mice. Cells were snap-frozen on liquid N_2_ for RNA extraction performed using Direct-zol RNA MicroPrep kit (Zymo Research).

### 2.6 Real-Time PCR

Quantitative gene expression analysis in the adipose tissue as well as ipsilateral hippocampus and peri-lesional cortex of Sham and CCI mice was performed using Taqman technology as previously described [8, 41]. Real-time PCR for target mRNAs was performed using TaqMan gene expression assays (NADPH oxidase 2 (NOX2), Mm01287743_m1; human neutrophil cytochrome blight chain (p22^phox^), Mm00514478_m1; Interleukin-1beta (IL-1β), Mm01336189_m1; Tumor necrosis factor-alpha (TNF-a), Mm00443258_m1; Interleukin-10 (IL-10), Mm01288386_m1; NLR family pyrin domain containing 3 (NLRP3), Mm00840904_m1; glial fibrillary acidic protein (GFAP), Mm01253033_m1; Integrin Subunit Alpha M (ITGAM; CD11b). Mm00434455_m1; and GAPDH, Mm99999915_g1; Applied Biosystems, Carlsbad, CA) on a QUANTSTUDIO 5 Real Time PCR machine (Applied Biosystems). Samples were assayed in duplicate in one run (40 cycles), which was composed of 3 stages, 50°C for 2 mins, 95°C for 10 sec for each cycle (denaturation) and finally the transcription step at 60°C for 1 min. Gene expression was normalized by GAPDH and compared to the control sample to determine relative expression values by 2^−ΔΔ*Ct*^ method.

### 2.7 Flow cytometry analysis

Immediately following euthanasia, a 30mg piece of adipose tissue was dissected from each mouse, minced into small pieces and incubated with 150U/ml Collagenase IV (Worthington Biochemical Corporation Lakewood, NJ) and 10 mg/ml DNase II (Sigma) for 1 hour at 37°C in a rotational shaker. The suspension was passed through a 70µm filter to mechanically dissociate adipose tissue. For immune cell surface markers, leukocytes were washed with FACS buffer (5% fetal bovine serum in 1x HBSS) with sodium azide (NaN_3_) and blocked with 1:50 mouse Fc Block (anti-CD16/32) prior to staining with primary antibody-conjugated fluorophores at 1:50 concentration, including CD45-eF450, and CD11b-APCeF780. All antibodies were commercially purchased from Biolegend. For live/dead discrimination, a fixable viability dye, Zombie Aqua (Biolegend), was diluted at 1:100 in Hank’s balanced salt solution (HBSS; Gibco). Cells were briefly fixed in 2 % paraformaldehyde (PFA). Data were acquired on a LSRII using FACSDiva 6.0 (BD Biosciences) and analyzed using FlowJo (Treestar Inc.).

To measure reactive oxygen species (ROS) levels, leukocytes were incubated with dihydrorhodamine (DHR)123 (5mM; 1:500 in RPMI; Ex/Em: 500/536), a cell-permeable fluorogenic probe (Life Technologies/Invitrogen, Waltham, MA). DHR123 passively diffuses into cells and is oxidized by peroxide and peroxynitrite, causing a reaction that produces a green fluorescence that can be measured by flow cytometry. Cells were loaded for 20 minutes in a 37°C water bath, washed twice with FACS buffer (without NaAz), and then stained for surface markers including viability dye, and subsequently fixed in PFA.

### 2.8 Leptin and Insulin determination

At 28 dpi, serum leptin and insulin were measured using the Mouse/Rat Leptin Quantikine ELISA Kit (Cat No. MOB00B, R&D Bio systems, Minneapolis, MN, USA) and the Insulin Mouse ELISA Kit (Cat No. EMINS; Thermo-Fisher, Waltham, MA, USA). The assays were performed according to the manufacturer’s instructions.

### 2.9 Determination of resting blood glucose levels

Prior to euthanasia at 28 dpi, mice were fasted for 12 hours. Using the AlphaTRAK Blood Glucose Monitoring System (Zoetis Inc. Parsippany, NJ, USA) resting blood glucose levels were measured via tail snip samples according to the manufacturer’s instructions.

### 2.10 Nanostring Gene Expression Data Analysis

The raw Nanostring files were analyzed using Rosalind software to generate the normalized gene expression data (used for all later analyses) and log2 fold changes. The average gene expression across groups as well as the expression Z-scores were calculated in Excel. Two-way ANOVA in R was used to study the gene expression variability among groups and identify interaction effects between injury and diet, as well as main effects - genes which are affected only by the injury or only by diet, independent of each other - as previously described [43]. The resulting p values and log2 fold changes were imported into IPA package to analyze the activation of various molecular pathways including Canonical, Upstream regulators (genes, RNAs, and proteins); abbreviated throughout as Upstream regulators – Genes, Upstream regulators (drugs and chemicals); abbreviated throughout as Upstream regulators – Drugs, and Diseases and Bio Functions; abbreviated throughout as Diseases.

Two-way ANOVA in GraphPad Prism was used to determine the significance of differences between cumulative gene expression Z-scores and the pathway activation Z-scores generated by IPA software. PCA analysis of the Rosalind differentially expressed genes was performed in R using the pcaExplorer software package [44]. The principal components were derived from the Z-score values of the differentially expressed subset of genes.

### 2.11 Statistical Analysis

Blinding was achieved by ensuring that the individual who carried out behavioral and stereological analyses were blinded to injury or diet groups. Quantitative data were expressed as mean ± SEM or mean ± STDEV with individual data points as indicated. Normality testing was performed; as datasets met normality requirements (D’Agostino and Pearson omnibus normality test), parametric analyses were used. Statistical analysis was performed using a two-way ANOVA with Tukey *post hoc* tests. When comparisons were made between two conditions, an unpaired Student’s *t* test was performed. Statistical analyses utilized Prism v8 for Windows (GraphPad Software) or in R. Significance level was set at p< 0.05.

The consistent application of two-way ANOVA model permits rigorous determination of the effects of each of the two factors (diet and injury), as indicated by the separate factor significance as well as the direction, and magnitude of their combined effects as evidenced by the interaction and *post-hoc* test significance. A non-significant interaction denotes a combined effect that is simply additive whereas, in a significant interaction, the effects of one of the factors of change depend on the condition of the other factor, and the combined effect reflects either a super additive amplification (synergy) or the opposite.

We used two-way ANOVA significant *post-hoc* p values to select genes for IPA analysis rather that t-test p-value (or multiple comparisons-adjusted p values). Although this approach is more stringent and may exclude some biologically significant changes, it is also more appropriate considering the multiple groups evaluated. Furthermore, the application of adjusted p values is not appropriate here in which we focus on Nanostring panels that include genes selected based on their contribution to similar pathways (e.g., inflammation) and may therefore be modulated in a similar fashion. The analysis of IPA data employed repeated two-way ANOVA to account for changes in specific pathways across groups. In the analyses, we introduce a baseline Sham-SD vs Sham-SD comparison with a z-score “0” across all pathways and replaced any IPA-generated N/A classification (not able to detect activation level) by a score of “0”.

The absence of additive effects can also result in a significant interaction factor and may reflect a “ceiling effect” in which the relatively severe TBI model used in our studies prevents detection of the additive impairments after combined TBI and HFD insults, as was observed for cognitive deficits and lesion volume (an indicator of post-traumatic neurodegeneration, data not shown).

### 2.12 Design limitations

**1)** Diet and injury do not have equivalent positions as diet is initiated before and continues after injury. Thus, it is not possible to distinguish between the pre- and post-injury effects of diet and we can only observe the effects of injury on already established diet effects. Furthermore, HFD and resultant obesity are not equivalent because HFD is maintained throughout the experiment, even after obesity has been established. Therefore, we cannot separate effects of continuous HFD on the VAT (and indirectly on the brain) versus more direct actions on the brain.
**2)** The Nanostring transcriptomic analysis, although including more than 750 genes, is less comprehensive than RNAseq. More importantly, the panels used are highly enriched in inflammatory molecules, and as such it is not possible to apply gene set enrichment analysis, which restricted our IPA work to pathway activation score analysis.
**3)** Sampling limitations include the probable presence of brain-resident macrophages and infiltrating monocytes as a smaller component in the CD11b positively selected fraction, the inclusion of combined injured hippocampus and cortex regions for cell isolation, and the absence of cell-specific isolation (macrophages) in the VAT. These and other limitations related to the presence of heterogeneous cellular populations in samples such as brain flowthrough (neuronal, astrocytic, oligodendrocytes, etc.) and adipose tissue (adipocytes, macrophages, etc.) complicate the analysis and may represent an important confounding factor when interpreting reported molecular/cellular changes in this data set. Thus, it may be challenging to distinguish between the “activation” of cell-specific pathways and the relative increase in the proportion of the host cell compared to the other cell types in the tissue.

## 3. RESULTS

### 3.1 High fat diet (HFD) increases body weight and elevates markers of metabolic function, whereas TBI has only isolated effects

Mice fed a chronic HFD diet starting at 12 weeks prior to TBI/Sham surgery exhibited a time-dependent increase in body weight when compared to mice fed on a SD **(Fig. 1A).** In diet-induced obesity models, mice are considered obese when their body weight is 25 percent greater than that of their counterparts; severe obese models have greater than 50 percent increases in body weight (g) [21]. Beginning at 11 weeks prior to injury, we observed a significant effect of time (F_(3.268, 117.6)_=47.77, p<0.0001; two-way repeated ANOVA; Fig. 1A), treatment (F_(30,36)_=281.5, p<0.0001) and an interaction between time * treatment (F_(30,360)_=8.538, p<0.001) on body weight (g). *Post hoc* analysis revealed a significant effect of HFD on body weight (p<0.05 vs Sham-SD mice; **Fig. 1A**); at 11- (p<0.01), 10- (p<0.001), 9- (p<0.0001), 8- (p<0.0001), 7- (p<0.001), 6- (p<0.001), 5- (p<0.01), 4- (p<0.01), 3- (p<0.01), 2- (p<0.001), and 1- (p<0.001) weeks prior to induction of TBI. After Sham or TBI, mice fed on a HFD showed significant increases in body weight at 0- (p<0.001), 1- (p<0.001), 3- (p=0.000), 7- (p=0.000), 14- (p=0.000), 21- (p=0.000) and 28- (p=0.000) dpi, compared to their Sham-SD and TBI-SD counterparts **(Fig. 1A).** TBI alone did not induce significant changes in body weight (g), nor did it influence HFD-induced increases in body weight, with no significant factor effect, interaction, or difference between Sham-HFD vs. TBI-HFD in *post-hoc* analysis **(Fig. 1A)**. In addition, HFD-induced increases in body weight were independent of total food intake in grams (data not shown). Both diet (F_(1,14)_=284.2, p<0.0001) and TBI (F_(1,14)_=5.175, p<0.05) had significant factor effects inducing significant increases in percent serum leptin at 28 dpi **(Fig. 1B),** but no significant interaction was detected. *Post hoc* analysis revealed that Sham-HFD mice had significantly increased percent serum leptin when compared to Sham-SD counterparts (p<0.0001), at 28 dpi. TBI-SD mice had significantly increased percent leptin when compared to Sham-SD counterparts (p<0.05) although the magnitude of the increase was much smaller compared to HFD groups. While TBI did not further exacerbate the HFD-induced increases in percent serum leptin, TBI-HFD mice displayed significant increases in percent serum leptin when compared to TBI-SD counterparts (p<0.0001), at 28 dpi **(Fig. 1B).**

**Figure 1:**
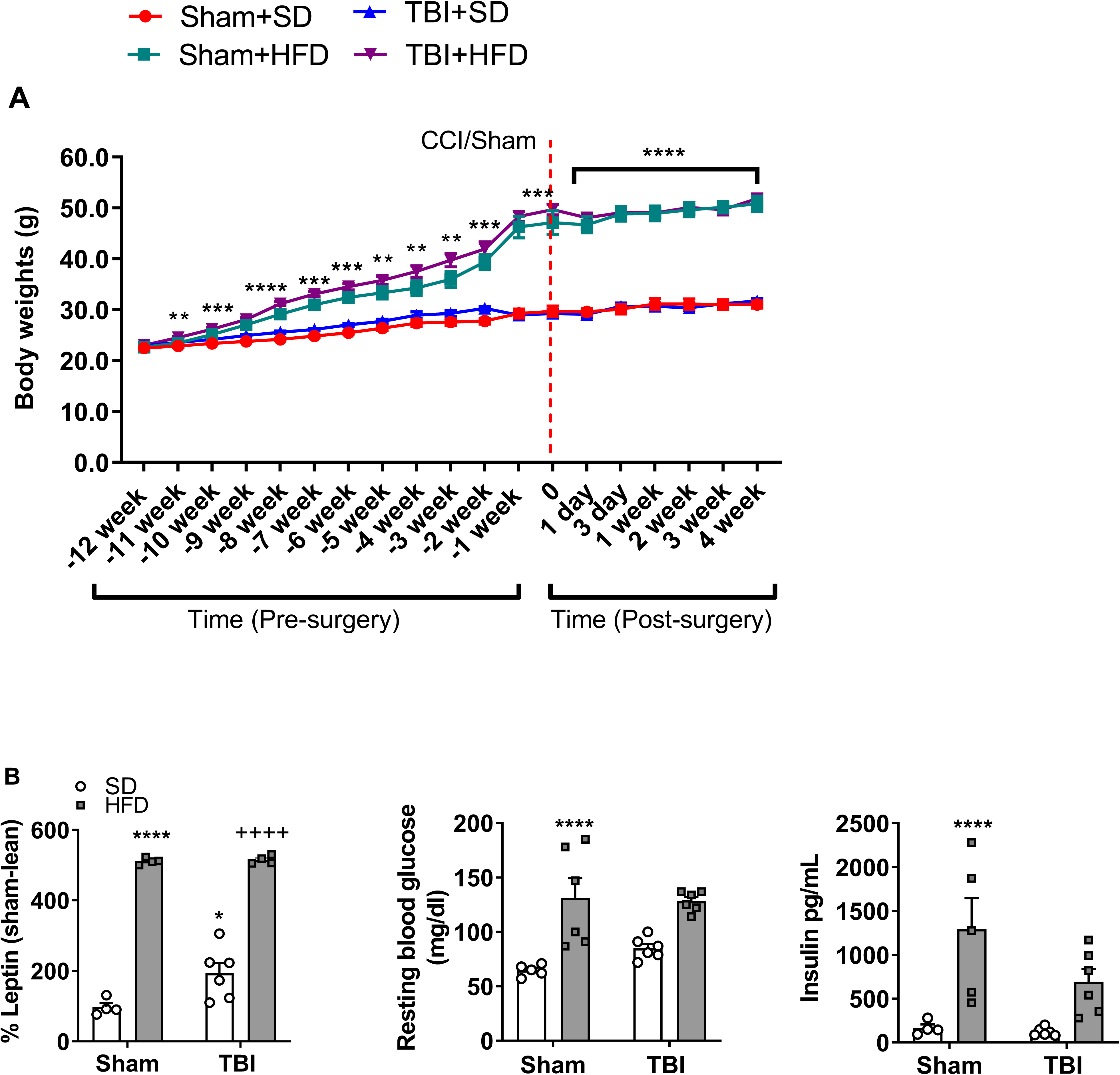
High fat diet (HFD) increases body weight (g) and induces alterations in metabolic responses. **A.** Beginning at 11 weeks prior to surgical intervention, HFD significantly increases body weight (g) through 28 days post-injury (dpi); TBI has no independent effects on body weight (n=8-12 per group). **B.** HFD significantly increases % circulating leptin, and although TBI has a significant yet more modest effect, no further increase is observed in HFD-TBI mice. HFD significantly increases **C.** resting blood glucose and **D.** circulating insulin, while TBI has no independent effects (n=4-6 per group). ** p < 0.01, *** p < 0.001, **** p < 0.0001 vs Sham+SD. ++++ p < 0.0001 vs TBI+SD. Data expressed as mean ± SEM.

Statistical analysis revealed that chronic HFD, but not TBI, resulted in increases in resting glucose (F_(1,19)_=30.02, p<0.0001) and insulin (F_(1,17)_=19.47, p<0.001) compared to their SD counterparts at 28 dpi. *Post hoc* analysis determined that Sham-HFD mice had significantly increased resting blood glucose and insulin when compared to their Sham-SD counterparts (p<0.01; **Fig. 1B)**. No significant interaction between diet and TBI factors was detected.

### 3.2 HFD increases visceral adipose tissue and elevates cellular and molecular markers of inflammation; TBI exacerbates selective HFD effects

Chronic HFD induced significant increases in VAT weight (g/g body weight) (F_(1,18)_=15.57, p<0.001), when compared to Sham-SD mice at 28 dpi; no significant TBI factor effect nor interaction were detected TBI **(Fig. 2A)**.

**Figure 2:**
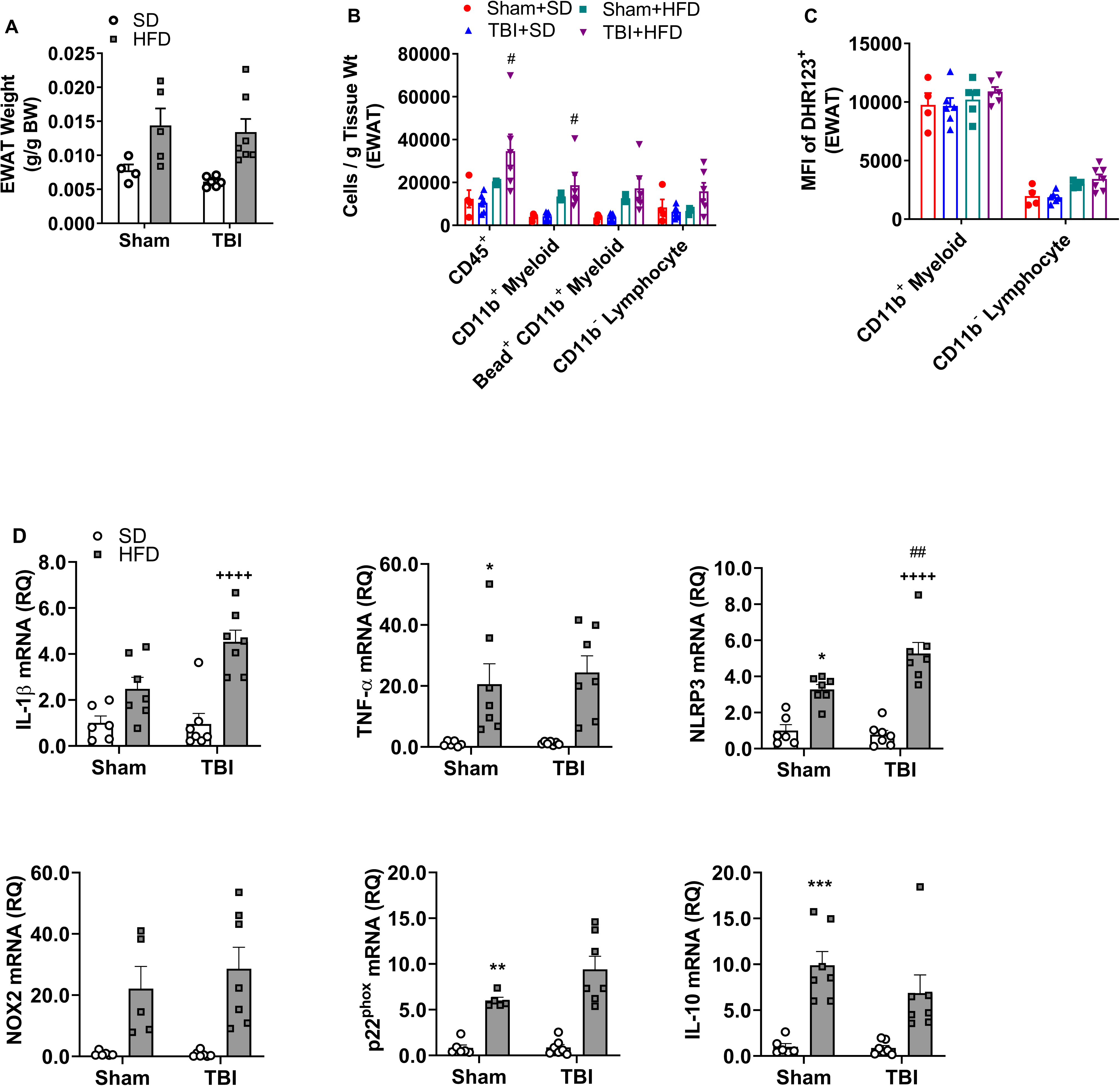
TBI exacerbates HFD-induced increases in VAT inflammatory responses. **A.** The effect of HFD and TBI on VAT weight (g/g BW) at 28 dpi. **B.** TBI-HFD mice display a significant increase in CD45^+^ and CD11b^+^ myeloid cells when compared to Sham-HFD counterparts. **C.** The effect of HFD and TBI on cellular ROS activity. **D.** Sham-HFD mice display a significant increase in VAT expression of *TNF-α*, *NLRP3*, *p22^phox^*, and *IL-10*, when compared to Sham-SD counterparts. TBI-HFD mice exhibit a significant increase in VAT expression of *IL-1β* and *NLRP3*, when compared to TBI-SD counterparts and notably compared to Sham-HFD in the case of NLRP3. * p < 0.05, ** p < 0.01, *** p < 0.001 vs Sham-SD; ++++ p < 0.0001 vs TBI-SD; ## p<0.01 vs Sham-HFD. N=6-7 per group. Data expressed as mean ± SEM.

Flow cytometry was used to characterize the VAT immune cells environment. Diet but not injury had significant factor effects on CD45^+^ (F_(1,15)_=8.838, p<0.01) and CD11b^+^ (F_(1,16)_=14.55, p<0.01) cell populations at 28 dpi; CD11b^-^ infiltrating lymphocytes were not impacted. *Post-hoc* analysis revealed that while HFD alone did not significantly alter lymphocyte cell infiltration, TBI-HFD mice had increased infiltration of CD45^+^ and CD11b^+^ lymphocytes when compared to Sham-HFD counterparts (p<0.05) at 28 dpi **(Fig. 2B)**. Such changes in cell infiltration were not associated with altered intracellular ROS in either CD11b^+^ or CD11b^-^ lymphocytes **(Fig. 2C)**.

To further assess the effect of chronic HFD and TBI on the VAT environment, we also used sensitive qRT-PCR measurement of mRNA expression of several molecular markers of inflammatory responses at 28 dpi. Diet had a significant factor effect on expression of *IL-1β* (F_(1,23)_=29.89, p<0.0001), *TNF-α* (F_(1,23)_=22.57, p<0.0001), *NLRP3* (F_(1,23)_=72.88, p<0.0001), *NOX-2* (F_(1,21)_ 25.08=, p<0.0001), *p22^phox^* (F_(1,21)_=64.21, p<0.0001), and *IL-10* (F_(1,23)_=31.95, p<0.0001) at 28 dpi **(Fig. 2D)**. TBI had a significant effect on the expression of *IL-1β* (F_(1,23)_=4.679, p<0.05) and *NLRP3* (F_(1,23)_=4.925, p<0.05). Furthermore, a significant interaction effect (diet * TBI) was observed on expression of *IL-1β* (F_(1,23)_=5.083, p<0.05), and *NLRP3* (F_(1,23)_=7.799, p<0.05). *Post-hoc* analysis revealed that chronic HFD feeding was associated with significant increases in *TNF-α* (p<0.05), *NLRP3* (p<0.01), *p22phox* (p<0.01), and *IL-10* (p<0.001) when compared to Sham-SD counterparts, at 28 dpi **(Fig. 2D)**. Furthermore, TBI-HFD mice displayed significantly increased expression of *IL-1β* (p<0.0001), and *NLRP3* (p<0.0001) when compared to TBI-SD mice, at 28 dpi, while *NLRP3* was significantly increased in TBI-HFD mice when compared to Sham-HFD mice (p<0.01) **(Fig. 2D)**.

### 3.3 TBI increases neuroinflammation in the injured hippocampus and cortex; HFD exacerbates selective TBI effects

We assessed inflammatory gene changes in key brain regions including the ipsilateral (injured hemisphere) hippocampus, and cortex at 28 dpi. In the hippocampus, we detected a significant factor effect of TBI on *IL-1β* (F_(1,20)_=19.02, p<0.001), *NOX-2* (F_(1,20)_=5.781, p<0.05), *p22^pho^*^x^ (F_(1,20)_=16.71, p<0.001), and *GFAP* (F_(1,20)_=8.408, p<0.01) **(Fig. 3)**. We also observed a significant effect of diet on hippocampal expression of *p22^phox^* (F_(1,20)_=16.71, p<0.001), and *CD11b* (F_(1,20)_=6.993, p<0.05). A significant interaction effect (injury * diet) was noted on hippocampal expression of *p22^phox^* (F_(1,20)_=12.55, p<0.01) at 28 dpi. *Post-hoc* analysis revealed that TBI resulted in a significant increase in hippocampal expression of *IL-1b* (p<0.05), when compared to Sham-SD counterparts **(Fig. 3)**. Although chronic HFD alone (Sham-HFD), did not significantly alter hippocampal expression of any of the inflammatory genes assessed, *post-hoc* analysis detected a significant elevation in *p22^phox^* in TBI-HFD mice when compared to Sham-HFD (p<0.001) and TBI-SD (p<0.001) counterparts (**Fig. 3A)**. In the cortex, there was a significant effect of TBI, but not HFD, on expression of *IL-1b* (F_(1,18)_=11.05, p<0.01), *TNF-a* (F_(1,17)_=30.04, p<0.0001), *NOX-2* (F_(1,17)_=22.74, p<0.001), and *p22^phox^* (F_(1,17)_=32.13, p<0.0001). No significant interaction (TBI * diet) was detected on cortical gene expression at 28 dpi **(Fig. 3B).** *Post-hoc* analysis revealed that TBI resulted in a significant increase in cortical expression of *TNF-α* (p<0.05), *NOX-2* (p<0.05) and *p22^phox^* (p<0.01), but not *IL-1b*, when compared to Sham-SD counterparts **(Fig. 3B)**. These injury effects were not altered by chronic HFD.

**Figure 3:**
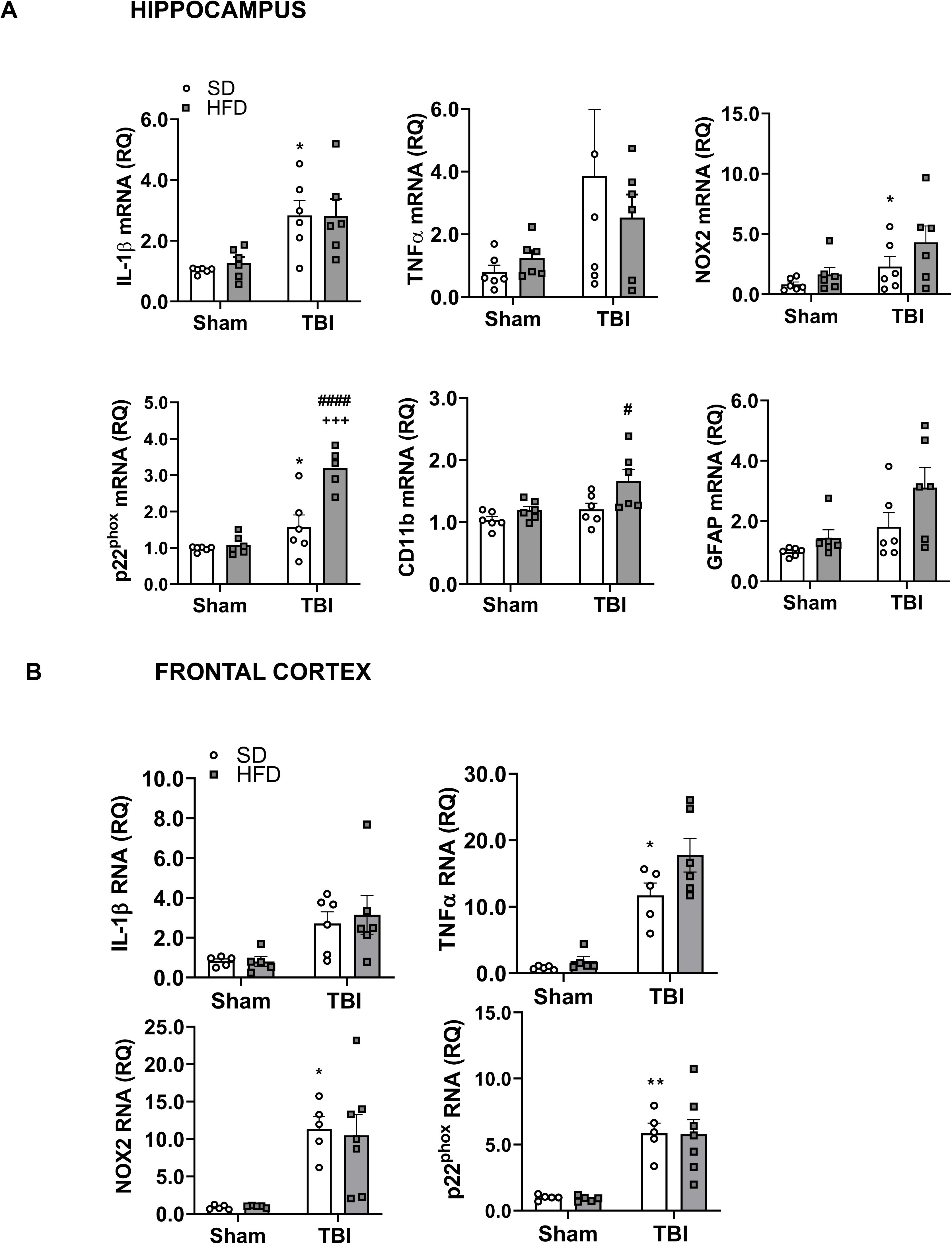
HFD selectively exacerbates TBI-induced increases in the expression of hippocampal, but not cortical neuroinflammatory genes. **A.** TBI-SD mice display a significant increase in hippocampal expression of *IL-1β*, *NOX2*, and *p22^phox^* when compared to Sham-SD counterparts, at 28 dpi. TBI-HFD mice exhibit a significant increase in hippocampal expression of *p22^phox^*when compared to Sham-HFD and TBI-SD. In addition, TBI-HFD displays a significant increase in hippocampal expression of *CD11b*, when compared to Sham-HFD counterparts. **B.** TBI-SD mice display an increase in cortical expression of *TNF-α*, *NOX-2*, and *p22^phox^* when compared to Sham-SD counterparts. * p < 0.05, ** p < 0.01 vs Sham-SD; +++ p < 0.001 vs TBI-SD; ### p < 0.001 vs Sham-HFD. N=6-7 per group. Data expressed as mean ± SEM.

### 3.4 TBI results in chronic memory/learning deficits; HFD exacerbates elements of cognitive dysfunction

To examine the effects of chronic HFD and TBI on long-term neurological outcomes, a second cohort of mice underwent a battery of neurobehavioral tasks probing cognitive functions up to 90 dpi. In the Y-maze, a task assessing hippocampal-dependent working memory, both TBI (F_(1,32)_=9.799, p<0.01) and diet (F_(1,32)_=4.276, p<0.05) factors had significant effects on % spontaneous alterations, but no significant interaction effect (TBI * diet) was observed, at 70 dpi **(Fig. 4A)**. *Post-hoc* analysis determined that TBI-SD mice had significantly less % spontaneous alterations when compared with their Sham-SD mice (p<0.05; **Fig. 4A**). All groups had equivalent numbers of arm entries in the Y-maze task **(Fig. 4A)**.

**Figure 4:**
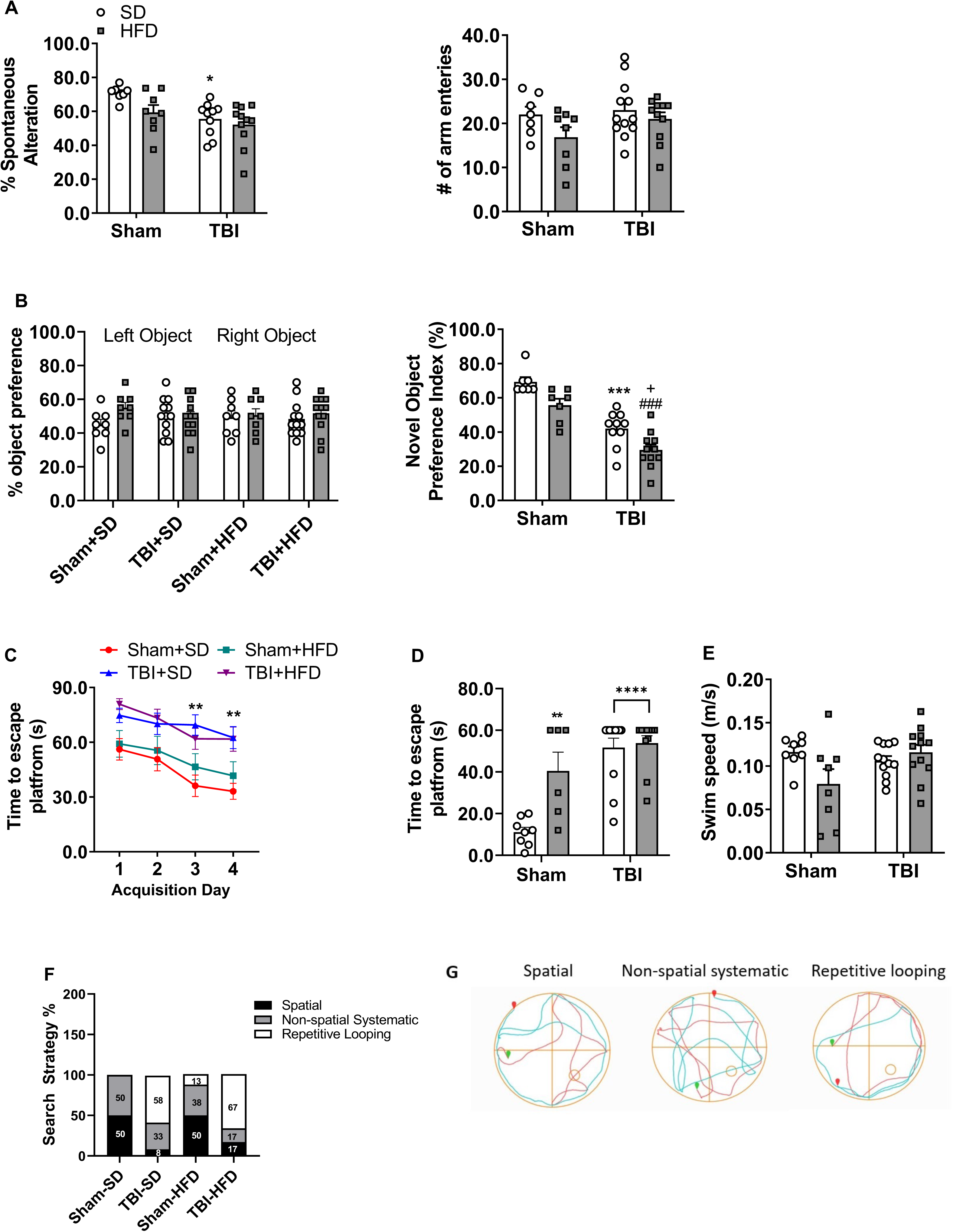
HFD selectively exacerbates TBI-induced deficits in cognitive function. **A.** TBI-SD mice exhibit a significant decrease in % spontaneous alterations in the Y Maze task when compared to Sham-SD counterparts, at 70 dpi; effects of which are independent of number of arm entries. **B.** During the familiar phase of the novel object recognition (NOR), performed at 77 dpi, all four groups spend a similar % time spent with the familiar objects. However, TBI-SD mice spend significantly less % time with the novel object when compared to Sham-SD counterparts, at 78 dpi. Notably, TBI-HFD mice spend significantly less % time with the novel object, when compared to TBI-SD mice. **C.** Compared to Sham-SD mice, TBI-SD mice take significantly longer to find the hidden platform (s) on acquisition day (AD)3 and AD4 of the acquisition phase of the Morris water maze (MWM) task, carried out on 80-83 dpi. **D.** TBI-SD and Sham-HFD mice display a significant increase in time to find the escape platform (s) in the probe trial at 84 dpi, when compared to their Sham-SD counterparts. **E.** The effect of TBI and HFD on swim speed (m/s). **F.** TBI-SD mice utilize an increased % repetitive looping search strategy (58%) when compared to Sham-SD counterparts (0%). Notably, TBI-HFD mice demonstrate an increased % of repetitive looping search strategy (67%) when compared to TBI-SD counterparts (58%). * p < 0.05, ** p < 0.01, *** p < 0.001, **** p < 0.0001 vs Sham-SD; + p < 0.05 vs TBI-SD; #### p < 0.0001 vs Sham-HFD. N=8-12 per group. Data expressed as mean ± SEM.

Non-spatial hippocampal-mediated memory was assessed using the NOR task at 77-78 dpi **(Fig. 4B)**. During the familiar phase of the task, no significant differences were detected between any of the four groups with regards to time spent with either the right or left familiar object. Twenty-four hours later, mice were retested with a novel object and analysis revealed a significant effect of injury (F_(1,31)_=62.41, p<0.001) and diet (F_(1,31)_=14.81, p<0.001) factors on percent time spent with the novel object, but no significant interaction was detected **(Fig. 4B)**. *Post-hoc* analysis showed that, compared with Sham-SD mice, TBI-SD mice spent significantly less time with the novel object (42.00±3.27% versus 69.29±3.27%; p<0.0001). Sham-HFD mice did not spend significantly less time with the novel object compared to the Sham-SD counterparts, but TBI-HFD mice spent significantly less time with the novel object when compared to TBI-SD fed mice (29.55±3.19% versus 42.00±3.27%, p<0.05), indicating that combined pre-existing chronic HFD and TBI further increases TBI-induced deficits in nonspatial hippocampal-mediated memory.

The MWM was designed based on the concept that the hippocampus creates a spatial cognitive map and is responsible for spatial navigation [45]. Spatial learning and memory were examined using the MWM task at 80-84 dpi. During the acquisition trials, two-way repeated ANOVA revealed an effect of time (F_(3,140)_=7.309, p<0.001), treatment (F_(9,140)_=17.94, p<0.001), but no interaction effect between time (acquisition day) * treatment group on latency (s) to the escape platform **(Fig. 4C)**. *Post-hoc* analysis revealed that TBI-SD mice had significantly increased latency to the escape platform when compared to Sham-SD on acquisition day (AD)3 (p<0.01) and AD4 (p=0.01). Twenty-four hours later, retention memory was assessed using the probe trial. There was a significant effect of injury (F_(1,33)_=29.69, p<0.0001) and diet (F_(1,33)_=10.17, p<0.01) factors as well as a significant interaction injury * diet (F_(1,33)_=7.59, p<0.01) on the time to the hidden platform **(Fig. 4D)**. *Post-hoc* analysis revealed that TBI-SD (p<0.0001), Sham-HFD (p<0.01) and TBI-HFD (p<0.0001) had significantly increased latency to the escape platform compared to Sham-SD counterparts. No significant differences were observed between TBI-HFD and TBI-SD **(Fig. 4D)**. There was no significant effect of diet or TBI factors on swim speed **(Fig. 4E)**. The swim pattern used to find the hidden platform was analyzed and assigned a search strategy, classified according to the type of learning taking place, with the least precise/efficient strategies being *repetitive looping*, followed by *non-spatial systematic* and the more precise strategies being *spatial* as described previously [8]. Search strategy analysis permits differentiation between hippocampus-dependent *spatial* (allocentric) and hippocampus-independent *non-spatial* (egocentric) strategies including *systematic* and *repetitive looping* search strategies [46]. A *spatial* pattern indicates an intact hippocampus and adequate spatial memory, whereas *non-spatial systematic* and *repetitive looping* patterns indicate increasing hippocampal dysfunction and growing spatial memory impairments, respectively [46]. Sham-SD mice demonstrated proper memory while increasing memory deficits were noted in TBI-SD, Sham-HFD and TBI-HFD (p<0.05, х =14.60) (**Fig. 4F,G)**.

### 3.5 TBI causes brain microglia transcriptome changes driving increased reactive states and decreased homeostatic phenotype; HFD amplifies posttraumatic activation of inflammation pathways

To identify microglia transcriptional changes, we performed CD11b-positive cells selection (using Miltenyi MACS technology; microglia are the principal component of this cellular fraction) from the cortex and hippocampus in the second cohort at 90 dpi followed by high-throughput gene expression examination using Nanostring panels. RNA was extracted and analyzed using the nCounter Mouse Glial panel (Nanostring Technologies) that profiles more than 750 genes related to glia functions and metabolism. Gene expression data was normalized using the Rosalind Nanostring analysis package and all further analyses were limited to genes whose average expression across all experimental groups was at least 50, to reduce the impact of large variations common in low expressor genes and increase biological significance. In the microglia fraction, 526 genes out of a total of 757 genes met the expression requirement and underwent a further Z-score normalization that permits examination of expression changes independently of absolute expression values. The Z-score gene list was analyzed using Morpheus, an unsupervised hierarchical clustering package (https://software.broadinstitute.org/morpheus). The generated heat map shows that while TBI is the primary determining factor for gene expression changes with HFD having a lesser impact; a consistent gradient of changes is observed with the following group order: Sham-SD, Sham-HFD, TBI-SD, TBI-HFD **(Fig. 5A)**. The gene function description is based on Nanostring annotations that were updated based on recent data. The general heat map **(Fig. 5A)** and the microglia phenotype-specific heat maps **(Fig. 5B)** identifies homeostatic microglia markers along with other microglia markers as among the most consistently downregulated phenotypes while the disease-associated microglia (DAM) and ‘M1’-like micorglia which are associated with the classical pro-inflammatory markers (pro-inflammatory-associated microglia) [47], were among the most consistently upregulated phenotypes. Markers of autophagy-phagocytosis and interferon signaling were also elevated **(Fig. 5A-B)**.

**Figure 5:**
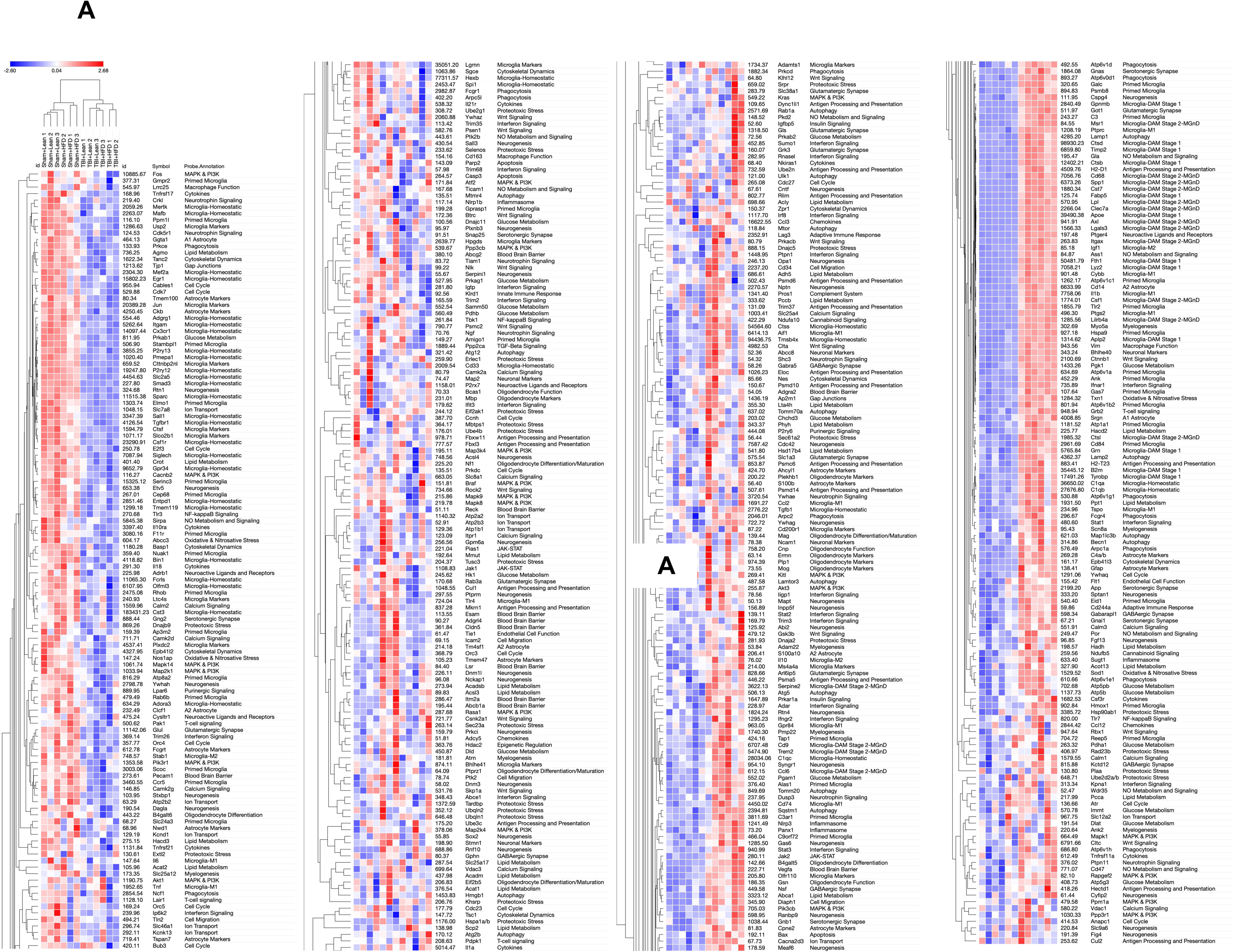
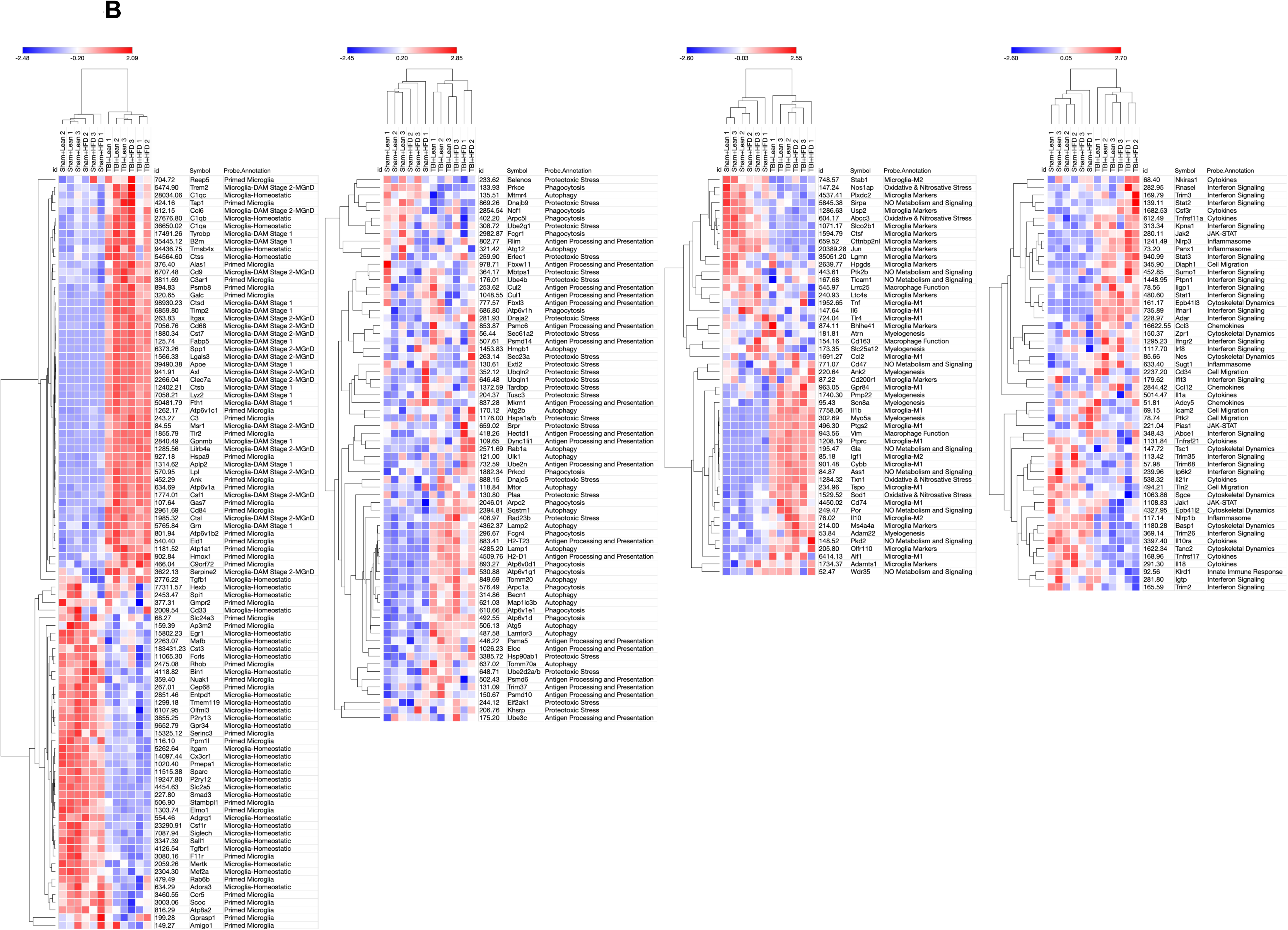
TBI and HFD selectively interact to amplify brain microglia transcriptome changes that indicate the activation of inflammation pathways. **A.** Analysis on isolated CD11b^+^ cells (microglia) ran on a nCounter Mouse Glial Panel (Nanostring Technologies; >750 genes related to glial functions and metabolism) using a general heat map generated by the Morpheus unsupervised hierarchical clustering reveals that TBI is the primary driver of changes in gene expression; HFD has lesser yet detectable effects and a consistent gradient of changes is observed with the following group order: Sham-SD, Sham-HFD, TBI-SD, TBI-HFD. **B.** Microglia-phenotype specific heat maps identify that TBI-SD mice display a consistent downregulation of genes associated with the homeostatic microglia and microglia markers phenotype, when compared to Sham-SD counterparts, at 90 dpi. In contrast, TBI-SD mice display an upregulation of genes associated with disease-associated microglia (DAM), pro-inflammatory-associated, and autophagy microglial phenotypes, when compared to Sham-SD counterparts. Notably, HFD consistently amplifies these effects. N=3 pooled samples/group.

To quantify the gene expression changes reflected by the heatmaps we summated in each sample the Z-scores that corresponded to genes defining specific cellular phenotypes. The generated Z-score sum is an overall indicator of the relative change in expression of the phenotype-specific genes in each sample and the values across groups were analyzed using two-way ANOVA **(Fig. 6)**. Statistical analysis revealed a significant effect of injury on homeostatic-associated microglia (F_(1,8)_= 39.63, p<0.0001), autophagic-associated microglia (F_(1,8)_= 78.60, p<0.0001), interferon-associated microglia (F_(1,8)_= 11.86, p<0.01), pro-inflammatory-associated microglia (F_(1,8)_= 27.90, p<0.001), DAM (F_(1,8)_= 225.4, p<0.0001) and microglia markers (F_(1,8)_= 34.43, p<0.001) **(Fig. 6**; **Fig. 14A-supplemental)**. No significant effect of diet or diet*injury interaction was observed, except a modest diet effect in microglia markers. *Post-hoc* analysis revealed that TBI-SD mice had a significant decrease in homeostatic-associated microglia (p<0.01) and microglial markers (p<0.01) while displaying a significant increase in autophagic-associated microglia (p<0.01), pro-inflammatory-associated microglia (p<0.01), and DAM (p<0.0001), when compared to Sham-SD counterparts. TBI-HFD showed similar changes compared to Sham-HFD. No significant changes were observed in Sham-HFD vs Sham-SD or TBI-HFD vs TBI-SD confirming that the *injury (TBI) factor* but not the *diet (HFD) factor* was associated with significant F ratios and p value in *post-hoc* comparisons **(Fig. 6).**

**Figure 6:**
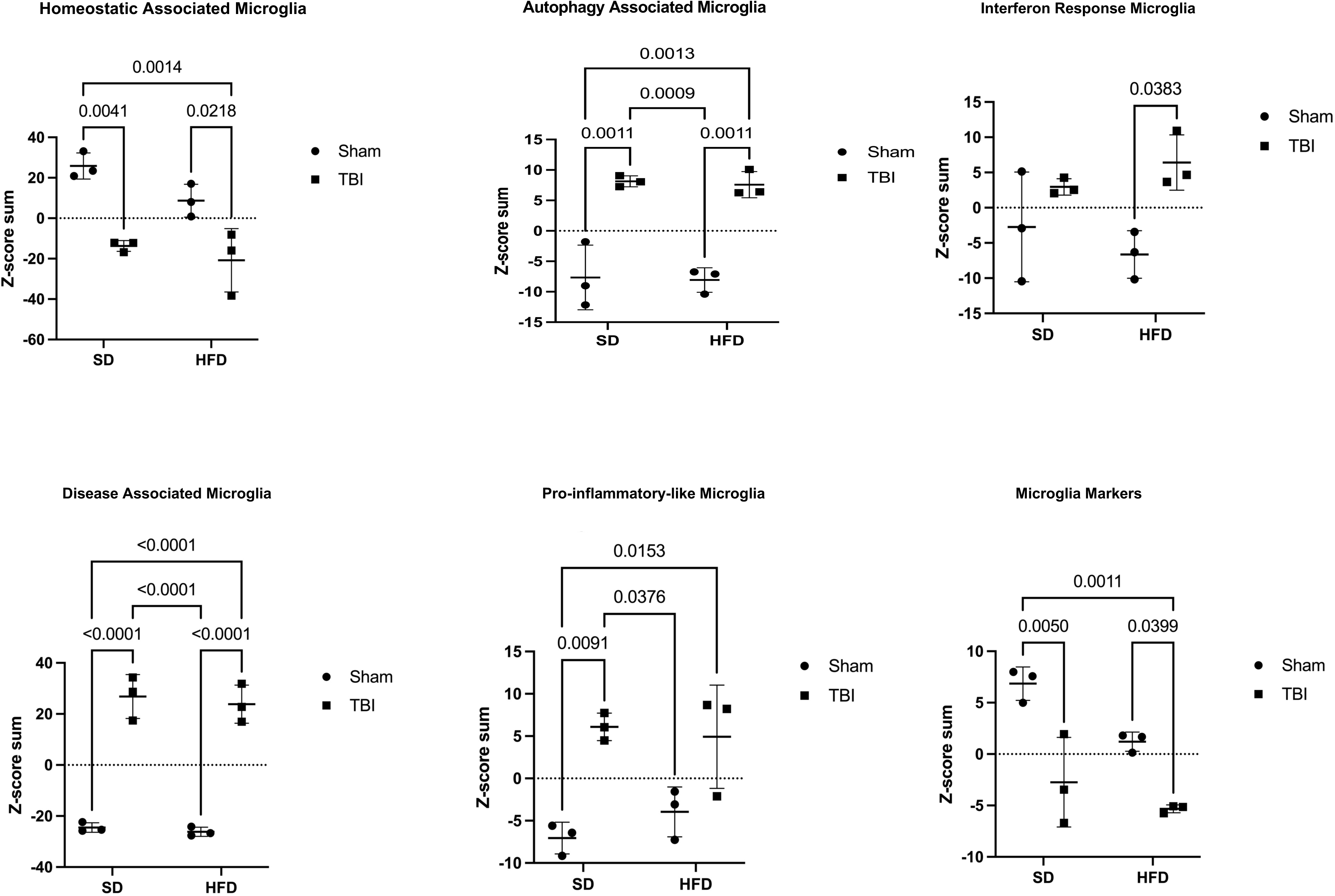
TBI is the main driver of changes in gene expression associated with specific microglial phenotypes. Z-score sum normalization reveals that TBI-SD mice displayed a significant increase in genes associated with the DAM, pro-inflammatory-associated, and autophagy phenotype, when compared to Sham-SD counterparts, at 90 dpi. In contrast, TBI-SD mice had a significant decrease in genes associated with homeostatic and microglia marker phenotypes. Genes associated with interferon-and autophagy-associated microglia, and DAM are significantly upregulated in TBI-HFD when compared to Sham-HFD counterparts. Genes associated with homeostatic and microglia markers are decreased in TBI-HFD when compared to Sham-HFD counterparts. N=3 pooled samples/group. Data expressed as mean ± STDEV.

We also analyzed the normalized expression levels (all genes with average expression >50) using a two-way ANOVA. Genes that demonstrated a significant p value (p≤0.05) in at least one factor, interaction or *post-hoc* test were selected for further analysis (n=312 genes). The Morpheus-generated unsupervised hierarchical clustering heat map **(Fig. 1-supplemental;** annotations include all ANOVA outcomes) demonstrates the same patterns as in **Fig. 5A**, although gene expression patterns are easier to detect due to the elimination of genes that do not significantly change with injury/diet.

Ingenuity Pathway Analysis (IPA) software (QIAGEN) is an advanced bioinformatics tool, based on built-in scientific literature databases (www.ingenuity.com) that identifies the most impacted molecular pathways and predicts cellular responses. For the 312 genes with significant ANOVA results we imported into IPA the Rosalind-generated log2FC of Sham-HFD vs Sham-SD, TBI-SD- vs Sham-SD and TBI-HFD vs Sham-SD and the corresponding *post-hoc* p values generated by the repeated two-way ANOVA analysis. We determined the activation of IPA canonical pathways based on the inclusion of genes with p≤0.05 and pathway z-score >2 or <-2 (the minimal requirements for IPA to conclude that a pathway is activated or inhibited, respectively) **(Fig. 7)**. The data suggest that TBI-HFD tends to cause more pathway changes including stronger activation of multiple inflammatory pathways. We also quantified these changes after separating the activated and inhibited pathways (based on the outcome in the TBI-HFD group) and analyzing the pathway activity z-scores with repeated two-way ANOVA. To assist analysis, we introduced a baseline Sham-SD vs Sham-SD comparison with a z-score “0” across all pathways and replaced any IPA-generated N/A classification (not able to detect activation level) with a score of “0”. Statistical analysis shows injury (F_(1,16)_= 199.8, p<0.0001), diet (F_(1,16)_= 10.29, p<0.01), and an injury*diet interaction (F_(1,16)_= 10.29, p<0.01) associated with activation/upregulation of a group of Canonical pathways mostly composed by pro-inflammatory pathways **(Fig. 7**; **Fig. 14A-supplemental)**. For the less numerous Canonical pathways that were downregulated/inhibited, significance was only observed for an injury effect (F_(1,4)_= 458.9, p<0.0001). *Post hoc* analysis revealed that TBI-SD mice had significantly increased Z-score (p<0.0001) vs Sham-SD mice. TBI-HFD showed similar changes when compared to Sham-HFD. Sham-HFD was not associated with pathway activation or inhibition. Notably, TBI-HFD mice displayed significantly increased Z-scores (p=0.0017) vs TBI-SD, for canonical upregulated pathways (**Fig. 7**). For the downregulated pathways, *post hoc* analysis revealed that TBI-SD mice had a significantly decreased Z-score (p<0.0001) vs Sham-SD and that TBI-HFD showed similar changes when compared to Sham-HFD.

**Figure 7:**
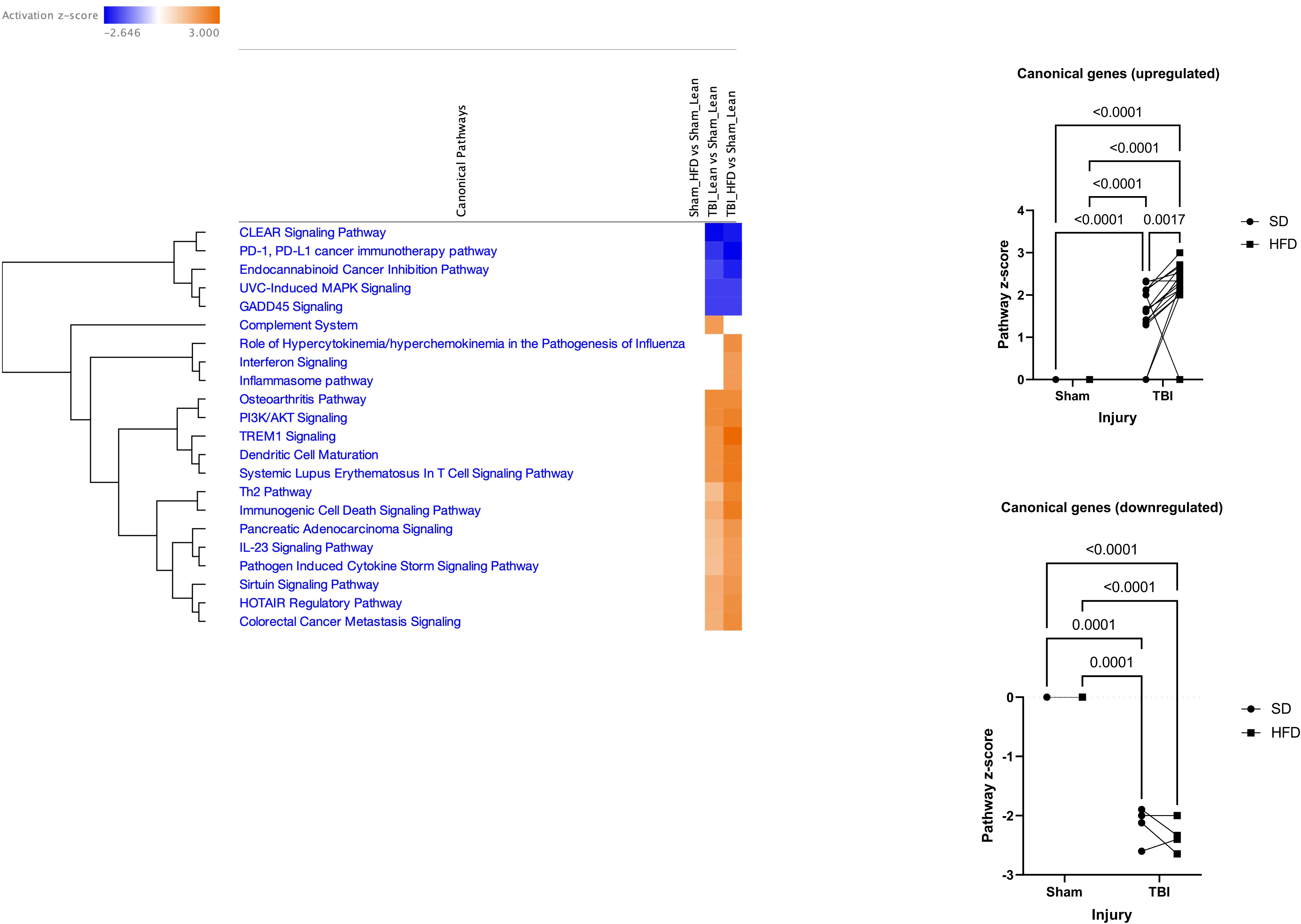
Ingenuity Pathway Analysis (IPA) shows that while TBI is the principal driver of pathway changes, HFD has a significant effect to amplify the activation of multiple inflammatory pathways. IPA determined the activation of its set of canonical pathways based on the inclusion of genes with p<0.05; the pathways’ z-score is an indicator of activity level (positive-upregulated; negative-downregulated); all pathways with z-score >2 or <-2 in at least one group are presented as these levels are considered the minimal criteria for significant upregulation or downregulation, respectively. The microglia from TBI-SD mice demonstrated significant upregulation and downregulation in specific canonical pathways when compared to Sham-SD. Notably, TBI-HFD mice displayed a significant increase in the activation level of canonical upregulated pathways, when compared to TBI-SD counterparts. Data expressed as mean ± STDEV.

We also examined changes in IPA Upstream regulators - Genes (**Fig. 2-supplemental**), Upstream regulators - Drugs (**Fig. 3-supplemental**) and Diseases (**Fig. 4-supplemental**). The injury factor and related *post-hoc* tests were significant in all cases, but significant diet factor was only observed in Upstream Genes and Drugs and significant interaction effects with significant TBI-HFD vs TBI-SD *post-hoc* test was only observed in Upstream Drugs upregulated pathways which included several inflammatory pathways (e.g., lipopolysaccharide (LPS), polyinosinic:polycytidylic (polyI-C), etc.)).

### 3.7 TBI is the primary factor driving changes in non-microglia cellular transcriptomes

To determine the specificity of the CD11b-positive selection and the resulting selective separation of microglia and CD11b-negative (non-microglia) populations (neurons, astrocytes, oligodendrocytes, etc), we averaged the gene expression levels across microglia and non-microglia fraction in each experimental group followed by Z-score transformation. The ratio between averaged microglia and flowthrough normalized expression data (560 genes out of a total of 769 genes had average expression >50 in the flowthrough) was used to determine the top 50 overexpressed and under expressed genes in microglia vs flowthrough. The Z-scores for these genes were uploaded into Morpheus for unsupervised hierarchical clustering **(Fig. 5- supplemental)**. The data confirm the quality of the separation as the CD11b-positive fraction demonstrates over hundredfold enrichment of key microglia markers. The reverse is true for the flowthrough in which neuronal, astrocytes and oligodendrocytes markers are highly enriched. Notably, even at this reduced scale the gradient of injury/diet changes is maintained and a decline in the microglia homeostatic phenotype and neurogenesis as well as increase in microglia and astrocyte altered phenotypes after injury/HFD are observed in the appropriate compartments.

Performing the Morpheus heat map on all 560 genes in the flow-through did not result in an ordered clustering across the groups highlighting the smaller number of genes with consistent and significant changes **(Fig. 6-supplemental)**. Conversely, when a further selection restricted the list to genes demonstrating a p≤0.05 in at least one of the two-way ANOVA outcomes (as described above; n=121 genes; **Fig. 8**) the resulting heat map displayed a consistent gradient of changes across groups that matched the clustering observed in the microglia (Sham-SD, Sham-HFD, TBI-SD, TBI-HFD). Genes downregulated included neurogenesis and neuronal signaling markers while genes upregulated included astrocyte activation phenotype markers **(Fig. 8**; **Fig.7- supplemental** includes the heatmap with ANOVA data). The generated Z-score sum demonstrates that injury is the only factor driving significant downregulation of neuronal markers (F_(1,8)_= 30.95, p<0.001), an upregulation of astrocytes (F_(1,8)_= 18.79, p<0.01) and oligodendrocytes markers (F_(1,8)_= 23.60, p<0.01); injury*diet interaction was significant for neuronal (F_(1,8)_= 8.78, p<0.05) and oligodendrocyte (F_(1,8)_= 5.41, p<0.05) markers but it was indicative of absence of additive effects as *post-hoc* statistical analysis revealed that TBI-SD (p < 0.01) and TBI-HFD (p < 0.01) had similar effects vs. Sham-SD while diet alone (Sham-HFD) had effects in the same direction as injury although these reached significance only in the case of neuronal markers (**Fig. 8**; **Fig. 14B-supplemental**). These changes may in part reflect declines in neuronal populations and increases in astrocytic and oligodendrocytes populations. The IPA examination of molecular responses failed to detect any Canonical pathways with significant changes (z>2 or z<-2). The few activated Upstream genes pathways showed significance only for the TBI factor and no significant interaction **(Fig. 8-supplemental)**. The Diseases pathways displayed multiple inhibited or activated genes; the majority of the activated pathways were inflammatory with only TBI as a significant factor and a modest but significant increase in this response was observed in the TBI-HFD group vs. TBI-SD (**Fig. 9-supplemental**).

**Figure 8:**
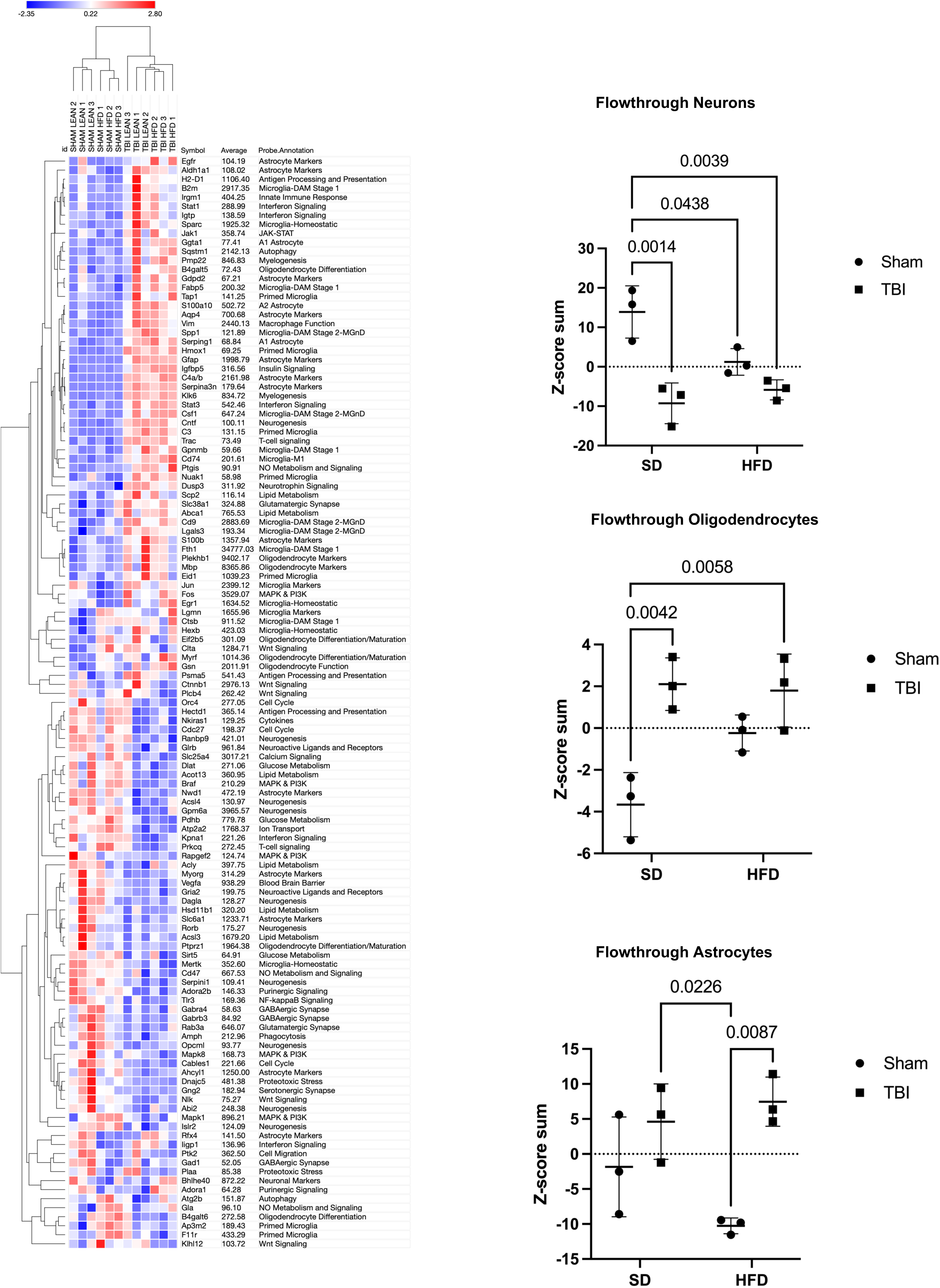
TBI is the primary factor of change in the flowthrough cellular transcriptome. nCounter Mouse Glial Panel analysis was performed in the isolated CD11b^-^ populations (neurons, astrocytes, oligodendrocytes, etc). Only the genes that showed evidence of significant changes (p<0.05 in at least one of the two-way ANOVA outcomes; n=121 genes) were included and the resulting heat map displayed a consistent gradient of changes across groups that matched the clustering previously observed in the microglia (Sham-SD, Sham-HFD, TBI-SD, TBI-HFD). TBI-SD mice displayed a significant decrease in neuronal genes, with a concurrent increase in genes associated with oligodendrocyte responses, when compared to Sham-SD counterparts. N=3 pooled samples/group. Data expressed as mean ± STDEV.

### 3.8 HFD and TBI interact to amplify VAT transcriptome changes that are characteristic of macrophage reactive states and inflammation pathways

We extracted the RNA from VAT in the experimental animals in the second cohort at 90 dpi and examined the gene expression changes using the Nanostring Neuroinflammation panel. The Nanostring data were analyzed as described above detecting 483 genes out of a total of 777 with average expression >50. The general heat map generated by the Morpheus unsupervised hierarchical clustering displays a consistent order in the gradient of differential expression (Sham-SD, TBI-SD, Sham-HFD, TBI-HFD) which in contrast to the brain changes, has diet as the primary factor. **(Fig. 9A)**. The cell-phenotype-directed heatmaps highlight the abundance of changes impacting the VAT immune/inflammation responses, representing the majority component among activated genes, that maintain the same clustering order **(Fig. 9B)**. The general heat map based only on the 369 genes with significant ANOVA outcomes display a similar order while bringing into sharper distinction the upregulated inflammation markers **(Fig. 10-supplemental)**. Statistical analysis reveals a significant effect of diet on the Z-score sum on adipose macrophage homeostatic (F_(1,8)_= 56.16, p<0.0001), adipose macrophage DAM (F_(1,8)_= 39.55, p<0.001), adipose tissue function (F_(1,8)_= 35.34, p<0.001), adipose inflammatory signaling (F_(1,8)_= 20.38, p<0.01), and macrophage pro-inflammatory (F_(1,8)_= 85.04, p<0.0001) but not adipose innate immunity (**Fig. 10**; **Fig. 14C-supplemental**). There was a significant injury effect reported for adipose macrophage function (F_(1,8)_= 11.30, p<0.01), adipose innate immunity (F_(1,8)_= 14.57, p<0.01) and macrophage pro-inflammatory (F_(1,8)_= 7.55, p<0.05). Furthermore, an interaction effect of diet*injury was reported for adipose macrophage homeostatic (F_(1,8)_= 7.12, p<0.05) and adipose innate immunity (F_(1,8)_= 5.81, p<0.05; **Fig. 14C-supplemental)**. *Post hoc* analysis revealed a significant increase in the Z-score sum of adipose macrophage homeostatic and adipose macrophage DAM (p<0.05) as well as pro-inflammatory associated macrophage phenotype (p<0.01) of Sham-HFD vs Sham-SD diet (**Fig. 10**). Notably, TBI-HFD mice displayed a significant increase in adipose macrophage homeostatic (p<0.05), adipose macrophage function (p<0.05), and adipose innate immunity (p<0.01) vs Sham-HFD mice. The pathways-centered analysis in IPA demonstrates that while diet (F_(1,52)_= 1293, p<0.001) is the dominant factor, the injury (F_(1,52)_= 119.2, p<0.0001) factor also reaches significance and that a significant diet*injury interaction (F_(1,52)_= 90.09, p<0.0001) is associated with the robust superadditive activation of multiple Canonical pathways(majority are inflammation-related), by combined TBI-HFD (p<0.0001) versus Sham-HFD **(Fig. 11**; **Fig. 14C-supplemental)**. The Upstream regulators - Genes **(Fig. 11-supplemental)**, Upstream regulators - Drugs **(Fig. 12- supplemental)** and Disease pathways **(Fig. 13-supplemental)** also display consistent HFD-driven activation as a principal factor accompanied by injury factor significance and *post-hoc* tests as well as significant diet*injury interaction associated with amplification of both activated (the majority are inflammation-related) and inhibited pathways by combined TBI-HFD, including upstream genes upregulated (p<0.0001), upregulated (p<0.0001) and downregulated (p<0.0001) upstream drugs, and diseases upregulated and downregulated (p<0.0001), when compared to Sham-HFD.

**Figure 9:**
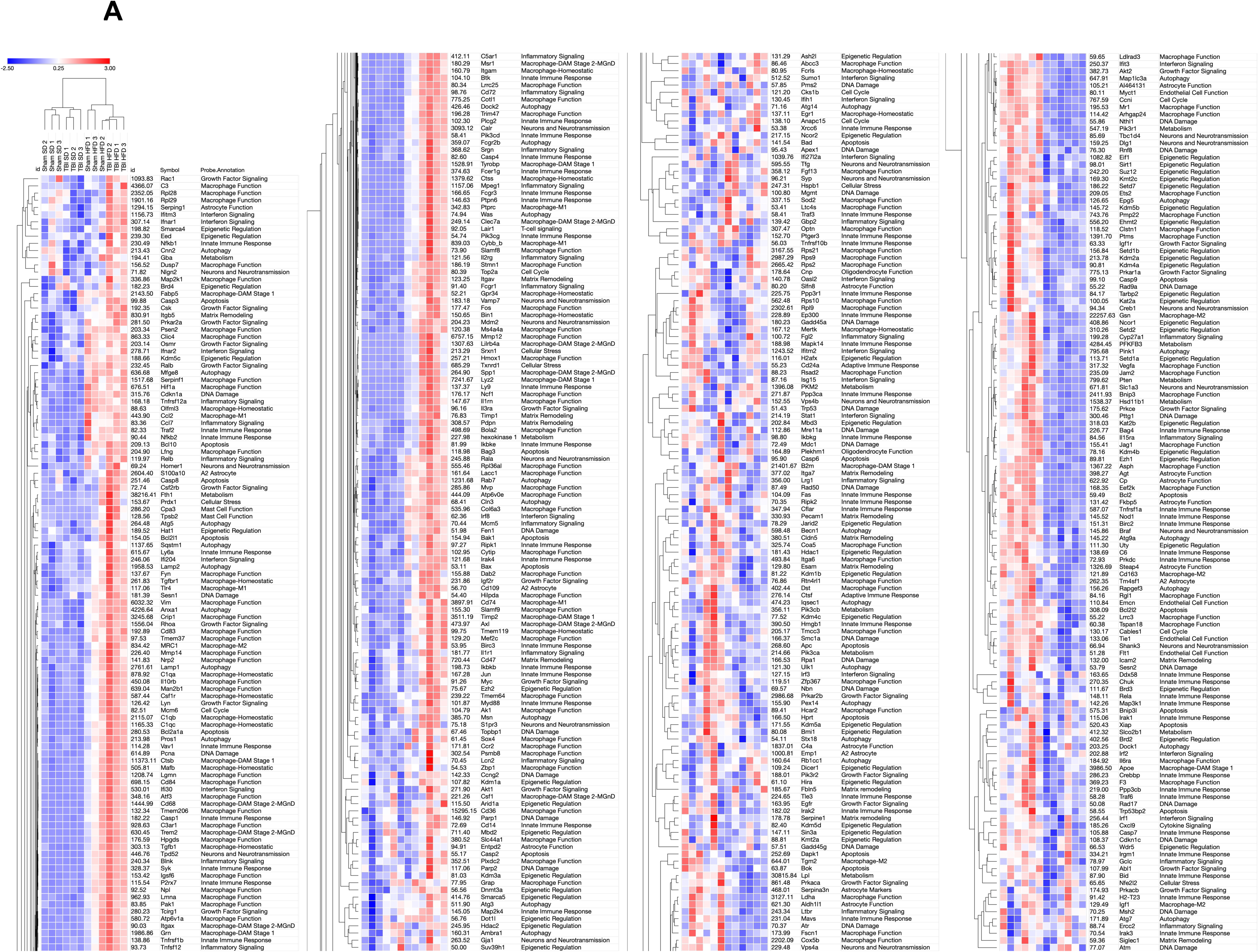
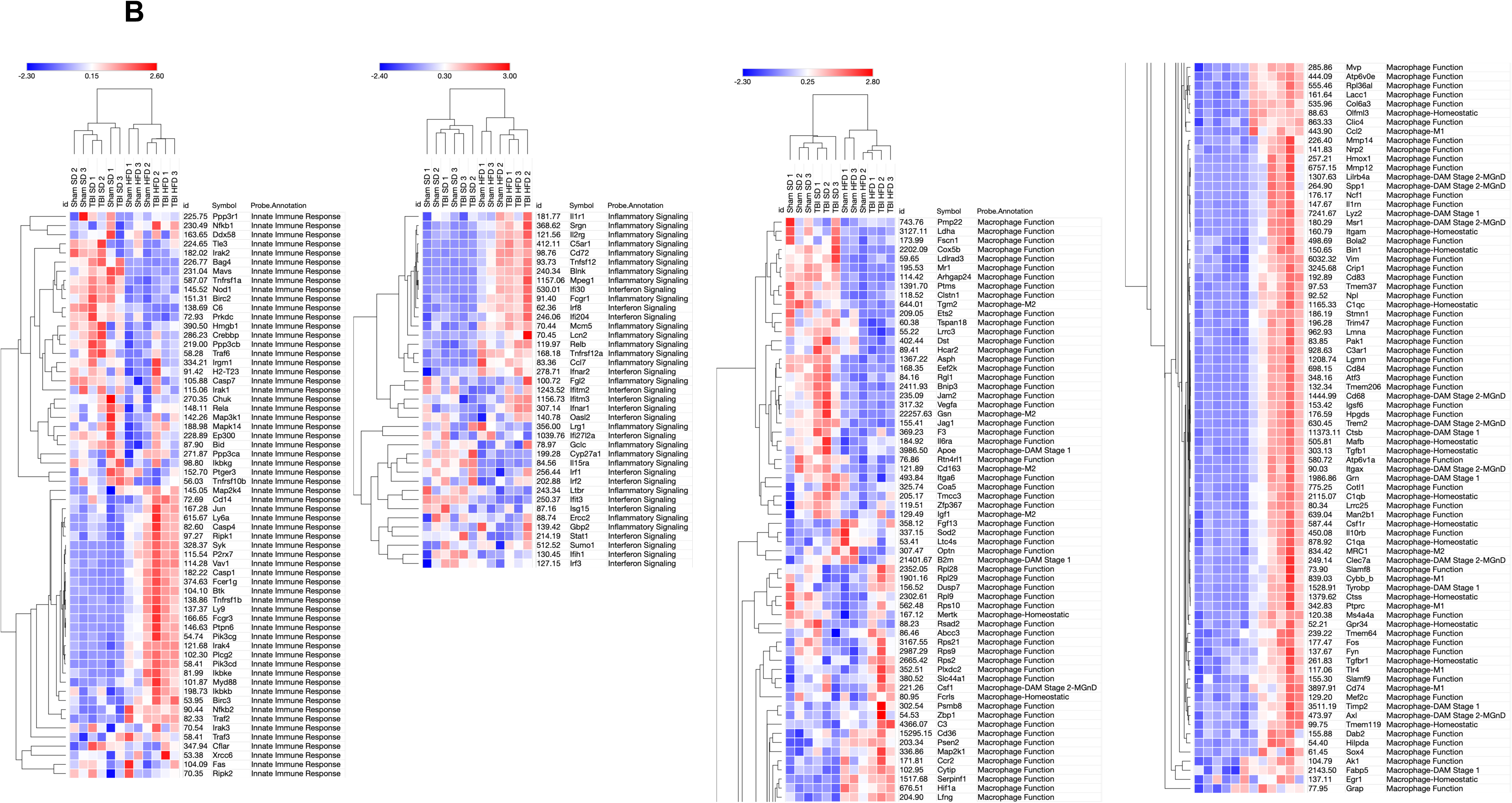
HFD and TBI interact to amplify VAT transcriptome changes that indicate the activation of inflammation pathways. **A.** Analysis on isolated VAT ran on a nCounter Mouse Neuroinflammation Panel (Nanostring Technologies; >750 genes related to immunity & inflammation). The general heat map generated by the Morpheus unsupervised hierarchical clustering displays a consistent order in the gradient of differential expression (Sham-SD, TBI-SD, Sham-HFD, TBI-HFD); in contrast to the brain changes, diet and not TBI is the primary factor. **B.** More cell-phenotype-directed heat maps highlight that Sham-HFD mice display increases in genes associated with homeostatic, DAM, and pro-inflammatory-associated adipose macrophage phenotype when compared to Sham-SD counterparts. TBI-HFD mice displayed increases in homeostatic, adipose macrophage function, and adipose innate immunity phenotype when compared to Sham-HFD counterparts. N=3 pooled samples/group.

**Figure 10:**
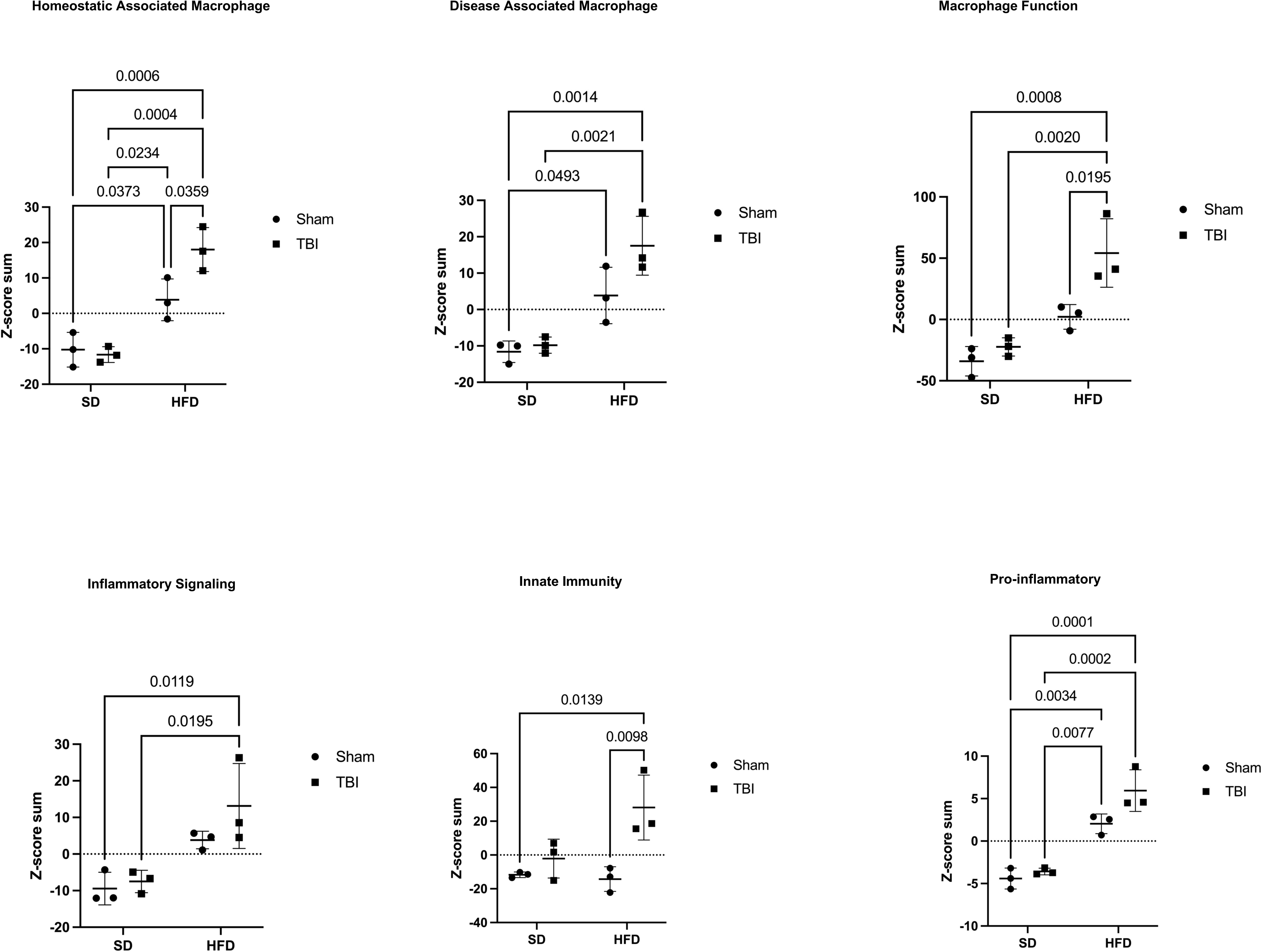
HFD and TBI interact to amplify VAT transcriptome changes associated with specific macrophage activated inflammatory states. Z-score sum normalization reveals that Sham-HFD mice had a significant increase in genes associated with adipose macrophage homeostatic, adipose macrophage DAM, and adipose macrophage pro-inflammatory phenotype when compared to Sham-SD counterparts. Furthermore, TBI-HFD mice displayed a significant increase in adipose macrophage homeostatic, adipose macrophage function, and adipose innate immunity vs Sham-HFD mice. Notably, TBI-HFD mice had a significant increase in genes associated with adipose macrophage homeostatic, adipose macrophage function, and adipose macrophage innate immunity, when compared to Sham-HFD counterparts. N=3 pooled samples/group. Data expressed as mean ± STDEV.

**Figure 11:**
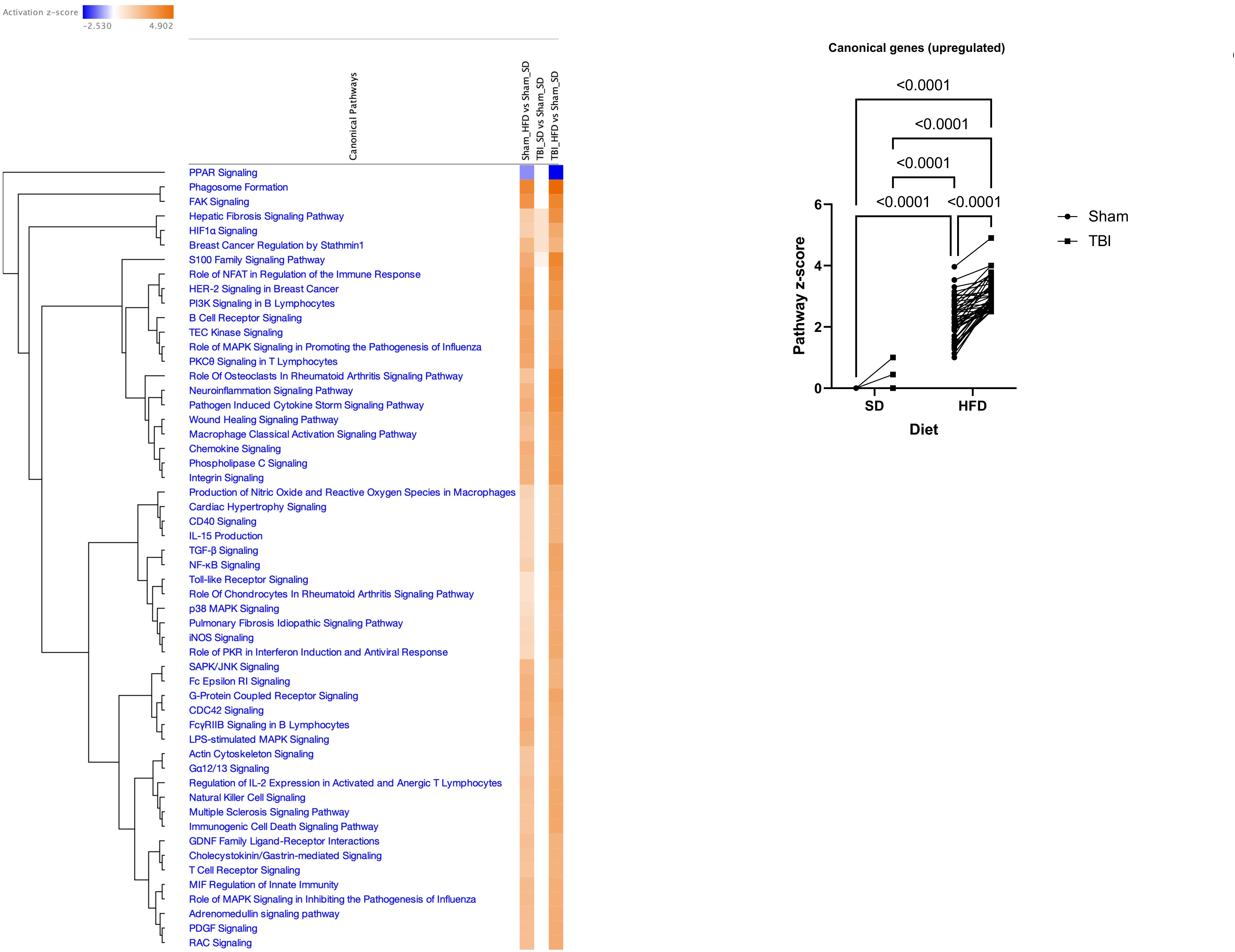
IPA indicates an interaction between HFD and TBI to amplify the activation of multiple inflammatory pathways in VAT. Sham-HFD mice display a significant increase in activation of multiple canonical upregulated pathways, compared to Sham-SD mice. Notably, TBI-HFD mice demonstrated a significant increase in the activation of multiple canonical upregulated pathways, compared to Sham-HFD mice.

### 3.9 HFD and TBI contribute to global separation of cellular phenotypes

We used principal component analysis, a linear dimensionality reduction technique to highlight the transcriptomic differences between specific groups. The gene expression changes for every cell fraction, based the Z-scores of genes that have been identified as differentially expressed in Rosalind, were analyzed by PCA. Consistent with previous data, the microglia and flowthrough show TBI as a primary factor for changes with HFD playing a secondary role, whereas the reverse was true for the VAT. Notably, when PCA was applied to an integrated data set (brain microglia and flowthrough as well as VAT) for every experimental animal, we generated a composite score that resulted in robust separation of all groups with combined TBI-HFD located at the union of individual factors (TBI and HFD) directions of change and most distinct from Sham-SD (**Fig. 12**).

**Figure 12:**
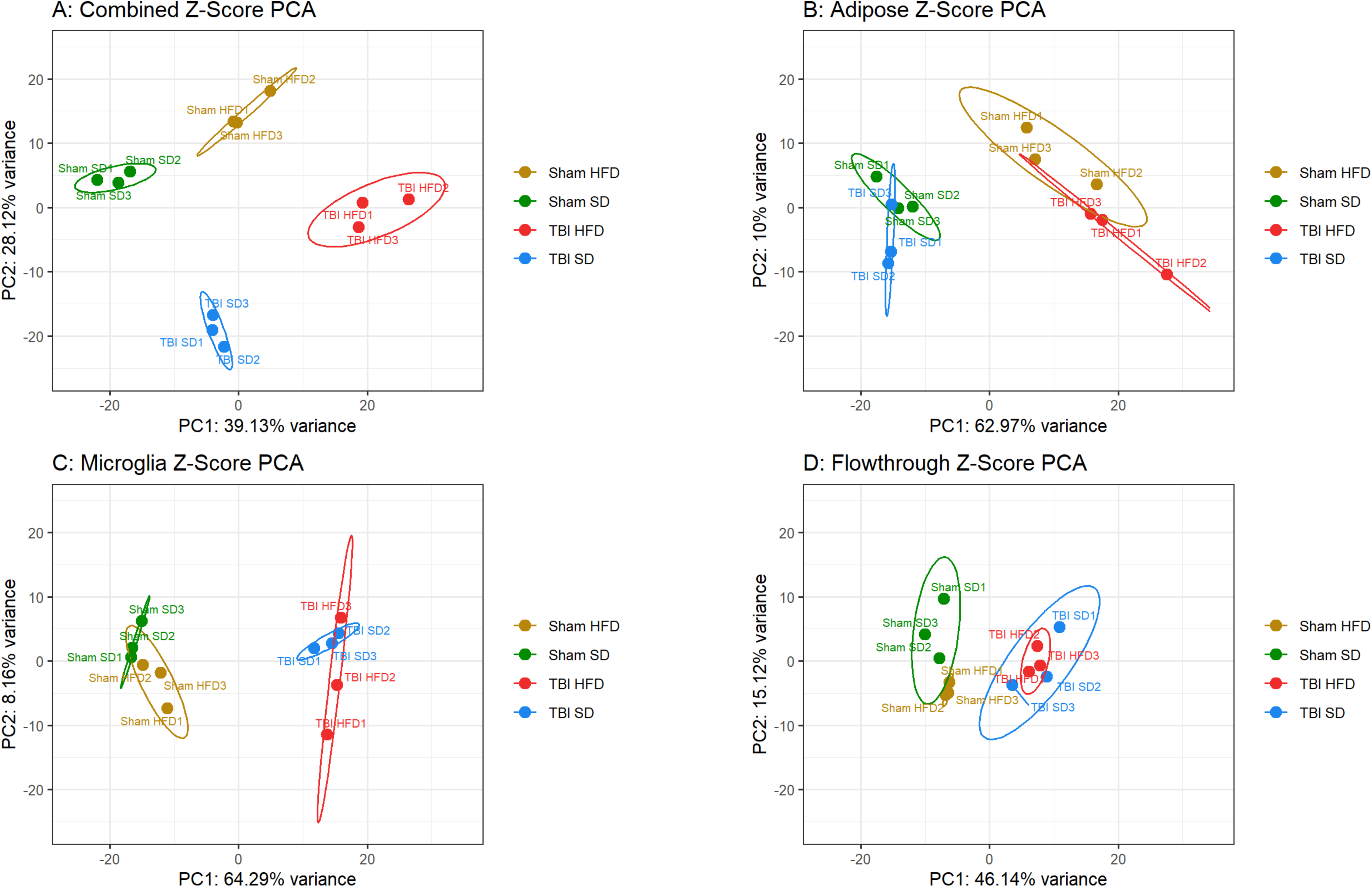
Combined HFD and TBI drive the global separation of cellular phenotypes. **A.** PCA analysis applied to the integrated data set (brain microglia and flowthrough as well as VAT) for every experimental animal reveals a robust separation of all groups with TBI-HFD as the sum of the individual vectors of change - TBI and HFD factors. **B.** HFD is the primary factor of change in the VAT. **C, D.** Microglia and flowthrough show TBI as the primary factor for change.

## 4. DISCUSSION

These studies examined the individual and combined effects of HFD and TBI on brain and VAT with a specific focus on cellular transcriptomic programs and cognitive changes. At 28 dpi (cohort 1), HFD is the major driver of time-dependent changes in body weight, metabolic markers, VAT weight, and immune cell infiltrate. In contrast, TBI induced neither independent effects (except elevating leptin) nor significant interaction on these parameters. HFD modified the VAT immune environment with significantly increased expression of the pro-inflammatory mediators *TNF-a, NLRP3, NOX-2,* and *p22^phox^*as well as the anti-inflammatory mediator *IL-10*. TBI plus HFD showed significant diet*injury interactions with exacerbation of HFD-dependent increases in VAT pro-inflammatory molecules *IL-1β* and *NLRP3,*indicating a brain trauma-dependent amplification of diet-induced adipose tissue inflammation.

In addition to systemic inflammatory responses [16, 48], diet-induced obesity is associated with brain neuroinflammation [16], including in the hypothalamus, leading to leptin and insulin resistance and weight gain [49, 50]. There are conflicting reports regarding the effect of HFD and obesity on extra-hypothalamic inflammation [21, 37, 51–53], with some studies reporting no changes [52, 53] and others identifying neuroinflammation in the hippocampus and cortex [37, 51]. Isolated cortical microglia from mice fed on a HFD regime released more TNF-α compared to SD-fed counterparts [54]. Diet-induced obesity, unlike short-term HFD feeding, leads to altered hippocampal microglial states, reductions in dendritic spines at sites of excitatory synapses and memory impairments; the cognitive deficits were reversed by partial knockdown of microglia and associated reduction in microglial phagocytosis [21]. In TBI patients and experimental models, microglia undergo a chronic transformation to reactive phenotypes that are associated with neurodegenerative processes and cognitive decline [10, 11]. Delayed transient removal of microglia after experimental TBI reduced the inflammatory lesion microenvironment and improved long-term neurological recovery[8]. Although previous studies have reported increased microglial altered states in the hypothalamus and the prefrontal cortex in the context of co-morbid diet-induced obesity and TBI [23, 26], the present study is the first to investigate hippocampal and cortical inflammatory responses at 28dpi (*cohort 1*). Unlike TBI, HFD significantly altered only two pro-inflammatory genes (*p22^phox^*and *CD11b*) in the hippocampus. However, combined TBI-HFD displayed a significant diet*injury interaction, including a elevation in hippocampal *p22^phox^* compared to TBI-SD mice. These results suggest that pre-existing and concurrent HFD primes the development of TBI-induced, pro-inflammatory responses.

To better characterize the interaction between HFD and TBI with regard to secondary injury, we examined cognitive (memory/learning) functions and large-scale transcriptomic changes in specific cellular compartments after chronic brain trauma as late as 90 dpi (*cohort 2*). The Y-Maze) and the NOR showed that both TBI and HFD had significant effects, but no significant diet*injury interaction was observed, suggesting a simple additive association. In contrast, HFD and TBI had significant factor effects in MWM but no additive effects. However, combined TBI-HFD caused an additive deficit with regard to MWM search strategy outcome.

Transcriptomic studies have proposed the use of transcriptome signatures to define the complex and dynamic microglial states [55]. To examine the impact of diet and injury on the molecular pathways potentially related to neurodegeneration and neurological dysfunction, we isolated and analyzed transcriptomic profiles of microglial populations from the perilesional cortex and hippocampus. The heatmap of microglia gene differential expression at 90 dpi displayed a consistent and ordered clustering of the experimental groups: Sham-SD, Sham-HFD, TBI-SD and TBI-HFD. The secondary grouping of Sham-SD with Sham-HFD and TBI-SD with TBI-HFD demonstrate that TBI is the dominant factor responsible for change, with HFD having a secondary, albeit persistent, impact. Major components of differential expression in response to TBI and/or HFD include downregulation of homeostatic microglia markers and upregulation of various microglia altered states, including disease-associated microglia (DAM). Homeostatic microglia actively support brain processes during physiological conditions (basal state). Multiple altered states are induced in response to pathological conditions (injury and/or disease) [55, 56] and may serve an adaptive role when acting to promote repair, but may also represent a maladaptive response when associated with dysfunctional proinflammatory states that perpetuate tissue damage[56]. The transcriptomic signature of the DAM and related microglia neurodegenerative phenotype (MGnD) state is one of the most consistent differentially expressed profile observed, although as with all other microglia pro-inflammatory signatures it reflects TBI, but not HFD, as a significant factor. The collective magnitude of overexpression of virtually all DAM/MGnD markers-including ApoE, Spp1, Trem2, Axl, Clec7a and Lpl [55, 56]-is paralleled by opposite directional changes for homeostatic markers, indicating a strong shift of microglial populations toward the DAM reactive phenotype after TBI/HFD. Downregulation of the homeostatic signature is one of the most common microglial changes shared across various neurodegenerative conditions [55]. The DAM/MGnD state, driven by the Trem2-APOE signaling pathway [57], is found in many neurodegenerative disorders, but the functional significance of the DAM signature remains to be elucidated [56]. *Trem2*, which along with *APOE* is elevated in microglia after TBI in our study, plays a key role in microglial responses to pathological signals such as amyloid accumulation, promoting their exit from the homeostatic state and shift to the DAM reactive state [58], especially DAM stage 2 [56]. Trem2 may be commonly required for microglia to switch from the homeostatic state to all reactive states [56]. Conversely, Trem2 absence impairs phagocytosis of key substrates, including APOE, resulting in an impaired response to beta-amyloid, suggesting that at least in certain circumstances the Trem2-dependent microglia shift to the DAM state may be an adaptive response [58].

In contrast to the differential expression approach, which identified only TBI as a significant factor, IPA analysis using a pathway-centric approach detected a significant effect for TBI and HFD, as well as the significant interaction responsible for the superadditive activation of multiple microglial reactive inflammatory pathways in combined TBI-HFD compared to TBI-SD.

Overall, our data indicate that in the more chronic phase (90 dpi), rather than returning to the homeostatic phenotype, microglia remain strongly shifted toward a heterogenous collection of reactive states/responses primarily driven by TBI, but with HFD having a secondary yet persistent effect. The widespread activation of numerous inflammatory pathways, increased presence of multiple reactive microglia states, and decreased homeostatic phenotype support the hypothesis that by 90 dpi a chronic post-TBI/HFD brain inflammatory disorder has been established. Thus microglia, due to inability to promote effective repair and especially in the presence of an additional insult (HFD), enter into persistent altered states that are maladaptive and may propagate secondary injury.

The gene expression changes in the non-microglia populations display TBI-dependent down-regulation of neuronal responses and neurogenesis, as well as upregulation of astrocyte and oligodendrocyte signatures, that are consistent with neurodegeneration [59, 60]. Notably, HFD is also able to decrease the neuronal signature. Pro-inflammatory changes were also found in non-microglia population, suggesting that microglia are not the only cell population involved in post-traumatic inflammatory responses.

The analysis of VAT shows an ordered clustering of differential gene expression, which in contrast to brain, establishes diet as the primary factor driving changes. Thus, DAM molecules as well as markers of various pro-inflammatory states, likely expressed in adipose tissue macrophages, are among the most numerous HFD/TBI upregulated genes, whereas other immune and epigenetic regulators are downregulated. Importantly, unlike in the brain, the VAT displays a significant effect of both TBI and HFD, and/or the presence of significant interactions driving superadditive pro-inflammatory effects. The TBI/HFD-dependent elevation was evident not only for the inflammatory states but also for the homeostatic state, suggesting a complex response in which HFD increases some macrophage sub-populations, whereas the combined TBI/HFD increases all macrophage sub-populations, typically at higher levels than HFD alone. IPA pathway analysis extends these findings, identifying the inflammatory pathways as the largest component of upregulated pathways, with both diet and injury as significant factors (although diet remains the dominant factor), and a significant diet-injury interaction driving a synergistic increase in the pathway activation levels after combined TBI-HFD compared to Sham-HFD.

Adipose tissue macrophages (ATM), originating from recruited circulating monocytes, accumulate lipids in obese mice and become polarized toward the lipid-associated macrophages (LAM) state [61]. LAM development is driven by Trem2 and characterized by upregulation of components of an enzymatic machinery that recognizes, scavenges and catabolizes lipids such as fatty acids transporter Cd36, fatty acid binding proteins 5 (Fabp5) and lipoprotein lipase (Lpl). Notably, these and other molecules defining LAM establish a profile that is virtually identical with the conserved signature for DAM [62]. The LAM/DAM signature in VAT samples is elevated by HFD and appears to be a Trem2-dependent uniform response and responsible, at least initially, for the clearance of injured cells and damaged tissue, resolution of inflammation, and improvement of metabolic changes in obesity [63, 64]. However, ATM also transform to pro-inflammatory phenotypes, including pro-inflammatory associated genes, changes that may underpin the disease-driving role played by ATM in metabolic conditions and obesity-associated inflammation [65]. As with the DAM state in the brain, which depending on the biological context, may change from adaptive [62] to maladaptive [57], the LAM state may undergo a disease-promoting pro-inflammatory shift [66]. This transformation may be triggered by conditions in which the initial cellular responses are modified by persistent injury and/or in multiple conditions such as combined TBI/HFD. A self-reinforcing loop involving pro-inflammatory molecules released by VAT that promote neurodegeneration [67] and chronic systemic inflammation triggered by TBI [68] may be responsible for the amplification of disease processes in TBI/HFD. Principal component analysis (PCA) illustrates the predominant importance of injury for brain changes and diet for VAT changes but also highlights the clear separation of collective responses across the various experimental groups, when a composite score including both brain and VAT changes, is generated.

Overall, our findings support the existence of bi-directional interactions between HFD and TBI. HFD enhances TBI-induced brain neuroinflammation and TBI augments HFD-induced VAT inflammatory changes, perhaps resulting in a self-propagating secondary injury loop responsible for chronic cognitive decline. One of the most surprising observations is that the VAT offers stronger evidence than the brain for the effect of HFD-TBI interaction to amplify the inflammatory responses, indicating that targeting the obesity co-morbidity may represent a potential therapeutic strategy for those TBI patients.

## Abbreviations

TBI: traumatic brain injury
CCI: controlled cortical impact
DIO: diet-induced obesity
HFD: high fat diet
SD: standard diet
VAT: visceral adipose tissue
DAM: disease associated microglia
MWM: morris water maze
NOR: novel object recognition

## Acknowledgements

We are grateful for the technical support provided by Xiaoyi Lin and Cera Huffman.

## Data Availability

Data will be made available upon request.

---

**Supplementary Figure 1: Morpheus unsupervised hierarchical clustering heat map on isolated microglia (CD11b^+^) cells.** Although only the genes that showed evidence of significant changes (p<0.05 in at least one of the two-way ANOVA outcomes) were included (as for Fig. 8) the pattern remained to Fig. 5A.

**Supplementary Figure 2: IPA detects activated and inhibited Upstream Regulator-Genes pathways in microglia.** IPA determined the activation of pathways based on the inclusion of genes with p<0.05; all pathways with z-score >2.5 or <-2.5 in at least one group are presented. TBI, but not HFD (except for a modest significant factor effect in upregulated) had strong significant factor effects in both upregulated (including several inflammatory-related pathways) and downregulated pathways; no significant TBI-HFD interaction was observed.

**Supplementary Figure 3: IPA detects activated and inhibited Upstream Regulator-Drugs pathways in microglia.** IPA determined the activation of pathways based on the inclusion of genes showing changes in each group vs Sham-SD with p<0.05; all pathways with z-score >2.5 or <-2.5 in at least one group are presented. There was a significant effect of TBI, HFD, and TBI-HFD interaction in upregulated upstream drug pathways which included several inflammatory pathways. Notably, TBI-HFD showed increased effects compared to TBI-SD. However, only TBI effects were reported on downregulated upstream drugs in microglia.

**Supplementary Figure 4: IPA detects upregulated and downregulated Diseases and Bio Functions pathways in microglia.** IPA determined the activation of pathways based on the inclusion of genes with p<0.05; all pathways with z-score >2.5 or <-2.5 in at least one group are presented. TBI, but not HFD, or TBI-HFD effects are reported on upregulated and downregulated diseases in microglia.

**Supplementary Figure 5: Morpheus unsupervised hierarchical clustering heat map on microglia vs. flowthrough cellular transcriptome.** The ratio between global microglia and flowthrough normalized expression data (560 genes out of a total of 769 genes had average expression >50) was used to determine the top 50 overexpressed and under-expressed genes in microglia vs flowthrough.

**Supplementary Figure 6: Morpheus unsupervised hierarchical clustering heat map on flowthrough cellular transcriptome.** Morpheus heat map on all 560 genes shows no ordered clustering across the group’s indicative of the smaller number of genes with consistent and significant changes.

**Supplementary Figure 7: Morpheus unsupervised hierarchical clustering heat map on flowthrough cellular transcriptome selected for genes undergoing significant expression changes.** When only genes showing significant changes (p<0.05 in at least one of the two-way ANOVA outcomes) were selected the heat map demonstrated the same ordered clustering across the groups observed in microglia. Genes downregulated included neurogenesis and neuronal signaling markers while genes upregulated included oligodendrocyte phenotype markers.

**Supplementary Figure 8: IPA detects Upstream Regulator-Genes upregulated in flowthrough cellular transcriptome.** IPA determined the activation of pathways based on the inclusion of genes with p<0.05. TBI, but no HFD or TBI-HFD showed significant effects on upstream genes upregulated in flowthrough cellular transcriptome.

**Supplementary Figure 9: IPA detects upregulated and downregulated Diseases and Bio Functions pathways in flowthrough cellular transcriptome.** IPA determined the activation of pathways based on the inclusion of genes with p<0.05; all pathways with z-score >2.0 or <-2.0 in at least one group are presented. Significant TBI, but no HFD or TBI-HFD interaction effects are observed in diseases upregulated and downregulated pathways in flowthrough cellular transcriptome. Notably, TBI-HFD showed increased effects compared to TBI-SD in upregulated pathways.

**Supplementary Figure 10: Morpheus unsupervised hierarchical clustering heat map on VAT cellular transcriptome.** Only genes that showed evidence of significant changes (p<0.05 in at least one of the two-way ANOVA outcomes) were included (369 genes). The heat map demonstrated the same ordered clustering across the groups observed Fig.9A.

**Supplementary Figure 11: IPA detects Upstream Regulator-Genes upregulated and downregulated genes in VAT.** IPA determined the activation of pathways based on the inclusion of genes with p<0.05; all pathways with z-score >2.8 or <-2.8 in at least one group are presented. Analysis of the Z-score of adipose upstream upregulated genes pathways including multiple inflammatory-related, identify significant TBI, HFD, and TBI-HFD interaction effects. Notably, TBI-HFD significantly increases upregulated pathways compared to TBI-SD. Only diet was a significant factor for downregulated genes.

**Supplementary Figure 12: IPA detects activated and inhibited Upstream Regulator-Drugs pathways in VAT.** IPA determined the activation of pathways based on the inclusion of genes with p<0.05; all pathways with z-score >2.8 or <-2.8 in at least one group are presented. Analysis of the Z-score of adipose upstream upregulated and downregulated drugs pathways, including multiple inflammatory-related, identify significant TBI, HFD, and TBI-HFD interaction effects. Notably, TBI-HFD significantly increases upregulated pathways and decreases downregulated pathways compared to TBI-SD.

**Supplementary Figure 13: IPA detects upregulated and downregulated Diseases and Bio Functions pathways in VAT.** IPA determined the activation of pathways based on the inclusion of genes with p<0.05; all pathways with z-score >3.5 or <-3.5 in at least one group are presented. Analysis of the Z-score of adipose disease pathways, including multiple inflammatory-related, identify significant TBI, HFD, and TBI-HFD interaction effects (except lack of significant injury effect in downregulated pathways). Notably, TBI-HFD significantly increases upregulated pathways and decreases downregulated pathways compared to TBI-SD.

**Supplementary Figure 14: Statistical analysis tables for the main figures**. We present the statistical tables for all the transcriptomic (Nanostring) and pathways analysis (IPA) data included in the main figures. The corresponding figure numbers are indicated.

## References

1. Corrigan, J.D., A.W. Selassie, and J.A. Orman, The epidemiology of traumatic brain injury. J Head Trauma Rehabil, 2010. 25(2): p. 72–80.

2. Chooi, Y.C., C. Ding, and F. Magkos, The epidemiology of obesity. Metabolism, 2019. 92: p. 6–10.

3. Winfield, R.D. and G.V. Bochicchio, The critically injured obese patient: a review and a look ahead. J Am Coll Surg, 2013. 216(6): p. 1193–206.

4. Chabok, S.Y., et al., The impact of body mass index on treatment outcomes among traumatic brain injury patients in intensive care units. Eur J Trauma Emerg Surg, 2014. 40(1): p. 51–5.

5. Christmas, A.B., et al., Morbid obesity impacts mortality in blunt trauma. Am Surg, 2007. 73(11): p. 1122–5.

6. Ditillo, M., et al., Morbid obesity predisposes trauma patients to worse outcomes: a National Trauma Data Bank analysis. J Trauma Acute Care Surg, 2014. 76(1): p. 176–9.

7. Mishra, R., et al., Obesity as a predictor of outcome following traumatic brain injury: A systematic review and meta-analysis. Clin Neurol Neurosurg, 2022. 217: p. 107260.

8. Henry, R.J., et al., Microglial Depletion with CSF1R Inhibitor During Chronic Phase of Experimental Traumatic Brain Injury Reduces Neurodegeneration and Neurological Deficits. J Neurosci, 2020. 40(14): p. 2960–2974.

9. Johnson, V.E., et al., Inflammation and white matter degeneration persist for years after a single traumatic brain injury. Brain, 2013. 136(Pt 1): p. 28–42.

10. Loane, D.J., et al., Progressive neurodegeneration after experimental brain trauma: association with chronic microglial activation. J Neuropathol Exp Neurol, 2014. 73(1): p. 14–29.

11. Simon, D.W., et al., The far-reaching scope of neuroinflammation after traumatic brain injury. Nat Rev Neurol, 2017. 13(3): p. 171–191.

12. Chong, A.J., et al., Effects of a High-Fat Diet on Neuroinflammation and Apoptosis in Acute Stage After Moderate Traumatic Brain Injury in Rats. Neurocrit Care, 2020. 33(1): p. 230–240.

13. Aguilar-Valles, A., et al., Obesity, adipokines and neuroinflammation. Neuropharmacology, 2015. 96(Pt A): p. 124–34.

14. Bousquet, M., et al., High-fat diet exacerbates MPTP-induced dopaminergic degeneration in mice. Neurobiol Dis, 2012. 45(1): p. 529–38.

15. Bruce-Keller, A.J., J.N. Keller, and C.D. Morrison, Obesity and vulnerability of the CNS. Biochim Biophys Acta, 2009. 1792(5): p. 395–400.

16. Miller, A.A. and S.J. Spencer, Obesity and neuroinflammation: a pathway to cognitive impairment. Brain Behav Immun, 2014. 42: p. 10–21.

17. Purkayastha, S. and D. Cai, Neuroinflammatory basis of metabolic syndrome. Mol Metab, 2013. 2(4): p. 356–63.

18. Campillo, B.W., et al., Short-term high-fat diet alters the mouse brain magnetic resonance imaging parameters consistently with neuroinflammation on males and metabolic rearrangements on females. A pre-clinical study with an optimized selection of linear mixed-effects models. Front Neurosci, 2022. 16: p. 1025108.

19. Valdearcos, M., et al., Microglial Inflammatory Signaling Orchestrates the Hypothalamic Immune Response to Dietary Excess and Mediates Obesity Susceptibility. Cell Metab, 2017. 26(1): p. 185–197 e3.

20. Valdearcos, M., et al., Microglia dictate the impact of saturated fat consumption on hypothalamic inflammation and neuronal function. Cell Rep, 2014. 9(6): p. 2124–38.

21. Cope, E.C., et al., Microglia Play an Active Role in Obesity-Associated Cognitive Decline. J Neurosci, 2018. 38(41): p. 8889–8904.

22. Henn, R.E., et al., Obesity-induced neuroinflammation and cognitive impairment in young adult versus middle-aged mice. Immun Ageing, 2022. 19(1): p. 67.

23. Sherman, M., et al., Adult obese mice suffer from chronic secondary brain injury after mild TBI. J Neuroinflammation, 2016. 13(1): p. 171.

24. Thomson, S., et al., Impact of High Fat Consumption on Neurological Functions after Traumatic Brain Injury in Rats. J Neurotrauma, 2022. 39(21-22): p. 1547–1560.

25. Ibeh, S., et al., High fat diet exacerbates long-term metabolic, neuropathological, and behavioral derangements in an experimental mouse model of traumatic brain injury. Life Sci, 2023. 314: p. 121316.

26. Karelina, K., et al., Traumatic brain injury and obesity induce persistent central insulin resistance. Eur J Neurosci, 2016. 43(8): p. 1034–43.

27. Byrnes, K.R., et al., Delayed mGluR5 activation limits neuroinflammation and neurodegeneration after traumatic brain injury. J Neuroinflammation, 2012. 9: p. 43.

28. Piao, C.S., et al., Late exercise reduces neuroinflammation and cognitive dysfunction after traumatic brain injury. Neurobiol Dis, 2013. 54: p. 252–63.

29. Catrysse, L. and G. van Loo, Adipose tissue macrophages and their polarization in health and obesity. Cell Immunol, 2018. 330: p. 114–119.

30. Kane, H. and L. Lynch, Innate Immune Control of Adipose Tissue Homeostasis. Trends Immunol, 2019. 40(9): p. 857–872.

31. Debette, S., et al., Midlife vascular risk factor exposure accelerates structural brain aging and cognitive decline. Neurology, 2011. 77(5): p. 461–8.

32. Fitzpatrick, A.L., et al., Midlife and late-life obesity and the risk of dementia: cardiovascular health study. Arch Neurol, 2009. 66(3): p. 336–42.

33. Whitmer, R.A., et al., Body mass index in midlife and risk of Alzheimer disease and vascular dementia. Curr Alzheimer Res, 2007. 4(2): p. 103–9.

34. Fourrier, C., et al., Brain tumor necrosis factor-alpha mediates anxiety-like behavior in a mouse model of severe obesity. Brain Behav Immun, 2019. 77: p. 25–36.

35. Gaudet, A.D., et al., miR-155 Deletion in Female Mice Prevents Diet-Induced Obesity. Sci Rep, 2016. 6: p. 22862.

36. Guo, D.H., et al., Visceral adipose NLRP3 impairs cognition in obesity via IL-1R1 on CX3CR1+ cells. J Clin Invest, 2020. 130(4): p. 1961–1976.

37. Pepping, J.K., et al., Myeloid-specific deletion of NOX2 prevents the metabolic and neurologic consequences of high fat diet. PLoS One, 2017. 12(8): p. e0181500.

38. Vandanmagsar, B., et al., The NLRP3 inflammasome instigates obesity-induced inflammation and insulin resistance. Nat Med, 2011. 17(2): p. 179–88.

39. Barrett, J.P., et al., Interferon-beta Plays a Detrimental Role in Experimental Traumatic Brain Injury by Enhancing Neuroinflammation That Drives Chronic Neurodegeneration. J Neurosci, 2020. 40(11): p. 2357–2370.

40. Barrett, J.P., et al., NOX2 deficiency alters macrophage phenotype through an IL-10/STAT3 dependent mechanism: implications for traumatic brain injury. J Neuroinflammation, 2017. 14(1): p. 65.

41. Henry, R.J., et al., Inhibition of miR-155 Limits Neuroinflammation and Improves Functional Recovery After Experimental Traumatic Brain Injury in Mice. Neurotherapeutics, 2019. 16(1): p. 216–230.

42. Henry, R.J., et al., Longitudinal Assessment of Sensorimotor Function after Controlled Cortical Impact in Mice: Comparison of Beamwalk, Rotarod, and Automated Gait Analysis Tests. J Neurotrauma, 2020. 37(24): p. 2709–2717.

43. Pavlidis, P., Using ANOVA for gene selection from microarray studies of the nervous system. Methods, 2003. 31(4): p. 282–9.

44. Marini, F. and H. Binder, pcaExplorer: an R/Bioconductor package for interacting with RNA-seq principal components. BMC Bioinformatics, 2019. 20(1): p. 331.

45. Rogers, J., et al., Search strategy selection in the Morris water maze indicates allocentric map formation during learning that underpins spatial memory formation. Neurobiol Learn Mem, 2017. 139: p. 37–49.

46. Curdt, N., et al., Search strategy analysis of Tg4-42 Alzheimer Mice in the Morris Water Maze reveals early spatial navigation deficits. Sci Rep, 2022. 12(1): p. 5451.

47. Colonna, M. and O. Butovsky, Microglia Function in the Central Nervous System During Health and Neurodegeneration. Annu Rev Immunol, 2017. 35: p. 441–468.

48. Gregor, M.F. and G.S. Hotamisligil, Inflammatory mechanisms in obesity. Annu Rev Immunol, 2011. 29: p. 415–45.

49. De Souza, C.T., et al., Consumption of a fat-rich diet activates a proinflammatory response and induces insulin resistance in the hypothalamus. Endocrinology, 2005. 146(10): p. 4192–9.

50. Posey, K.A., et al., Hypothalamic proinflammatory lipid accumulation, inflammation, and insulin resistance in rats fed a high-fat diet. Am J Physiol Endocrinol Metab, 2009. 296(5): p. E1003–12.

51. Jeon, B.T., et al., Resveratrol attenuates obesity-associated peripheral and central inflammation and improves memory deficit in mice fed a high-fat diet. Diabetes, 2012. 61(6): p. 1444–54.

52. Milanski, M., et al., Saturated fatty acids produce an inflammatory response predominantly through the activation of TLR4 signaling in hypothalamus: implications for the pathogenesis of obesity. J Neurosci, 2009. 29(2): p. 359–70.

53. Thaler, J.P., et al., Obesity is associated with hypothalamic injury in rodents and humans. J Clin Invest, 2012. 122(1): p. 153–62.

54. Puig, K.L., et al., Amyloid precursor protein and proinflammatory changes are regulated in brain and adipose tissue in a murine model of high fat diet-induced obesity. PLoS One, 2012. 7(1): p. e30378.

55. Paolicelli, R.C., et al., Microglia states and nomenclature: A field at its crossroads. Neuron, 2022. 110(21): p. 3458–3483.

56. Chen, Y. and M. Colonna, Microglia in Alzheimer’s disease at single-cell level. Are there common patterns in humans and mice? J Exp Med, 2021. 218(9).

57. Krasemann, S., et al., The TREM2-APOE Pathway Drives the Transcriptional Phenotype of Dysfunctional Microglia in Neurodegenerative Diseases. Immunity, 2017. 47(3): p. 566–581 e9.

58. McQuade, A., et al., Gene expression and functional deficits underlie TREM2-knockout microglia responses in human models of Alzheimer’s disease. Nat Commun, 2020. 11(1): p. 5370.

59. Brandebura, A.N., et al., Astrocyte contribution to dysfunction, risk and progression in neurodegenerative disorders. Nat Rev Neurosci, 2023. 24(1): p. 23–39.

60. Pandey, S., et al., Disease-associated oligodendrocyte responses across neurodegenerative diseases. Cell Rep, 2022. 40(8): p. 111189.

61. Reich, T., et al., TREM2 has a significant, gender-specific, effect on human obesity. Sci Rep, 2023. 13(1): p. 482.

62. Jaitin, D.A., et al., Lipid-Associated Macrophages Control Metabolic Homeostasis in a Trem2-Dependent Manner. Cell, 2019. 178(3): p. 686–698 e14.

63. Kim, K., et al., Characteristics of plaque lipid-associated macrophages and their possible roles in the pathogenesis of atherosclerosis. Curr Opin Lipidol, 2022. 33(5): p. 283–288.

64. Worthmann, A.H., J., TREM2-Positive Lipid-Associated Macrophages (LAMs) Control White Adipose Tissue Remodeling and Metabolic Adaptation in Obesity. Immunometabolism, 2020. 2(2):e200014.

65. Florance, I. and S. Ramasubbu, Current Understanding on the Role of Lipids in Macrophages and Associated Diseases. Int J Mol Sci, 2022. 24(1).

66. Prieur, X., et al., Differential lipid partitioning between adipocytes and tissue macrophages modulates macrophage lipotoxicity and M2/M1 polarization in obese mice. Diabetes, 2011. 60(3): p. 797–809.

67. Jin, Y., et al., Exosomes from Inflamed Macrophages Promote the Progression of Parkinson’s Disease by Inducing Neuroinflammation. Mol Neurobiol, 2023.

68. Kumar, R.G., J.A. Boles, and A.K. Wagner, Chronic Inflammation After Severe Traumatic Brain Injury: Characterization and Associations With Outcome at 6 and 12 Months Postinjury. J Head Trauma Rehabil, 2015. 30(6): p. 369–81.

