## Supplemental Figures for "Interaction of high-fat diet and brain trauma alters adipose tissue macrophages and brain microglia associated with exacerbated cognitive dysfunction"

### Figure 1-Supplemental

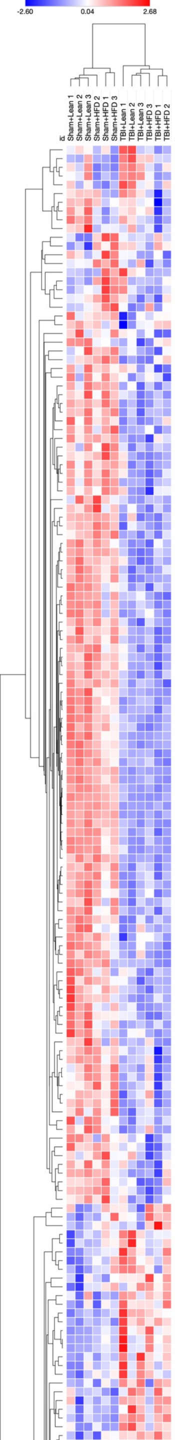

| Symbol | Average | Probe.Annotation | p_val_Diet | p_val_Condition | p_val_diet:condition | p_TBILean:ShamLean | p_ShamHFD:ShamLean | p_TBIHFD:ShamHFD | p_TBIHFD:ShamLean | p_TBIHFD:TBILean |
|---|---|---|---|---|---|---|---|---|---|---|
| Cd34 | 2237.20 | Cell Migration | 0.03 | 0.52 | 0.23 | 0.54 | 0.81 | 0.97 | 0.57 | 0.10 |
| Pccb | 333.62 | Lipid Metabolism | 0.02 | 0.01 | 0.51 | 0.10 | 0.50 | 0.38 | 0.99 | 0.15 |
| Pros1 | 1341.40 | Complement System | 0.00 | 0.03 | 0.87 | 0.35 | 0.03 | 0.26 | 0.45 | 0.04 |
| Nptn | 2270.57 | Neurogenesis | 0.00 | 0.09 | 0.32 | 0.23 | 0.06 | 0.93 | 0.15 | 0.01 |
| Cdc42 | 7587.42 | Neurogenesis | 0.04 | 0.06 | 0.87 | 0.39 | 0.42 | 0.51 | 1.00 | 0.32 |
| Bub3 | 420.11 | Cell Cycle | 0.01 | 0.61 | 0.21 | 0.94 | 0.57 | 0.57 | 0.11 | 0.05 |
| Sgce | 1063.86 | Cytoskeletal Dynamics | 0.04 | 0.08 | 0.57 | 0.74 | 0.59 | 0.31 | 0.05 | 0.22 |
| Hexb | 77311.57 | Microglia-Homeostatic | 0.02 | 0.37 | 0.53 | 0.68 | 0.13 | 1.00 | 0.09 | 0.42 |
| Lgmn | 35051.20 | Microglia Markers | 0.00 | 0.05 | 0.38 | 0.17 | 0.01 | 0.75 | 0.00 | 0.05 |
| Psmc2 | 790.77 | Wnt Signaling | 0.05 | 0.12 | 0.48 | 0.36 | 0.21 | 0.89 | 0.08 | 0.69 |
| Pias1 | 221.04 | JAK-STAT | 0.01 | 0.34 | 0.02 | 0.50 | 0.01 | 0.08 | 0.35 | 0.99 |
| Adamts1 | 1734.37 | Microglia Markers | 0.06 | 0.48 | 0.03 | 0.15 | 0.04 | 0.54 | 0.24 | 0.98 |
| Reck | 51.11 | Blood Brain Barrier | 0.64 | 0.04 | 0.18 | 0.89 | 0.55 | 0.09 | 0.53 | 0.90 |
| Slc25a12 | 173.35 | Myelogenesis | 0.74 | 0.01 | 0.01 | 1.00 | 0.07 | 0.00 | 0.18 | 0.15 |
| Tie1 | 61.47 | Endothelial Cell Function | 0.69 | 0.88 | 0.02 | 0.18 | 0.31 | 0.25 | 1.00 | 0.14 |
| Mkrn1 | 837.28 | Antigen Processing and Presentation | 0.03 | 0.16 | 0.01 | 0.55 | 0.01 | 0.03 | 0.87 | 0.93 |
| Icam2 | 69.15 | Cell Migration | 0.57 | 0.23 | 0.04 | 0.85 | 0.22 | 0.11 | 0.96 | 0.58 |
| Pak1 | 500.62 | T-cell signaling | 0.23 | 0.02 | 0.16 | 0.77 | 0.26 | 0.05 | 0.68 | 1.00 |
| Stmn1 | 198.90 | Neuronal Markers | 0.03 | 0.10 | 0.42 | 0.89 | 0.13 | 0.29 | 0.93 | 0.58 |
| Mtmr4 | 135.51 | Autophagy | 0.20 | 0.02 | 0.32 | 0.10 | 0.99 | 0.64 | 0.77 | 0.37 |
| Ticam1 | 167.68 | NO Metabolism and Signaling | 0.20 | 0.07 | 0.04 | 0.05 | 0.85 | 0.99 | 0.96 | 0.09 |
| Fos | 10885.67 | MAPK & PI3K | 0.30 | 0.04 | 0.45 | 0.65 | 1.00 | 0.17 | 0.13 | 0.56 |
| Camk2d | 711.71 | Calcium Signaling | 0.35 | 0.02 | 0.02 | 0.02 | 0.11 | 1.00 | 0.10 | 0.61 |
| Clcf1 | 232.49 | A2 Astrocyte | 0.91 | 0.02 | 0.34 | 0.55 | 0.92 | 0.09 | 0.21 | 0.85 |
| Nlrp1b | 117.14 | Inflammasome | 0.03 | 0.00 | 0.99 | 0.09 | 0.32 | 0.09 | 0.81 | 0.31 |
| Klrd1 | 92.56 | Innate Immune Response | 0.19 | 0.03 | 0.42 | 0.63 | 0.97 | 0.15 | 0.08 | 0.42 |
| Trim2 | 165.59 | Interferon Signaling | 0.94 | 0.05 | 0.94 | 0.42 | 1.00 | 0.37 | 0.37 | 1.00 |
| Fcgrt | 612.78 | Astrocyte Markers | 0.90 | 0.00 | 0.86 | 0.08 | 1.00 | 0.05 | 0.07 | 1.00 |
| Orc4 | 357.77 | Cell Cycle | 0.66 | 0.00 | 0.96 | 0.08 | 0.99 | 0.08 | 0.05 | 0.98 |
| Glul | 11142.06 | Glutamatergic Synapse | 0.39 | 0.00 | 0.67 | 0.00 | 0.99 | 0.00 | 0.00 | 0.78 |
| Trim26 | 369.14 | Interferon Signaling | 0.71 | 0.00 | 0.38 | 0.00 | 0.98 | 0.03 | 0.02 | 0.79 |
| Stab1 | 748.57 | Microglia-M2 | 0.88 | 0.00 | 0.13 | 0.01 | 0.71 | 0.35 | 0.08 | 0.59 |
| Kcnd1 | 129.19 | Ion Transport | 0.86 | 0.00 | 0.88 | 0.07 | 1.00 | 0.10 | 0.10 | 1.00 |
| Pik3r1 | 1353.58 | MAPK & PI3K | 0.71 | 0.00 | 0.77 | 0.05 | 1.00 | 0.09 | 0.10 | 0.96 |
| Atp2b2 | 63.29 | Ion Transport | 0.42 | 0.01 | 0.39 | 0.39 | 0.61 | 0.07 | 0.38 | 1.00 |
| Pecam1 | 273.61 | Blood Brain Barrier | 0.52 | 0.00 | 0.05 | 0.19 | 0.24 | 0.00 | 0.04 | 0.68 |
| Camk2g | 146.85 | Calcium Signaling | 0.46 | 0.00 | 0.65 | 0.12 | 1.00 | 0.05 | 0.04 | 0.82 |
| Ccr5 | 3460.55 | Primed Microglia | 0.70 | 0.00 | 0.48 | 0.12 | 0.99 | 0.03 | 0.04 | 0.85 |
| Stxbp1 | 103.95 | Neurogenesis | 0.74 | 0.00 | 0.51 | 0.03 | 0.88 | 0.01 | 0.02 | 0.99 |
| Scoc | 3003.06 | Primed Microglia | 1.00 | 0.03 | 0.63 | 0.44 | 0.98 | 0.18 | 0.28 | 0.98 |
| Tnfrsf17 | 168.96 | Cytokines | 0.78 | 0.01 | 0.99 | 0.15 | 1.00 | 0.15 | 0.20 | 1.00 |
| Il18 | 291.30 | Cytokines | 0.78 | 0.01 | 0.59 | 0.18 | 1.00 | 0.06 | 0.08 | 0.93 |
| Basp1 | 1180.28 | Cytoskeletal Dynamics | 0.68 | 0.00 | 0.58 | 0.07 | 1.00 | 0.02 | 0.02 | 0.89 |
| Nuak1 | 359.40 | Primed Microglia | 0.73 | 0.00 | 0.28 | 0.02 | 0.71 | 0.00 | 0.01 | 0.94 |
| Bin1 | 4118.82 | Microglia-Homeostatic | 0.07 | 0.00 | 0.37 | 0.01 | 0.22 | 0.00 | 0.02 | 0.86 |
| Tanc2 | 1622.34 | Cytoskeletal Dynamics | 0.75 | 0.00 | 0.39 | 0.01 | 0.81 | 0.06 | 0.02 | 0.97 |
| Tjp1 | 1213.62 | Gap Junctions | 0.57 | 0.00 | 0.45 | 0.00 | 1.00 | 0.00 | 0.00 | 0.76 |
| Agmo | 736.25 | Lipid Metabolism | 0.37 | 0.00 | 0.61 | 0.02 | 0.73 | 0.04 | 0.01 | 0.99 |
| Ggta1 | 464.13 | A1 Astrocyte | 0.59 | 0.00 | 0.26 | 0.00 | 0.61 | 0.02 | 0.00 | 0.96 |
| Prkce | 133.93 | Phagocytosis | 0.59 | 0.00 | 0.34 | 0.00 | 0.69 | 0.01 | 0.00 | 0.99 |
| Mef2a | 2304.30 | Microglia-Homeostatic | 0.05 | 0.00 | 0.23 | 0.01 | 0.13 | 0.09 | 0.00 | 0.89 |
| Cdk7 | 529.88 | Cell Cycle | 0.03 | 0.00 | 0.78 | 0.03 | 0.37 | 0.02 | 0.00 | 0.22 |
| Cables1 | 955.94 | Cell Cycle | 0.11 | 0.00 | 0.54 | 0.01 | 0.38 | 0.02 | 0.00 | 0.85 |
| Egr1 | 15802.23 | Microglia-Homeostatic | 0.41 | 0.00 | 0.23 | 0.02 | 0.47 | 0.22 | 0.03 | 0.99 |
| Ckb | 4250.45 | Astrocyte Markers | 0.75 | 0.00 | 0.52 | 0.01 | 0.99 | 0.02 | 0.02 | 0.89 |
| Tmem100 | 80.34 | Astrocyte Markers | 0.21 | 0.00 | 0.53 | 0.07 | 0.96 | 0.02 | 0.01 | 0.52 |
| Jun | 20389.28 | Microglia Markers | 0.32 | 0.00 | 0.90 | 0.01 | 0.91 | 0.01 | 0.00 | 0.83 |
| Crot | 401.40 | Lipid Metabolism | 0.35 | 0.00 | 0.96 | 0.05 | 0.88 | 0.05 | 0.02 | 0.90 |
| Cacnb2 | 116.27 | MAPK & PI3K | 0.14 | 0.00 | 0.34 | 0.02 | 0.97 | 0.00 | 0.00 | 0.31 |
| Gpr34 | 9652.79 | Microglia-Homeostatic | 0.16 | 0.00 | 0.71 | 0.02 | 0.85 | 0.01 | 0.00 | 0.56 |
| Siglech | 7087.94 | Microglia-Homeostatic | 0.43 | 0.00 | 0.34 | 0.00 | 0.58 | 0.01 | 0.00 | 1.00 |
| E2f3 | 250.78 | Cell Cycle | 0.27 | 0.00 | 0.51 | 0.00 | 0.98 | 0.00 | 0.00 | 0.58 |
| Ctsf | 1594.79 | Microglia Markers | 0.00 | 0.00 | 0.01 | 0.00 | 0.00 | 0.00 | 0.00 | 0.27 |
| Slco2b1 | 1071.17 | Microglia Markers | 0.01 | 0.00 | 0.02 | 0.00 | 0.01 | 0.00 | 0.00 | 1.00 |
| Csf1r | 23290.91 | Microglia-Homeostatic | 0.02 | 0.00 | 0.05 | 0.00 | 0.03 | 0.00 | 0.00 | 0.97 |
| Sall1 | 3347.39 | Microglia-Homeostatic | 0.01 | 0.00 | 0.06 | 0.00 | 0.01 | 0.00 | 0.00 | 0.66 |
| Tgfbr1 | 4126.54 | Microglia-Homeostatic | 0.07 | 0.00 | 0.49 | 0.00 | 0.27 | 0.00 | 0.00 | 0.77 |
| Slc7a8 | 1048.15 | Ion Transport | 0.00 | 0.00 | 0.00 | 0.00 | 0.00 | 0.00 | 0.00 | 0.99 |
| Elmo1 | 1303.74 | Primed Microglia | 0.02 | 0.00 | 0.02 | 0.00 | 0.02 | 0.00 | 0.00 | 1.00 |
| Cx3cr1 | 14097.44 | Microglia-Homeostatic | 0.00 | 0.00 | 0.13 | 0.00 | 0.02 | 0.00 | 0.00 | 0.45 |
| Itgam | 5262.64 | Microglia-Homeostatic | 0.15 | 0.00 | 0.41 | 0.00 | 0.36 | 0.01 | 0.00 | 0.95 |
| Cttnbp2nl | 659.52 | Microglia Markers | 0.03 | 0.00 | 0.05 | 0.00 | 0.03 | 0.00 | 0.00 | 1.00 |
| Pmepa1 | 1020.40 | Microglia-Homeostatic | 0.06 | 0.00 | 0.07 | 0.00 | 0.06 | 0.00 | 0.00 | 1.00 |
| P2ry12 | 19247.80 | Microglia-Homeostatic | 0.04 | 0.00 | 0.29 | 0.00 | 0.14 | 0.00 | 0.00 | 0.81 |
| Slc2a5 | 4454.63 | Microglia-Homeostatic | 0.09 | 0.00 | 0.30 | 0.00 | 0.22 | 0.00 | 0.00 | 0.93 |
| Smad3 | 227.80 | Microglia-Homeostatic | 0.60 | 0.00 | 0.40 | 0.00 | 0.75 | 0.00 | 0.00 | 0.99 |
| Rtn1 | 324.68 | Neurogenesis | 0.98 | 0.00 | 0.21 | 0.00 | 0.76 | 0.00 | 0.00 | 0.78 |
| Sparc | 11515.38 | Microglia-Homeostatic | 0.07 | 0.00 | 0.31 | 0.00 | 0.18 | 0.00 | 0.00 | 0.88 |
| P2ry13 | 3855.25 | Microglia-Homeostatic | 0.29 | 0.00 | 0.76 | 0.00 | 0.73 | 0.00 | 0.00 | 0.94 |
| Prkab1 | 811.95 | Glucose Metabolism | 0.25 | 0.00 | 0.07 | 0.00 | 0.16 | 0.00 | 0.00 | 0.93 |
| Stambpl1 | 506.90 | Primed Microglia | 0.17 | 0.00 | 0.37 | 0.00 | 0.36 | 0.01 | 0.00 | 0.98 |
| Adgrg1 | 554.46 | Microglia-Homeostatic | 0.99 | 0.00 | 0.60 | 0.00 | 0.98 | 0.00 | 0.00 | 0.98 |
| Tmem119 | 1299.18 | Microglia-Homeostatic | 0.70 | 0.00 | 0.99 | 0.01 | 0.99 | 0.01 | 0.00 | 0.99 |
| Entpd1 | 2851.46 | Microglia-Homeostatic | 1.00 | 0.00 | 0.69 | 0.01 | 0.99 | 0.02 | 0.01 | 0.99 |
| Cep68 | 267.01 | Primed Microglia | 0.97 | 0.00 | 0.16 | 0.00 | 0.68 | 0.03 | 0.01 | 0.71 |
| Etv5 | 653.38 | Neurogenesis | 0.24 | 0.00 | 0.16 | 0.01 | 0.27 | 0.10 | 0.01 | 1.00 |
| Tlr3 | 270.68 | NF-kappaB Signaling | 0.81 | 0.00 | 0.34 | 0.03 | 0.81 | 0.22 | 0.06 | 0.95 |
| Serinc3 | 15325.12 | Primed Microglia | 0.14 | 0.00 | 0.16 | 0.00 | 0.19 | 0.06 | 0.00 | 1.00 |
| Mertk | 2059.26 | Microglia-Homeostatic | 0.21 | 0.00 | 0.08 | 0.00 | 0.16 | 0.02 | 0.00 | 0.97 |
| Crkl | 219.40 | Neurotrophin Signaling | 0.04 | 0.00 | 0.07 | 0.01 | 0.05 | 0.52 | 0.01 | 0.99 |
| Mafb | 2263.07 | Microglia-Homeostatic | 0.95 | 0.01 | 0.23 | 0.03 | 0.78 | 0.38 | 0.11 | 0.82 |
| Usp2 | 1286.63 | Microglia Markers | 0.03 | 0.00 | 0.28 | 0.00 | 0.10 | 0.03 | 0.00 | 0.70 |
| Cdk5r1 | 124.53 | Neurotrophin Signaling | 0.14 | 0.00 | 0.64 | 0.06 | 0.84 | 0.02 | 0.01 | 0.47 |
| Ppm1l | 116.10 | Primed Microglia | 0.12 | 0.00 | 0.69 | 0.02 | 0.47 | 0.04 | 0.01 | 0.79 |
| Plxdc2 | 4537.41 | Microglia Markers | 0.02 | 0.01 | 0.22 | 0.04 | 0.07 | 0.47 | 0.01 | 0.69 |
| Map2k1 | 1033.94 | MAPK & PI3K | 0.31 | 0.00 | 0.38 | 0.02 | 0.52 | 0.10 | 0.01 | 1.00 |
| Mapk14 | 1061.74 | MAPK & PI3K | 0.15 | 0.00 | 0.57 | 0.01 | 0.45 | 0.05 | 0.01 | 0.89 |
| Nos1ap | 147.24 | Oxidative & Nitrosative Stress | 0.31 | 0.00 | 0.09 | 0.01 | 0.23 | 0.19 | 0.01 | 0.94 |
| Il10ra | 3397.40 | Cytokines | 0.02 | 0.00 | 0.12 | 0.01 | 0.04 | 0.14 | 0.00 | 0.83 |
| F11r | 3080.16 | Primed Microglia | 0.00 | 0.00 | 0.46 | 0.00 | 0.02 | 0.01 | 0.00 | 0.10 |
| Abcc3 | 604.17 | Oxidative & Nitrosative Stress | 0.04 | 0.00 | 0.03 | 0.00 | 0.03 | 0.15 | 0.00 | 1.00 |
| Sirpa | 5845.38 | NO Metabolism and Signaling | 0.00 | 0.00 | 0.08 | 0.01 | 0.01 | 0.32 | 0.00 | 0.32 |
| Slc46a1 | 296.74 | Ion Transport | 0.38 | 0.02 | 0.33 | 0.08 | 0.53 | 0.52 | 0.09 | 1.00 |
| Olfml3 | 6107.95 | Microglia-Homeostatic | 0.25 | 0.00 | 0.60 | 0.15 | 0.96 | 0.05 | 0.03 | 0.60 |
| Fcrls | 11065.30 | Microglia-Homeostatic | 0.46 | 0.01 | 0.99 | 0.10 | 0.94 | 0.10 | 0.05 | 0.95 |
| Calm2 | 1559.96 | Calcium Signaling | 0.62 | 0.01 | 0.71 | 0.20 | 0.92 | 0.09 | 0.23 | 1.00 |
| Cst3 | 183431.23 | Microglia-Homeostatic | 0.78 | 0.01 | 0.66 | 0.25 | 0.95 | 0.11 | 0.22 | 1.00 |
| Gng2 | 888.44 | Serotonergic Synapse | 0.41 | 0.01 | 0.41 | 0.35 | 0.63 | 0.07 | 0.35 | 1.00 |
| Ltc4s | 240.93 | Microglia Markers | 0.64 | 0.03 | 0.94 | 0.28 | 0.99 | 0.32 | 0.45 | 0.98 |
| Rhob | 2475.08 | Primed Microglia | 0.71 | 0.00 | 1.00 | 0.07 | 0.99 | 0.07 | 0.10 | 0.99 |
| Adrb1 | 225.98 | Neuroactive Ligands and Receptors | 0.26 | 0.00 | 0.89 | 0.07 | 0.77 | 0.06 | 0.21 | 0.87 |
| Ap3m2 | 159.39 | Primed Microglia | 0.76 | 0.02 | 0.38 | 0.10 | 0.82 | 0.52 | 0.18 | 0.97 |
| Dnajb9 | 869.26 | Proteotoxic Stress | 0.96 | 0.02 | 0.25 | 0.07 | 0.84 | 0.66 | 0.26 | 0.80 |
| Hacd3 | 275.15 | Lipid Metabolism | 0.04 | 0.02 | 0.98 | 0.27 | 0.40 | 0.26 | 0.03 | 0.38 |
| Dagla | 190.54 | Neurogenesis | 0.08 | 0.00 | 0.22 | 0.15 | 0.96 | 0.01 | 0.01 | 0.16 |
| B4galt6 | 443.22 | Oligodendrocyte Differentiation | 0.71 | 0.04 | 0.05 | 1.00 | 0.55 | 0.04 | 0.28 | 0.30 |
| Adora3 | 634.29 | Microglia-Homeostatic | 0.32 | 0.03 | 0.95 | 0.28 | 0.86 | 0.32 | 0.11 | 0.89 |
| Rab6b | 479.49 | Primed Microglia | 0.62 | 0.01 | 0.48 | 0.06 | 0.82 | 0.24 | 0.07 | 1.00 |
| Ywhah | 2798.78 | Neurogenesis | 0.77 | 0.04 | 0.74 | 0.46 | 0.97 | 0.25 | 0.44 | 1.00 |
| Atp8a2 | 816.29 | Primed Microglia | 0.34 | 0.00 | 0.34 | 0.21 | 1.00 | 0.03 | 0.03 | 0.52 |
| Lpar6 | 889.95 | Purinergic Signaling | 0.74 | 0.03 | 0.67 | 0.42 | 1.00 | 0.19 | 0.21 | 0.94 |
| Gphn | 80.37 | GABAergic Synapse | 0.03 | 0.42 | 0.37 | 1.00 | 0.65 | 0.61 | 0.14 | 0.13 |
| Pdha1 | 263.32 | Glucose Metabolism | 0.03 | 0.12 | 0.97 | 0.65 | 0.30 | 0.62 | 0.05 | 0.28 |
| Rnasel | 282.95 | Interferon Signaling | 0.01 | 0.18 | 0.20 | 1.00 | 0.40 | 0.25 | 0.02 | 0.03 |
| Psmd10 | 150.67 | Antigen Processing and Presentation | 0.50 | 0.02 | 0.01 | 0.01 | 0.33 | 0.99 | 0.47 | 0.09 |
| Nes | 85.66 | Cytoskeletal Dynamics | 0.77 | 0.05 | 0.05 | 0.04 | 0.51 | 1.00 | 0.52 | 0.31 |
| Chchd3 | 203.02 | Glucose Metabolism | 0.85 | 0.02 | 0.09 | 0.04 | 0.64 | 0.91 | 0.31 | 0.48 |
| Ap2m1 | 1436.19 | Gap Junctions | 0.51 | 0.02 | 0.03 | 0.02 | 0.18 | 0.99 | 0.12 | 0.58 |
| Lta4h | 355.30 | Lipid Metabolism | 0.93 | 0.03 | 0.01 | 0.01 | 0.17 | 0.97 | 0.31 | 0.15 |
| Calm1 | 1579.55 | Calcium Signaling | 0.31 | 0.02 | 0.57 | 0.47 | 0.98 | 0.16 | 0.10 | 0.65 |
| Kctd12 | 815.88 | GABAergic Synapse | 0.05 | 0.05 | 0.22 | 0.13 | 0.12 | 0.90 | 0.05 | 0.89 |
| S100b | 56.40 | Astrocyte Markers | 0.47 | 0.02 | 0.35 | 0.59 | 0.63 | 0.10 | 0.50 | 1.00 |
| Ahcyl1 | 424.70 | Astrocyte Markers | 0.86 | 0.03 | 0.77 | 0.23 | 1.00 | 0.40 | 0.36 | 0.98 |
| Lamtor3 | 487.58 | Autophagy | 0.20 | 0.00 | 0.06 | 0.00 | 0.94 | 0.16 | 0.07 | 0.13 |
| Akt3 | 255.87 | MAPK & PI3K | 0.45 | 0.00 | 0.35 | 0.01 | 1.00 | 0.04 | 0.04 | 0.61 |
| Mog | 73.55 | Oligodendrocyte Markers | 0.67 | 0.01 | 0.39 | 0.08 | 0.78 | 0.43 | 0.12 | 0.99 |
| Plp1 | 974.39 | Oligodendrocyte Markers | 0.55 | 0.04 | 0.63 | 0.22 | 0.86 | 0.52 | 0.20 | 1.00 |
| Ermn | 63.14 | Oligodendrocyte Markers | 0.69 | 0.04 | 0.56 | 0.21 | 0.89 | 0.57 | 0.24 | 1.00 |
| Iigp1 | 78.56 | Interferon Signaling | 0.47 | 0.01 | 0.45 | 0.09 | 1.00 | 0.41 | 0.39 | 0.70 |
| Mapt | 50.13 | Neurogenesis | 0.52 | 0.04 | 0.19 | 0.09 | 0.94 | 0.88 | 0.59 | 0.48 |
| Inpp5f | 156.89 | Neurogenesis | 0.92 | 0.04 | 0.04 | 0.04 | 0.36 | 1.00 | 0.33 | 0.42 |
| Kpna1 | 313.34 | Interferon Signaling | 0.36 | 0.05 | 0.41 | 0.18 | 1.00 | 0.72 | 0.76 | 0.59 |
| Ptpn1 | 1448.95 | Interferon Signaling | 0.30 | 0.03 | 0.69 | 0.43 | 0.71 | 0.20 | 0.69 | 0.96 |
| Tnfrsf11a | 612.49 | Cytokines | 0.67 | 0.04 | 0.95 | 0.32 | 0.99 | 0.37 | 0.49 | 0.98 |
| Hadh | 198.57 | Lipid Metabolism | 0.47 | 0.01 | 0.75 | 0.12 | 0.99 | 0.23 | 0.34 | 0.87 |
| Fgf13 | 96.85 | Neurogenesis | 0.30 | 0.00 | 0.62 | 0.01 | 0.97 | 0.03 | 0.05 | 0.67 |
| Opa1 | 246.13 | Neurogenesis | 0.21 | 0.01 | 0.17 | 0.02 | 1.00 | 0.45 | 0.40 | 0.25 |
| Atp5g3 | 408.73 | Glucose Metabolism | 0.38 | 0.01 | 0.79 | 0.09 | 0.83 | 0.15 | 0.05 | 0.97 |
| Rapgef2 | 82.10 | MAPK & PI3K | 0.52 | 0.01 | 0.66 | 0.12 | 0.85 | 0.30 | 0.10 | 1.00 |
| Ptpn11 | 376.02 | Neurotrophin Signaling | 0.60 | 0.01 | 0.77 | 0.22 | 1.00 | 0.12 | 0.09 | 0.93 |
| Rab1a | 2571.69 | Autophagy | 0.63 | 0.01 | 0.15 | 0.57 | 0.86 | 0.03 | 0.09 | 0.49 |
| Gls | 1318.50 | Glutamatergic Synapse | 0.15 | 0.05 | 0.41 | 0.74 | 0.96 | 0.18 | 0.09 | 0.36 |
| Igfbp5 | 52.60 | Insulin Signaling | 0.07 | 0.02 | 0.11 | 0.87 | 1.00 | 0.04 | 0.03 | 0.10 |
| Pkd2 | 148.52 | NO Metabolism and Signaling | 0.47 | 0.01 | 0.35 | 0.38 | 1.00 | 0.06 | 0.07 | 0.62 |
| Stat3 | 940.99 | Interferon Signaling | 0.66 | 0.01 | 0.47 | 0.26 | 1.00 | 0.06 | 0.08 | 0.82 |
| Nsf | 449.58 | GABAergic Synapse | 1.00 | 0.00 | 0.80 | 0.03 | 1.00 | 0.05 | 0.04 | 1.00 |
| Diaph1 | 345.90 | Cell Migration | 0.60 | 0.00 | 0.46 | 0.01 | 0.79 | 0.06 | 0.02 | 1.00 |
| Abca1 | 3323.12 | Lipid Metabolism | 0.80 | 0.00 | 0.86 | 0.01 | 1.00 | 0.01 | 0.00 | 0.99 |
| Jak2 | 280.11 | JAK-STAT | 0.27 | 0.00 | 0.26 | 0.04 | 1.00 | 0.00 | 0.00 | 0.39 |
| Vegfa | 222.71 | Blood Brain Barrier | 0.04 | 0.00 | 0.69 | 0.00 | 0.49 | 0.00 | 0.00 | 0.24 |
| B4galt5 | 142.66 | Oligodendrocyte Differentiation | 0.03 | 0.00 | 0.11 | 0.01 | 0.95 | 0.00 | 0.00 | 0.06 |
| Olfr110 | 205.80 | Microglia Markers | 0.21 | 0.00 | 0.52 | 0.01 | 0.52 | 0.02 | 0.00 | 0.96 |
| Gsn | 186.35 | Oligodendrocyte Function | 0.03 | 0.00 | 0.95 | 0.00 | 0.31 | 0.00 | 0.00 | 0.34 |
| Pik3cb | 705.03 | MAPK & PI3K | 0.74 | 0.00 | 0.78 | 0.04 | 0.97 | 0.06 | 0.03 | 1.00 |
| Ranbp9 | 598.95 | Neurogenesis | 0.36 | 0.01 | 0.86 | 0.13 | 0.94 | 0.09 | 0.04 | 0.85 |
| Gnb1 | 1038.44 | Serotonergic Synapse | 0.69 | 0.01 | 0.92 | 0.13 | 1.00 | 0.11 | 0.08 | 0.98 |
| Cltc | 6791.66 | Wnt Signaling | 0.98 | 0.03 | 0.95 | 0.36 | 1.00 | 0.32 | 0.35 | 1.00 |
| Psma5 | 446.22 | Antigen Processing and Presentation | 0.23 | 0.03 | 0.84 | 0.26 | 0.87 | 0.37 | 0.78 | 0.72 |
| Arl6ip5 | 828.66 | Glutamatergic Synapse | 0.40 | 0.01 | 0.76 | 0.22 | 0.83 | 0.12 | 0.37 | 0.98 |
| Atg5 | 506.13 | Autophagy | 0.13 | 0.00 | 0.36 | 0.02 | 0.31 | 0.00 | 0.04 | 0.96 |
| Serpine2 | 3622.13 | Microglia-DAM Stage 2-MGnD | 0.02 | 0.00 | 0.43 | 0.08 | 0.11 | 0.01 | 0.50 | 0.50 |
| Prkar1a | 1647.89 | Insulin Signaling | 0.23 | 0.01 | 0.30 | 0.31 | 0.38 | 0.04 | 0.36 | 1.00 |
| Adar | 228.97 | Interferon Signaling | 0.15 | 0.01 | 0.88 | 0.15 | 0.63 | 0.11 | 0.52 | 0.75 |
| Eloc | 1026.23 | Antigen Processing and Presentation | 0.58 | 0.02 | 0.39 | 0.10 | 0.73 | 0.52 | 0.14 | 0.99 |
| Slc25a4 | 1003.41 | Calcium Signaling | 0.69 | 0.03 | 0.20 | 0.09 | 0.60 | 0.85 | 0.23 | 0.90 |
| Ndufb5 | 259.56 | Cannabinoid Signaling | 0.24 | 0.00 | 0.02 | 0.00 | 0.07 | 0.64 | 0.01 | 0.67 |
| Sugt1 | 633.40 | Inflammasome | 0.22 | 0.01 | 0.10 | 0.03 | 0.20 | 0.77 | 0.05 | 0.98 |
| Acot13 | 327.90 | Lipid Metabolism | 0.88 | 0.01 | 0.67 | 0.12 | 1.00 | 0.28 | 0.22 | 0.97 |
| Atp5pb | 702.68 | Glucose Metabolism | 0.82 | 0.00 | 0.22 | 0.02 | 0.70 | 0.24 | 0.05 | 0.87 |
| Atp6v1e1 | 610.66 | Phagocytosis | 0.72 | 0.01 | 0.23 | 0.03 | 0.91 | 0.41 | 0.18 | 0.66 |
| Sod1 | 1529.52 | Oxidative & Nitrosative Stress | 0.32 | 0.00 | 0.07 | 0.00 | 0.19 | 0.11 | 0.01 | 0.87 |
| Ndufa10 | 422.29 | Cannabinoid Signaling | 0.25 | 0.01 | 0.17 | 0.04 | 1.00 | 0.61 | 0.51 | 0.29 |
| Ctss | 54564.60 | Microglia-Homeostatic | 0.05 | 0.00 | 0.27 | 0.01 | 0.86 | 0.14 | 0.39 | 0.14 |
| Aif1 | 6414.13 | Microglia-M1 | 0.58 | 0.04 | 0.29 | 0.14 | 0.98 | 0.80 | 0.59 | 0.64 |
| Tmsb4x | 94436.75 | Microglia-Homeostatic | 0.66 | 0.01 | 0.28 | 0.05 | 0.96 | 0.45 | 0.25 | 0.67 |
| Clta | 4982.53 | Wnt Signaling | 0.49 | 0.01 | 0.11 | 0.02 | 0.87 | 0.64 | 0.27 | 0.35 |
| Abcc8 | 52.36 | Neuronal Markers | 0.04 | 0.00 | 0.03 | 0.00 | 1.00 | 0.54 | 0.47 | 0.03 |
| Cpne2 | 81.83 | Astrocyte Markers | 0.57 | 0.00 | 0.12 | 0.01 | 0.40 | 0.28 | 0.03 | 0.84 |
| Atp5b | 1137.73 | Glucose Metabolism | 0.74 | 0.02 | 0.63 | 0.15 | 0.93 | 0.39 | 0.18 | 1.00 |
| Nlrp3 | 1241.49 | Inflammasome | 0.11 | 0.00 | 0.19 | 0.09 | 0.99 | 0.01 | 0.00 | 0.18 |
| Panx1 | 73.20 | Inflammasome | 0.08 | 0.00 | 0.78 | 0.05 | 0.63 | 0.03 | 0.01 | 0.41 |
| Sqstm1 | 2394.81 | Autophagy | 0.57 | 0.00 | 0.24 | 0.11 | 0.96 | 0.01 | 0.02 | 0.58 |
| C3ar1 | 3811.69 | Primed Microglia | 0.69 | 0.00 | 0.53 | 0.06 | 0.87 | 0.02 | 0.05 | 1.00 |
| Gas6 | 1285.50 | Neurogenesis | 0.92 | 0.00 | 0.82 | 0.11 | 1.00 | 0.07 | 0.08 | 0.99 |
| C9orf72 | 466.04 | Primed Microglia | 0.65 | 0.00 | 0.61 | 0.02 | 1.00 | 0.01 | 0.01 | 0.89 |
| Calm3 | 551.91 | Calcium Signaling | 0.16 | 0.00 | 0.50 | 0.02 | 0.44 | 0.06 | 0.01 | 0.93 |
| Cd244a | 59.86 | Adaptive Immune Response | 0.95 | 0.00 | 0.33 | 0.01 | 0.86 | 0.05 | 0.02 | 0.90 |
| Eid1 | 540.40 | Primed Microglia | 0.90 | 0.00 | 0.32 | 0.00 | 0.91 | 0.01 | 0.01 | 0.83 |
| Gabarapl1 | 598.34 | GABAergic Synapse | 0.11 | 0.00 | 0.15 | 0.00 | 0.15 | 0.02 | 0.00 | 1.00 |
| Gnai1 | 67.21 | Serotonergic Synapse | 0.06 | 0.00 | 0.16 | 0.00 | 0.11 | 0.04 | 0.00 | 0.97 |
| Flt1 | 155.42 | Endothelial Cell Function | 0.38 | 0.01 | 0.08 | 0.02 | 0.23 | 0.80 | 0.07 | 0.86 |
| Sptan1 | 333.20 | Neurogenesis | 0.02 | 0.00 | 0.01 | 0.00 | 0.01 | 0.12 | 0.00 | 0.98 |
| App | 2199.20 | Serotonergic Synapse | 0.10 | 0.00 | 0.03 | 0.00 | 0.05 | 0.05 | 0.00 | 0.94 |
| Ms4a4a | 214.00 | Microglia Markers | 0.13 | 0.01 | 0.97 | 0.11 | 0.64 | 0.12 | 0.02 | 0.66 |
| Il10 | 76.02 | Microglia-M2 | 0.22 | 0.01 | 0.77 | 0.08 | 0.67 | 0.16 | 0.03 | 0.89 |
| Map1lc3b | 621.03 | Autophagy | 0.68 | 0.00 | 0.60 | 0.04 | 0.90 | 0.11 | 0.04 | 1.00 |
| Becn1 | 314.86 | Autophagy | 0.57 | 0.01 | 0.90 | 0.09 | 0.95 | 0.12 | 0.06 | 0.99 |
| Gfap | 138.41 | Astrocyte Markers | 0.90 | 0.00 | 0.84 | 0.02 | 0.99 | 0.03 | 0.02 | 1.00 |
| Ywhaq | 1291.06 | Cell Cycle | 0.27 | 0.00 | 0.51 | 0.01 | 0.57 | 0.02 | 0.00 | 0.98 |
| C4a/b | 269.28 | Astrocyte Markers | 0.88 | 0.00 | 0.46 | 0.00 | 0.91 | 0.00 | 0.00 | 0.97 |
| Epb41l3 | 161.17 | Cytoskeletal Dynamics | 0.37 | 0.00 | 0.43 | 0.00 | 0.61 | 0.00 | 0.00 | 1.00 |
| Arpc1a | 576.49 | Phagocytosis | 0.50 | 0.00 | 0.93 | 0.00 | 0.94 | 0.00 | 0.00 | 0.97 |
| Bax | 192.11 | Apoptosis | 0.40 | 0.00 | 0.21 | 0.14 | 0.43 | 0.01 | 0.09 | 0.99 |
| Cacna2d3 | 67.73 | Ion Transport | 0.30 | 0.00 | 0.51 | 0.01 | 0.60 | 0.00 | 0.02 | 0.99 |
| Stat1 | 480.60 | Interferon Signaling | 0.71 | 0.00 | 0.47 | 0.09 | 0.85 | 0.02 | 0.06 | 0.99 |
| Scn8a | 95.43 | Myelogenesis | 0.92 | 0.00 | 0.38 | 0.07 | 0.94 | 0.01 | 0.02 | 0.88 |
| H2-T23 | 883.41 | Antigen Processing and Presentation | 0.44 | 0.00 | 0.81 | 0.02 | 0.87 | 0.01 | 0.03 | 0.98 |
| B2m | 35445.12 | Microglia-DAM Stage 1 | 0.24 | 0.00 | 0.37 | 0.00 | 1.00 | 0.00 | 0.01 | 0.45 |
| Tyrobp | 17491.26 | Microglia-DAM Stage 1 | 0.87 | 0.00 | 0.49 | 0.01 | 0.98 | 0.03 | 0.02 | 0.92 |
| C1qa | 36650.02 | Microglia-Homeostatic | 0.61 | 0.00 | 0.60 | 0.01 | 1.00 | 0.03 | 0.03 | 0.87 |
| C1qb | 27676.80 | Microglia-Homeostatic | 0.64 | 0.00 | 0.58 | 0.02 | 1.00 | 0.08 | 0.07 | 0.87 |
| Ppt1 | 1931.50 | Lipid Metabolism | 0.42 | 0.00 | 0.39 | 0.01 | 1.00 | 0.08 | 0.07 | 0.62 |
| Atp6v1g1 | 530.88 | Phagocytosis | 0.79 | 0.00 | 0.75 | 0.02 | 0.97 | 0.03 | 0.02 | 1.00 |
| Tspo | 234.96 | Microglia-M1 | 0.49 | 0.00 | 0.15 | 0.00 | 0.42 | 0.04 | 0.00 | 0.93 |
| Fcgr4 | 296.67 | Phagocytosis | 0.42 | 0.00 | 0.06 | 0.00 | 0.78 | 0.02 | 0.01 | 0.22 |
| Lamp2 | 4362.37 | Autophagy | 0.09 | 0.00 | 0.61 | 0.00 | 0.77 | 0.00 | 0.01 | 0.37 |
| Grn | 5765.84 | Microglia-DAM Stage 1 | 0.05 | 0.00 | 0.95 | 0.00 | 0.41 | 0.00 | 0.01 | 0.46 |
| Ctsl | 1985.32 | Microglia-DAM Stage 2-MGnD | 0.03 | 0.00 | 0.54 | 0.00 | 0.51 | 0.00 | 0.00 | 0.17 |
| Cd84 | 2961.69 | Primed Microglia | 0.02 | 0.00 | 0.67 | 0.00 | 0.41 | 0.00 | 0.00 | 0.18 |
| Srgn | 4008.85 | A1 Astrocyte | 0.36 | 0.00 | 0.59 | 0.00 | 0.71 | 0.01 | 0.00 | 0.99 |
| Cd14 | 6633.99 | A2 Astrocyte | 0.91 | 0.00 | 0.51 | 0.00 | 0.98 | 0.00 | 0.00 | 0.94 |
| Il1b | 7758.06 | Microglia-M1 | 0.77 | 0.00 | 0.78 | 0.00 | 0.97 | 0.00 | 0.00 | 1.00 |
| Csf1 | 1774.01 | Microglia-DAM Stage 2-MGnD | 0.81 | 0.00 | 0.53 | 0.00 | 0.91 | 0.00 | 0.00 | 0.99 |
| Tlr2 | 1855.79 | Primed Microglia | 0.24 | 0.00 | 0.55 | 0.00 | 0.97 | 0.00 | 0.00 | 0.57 |
| H2-D1 | 4509.76 | Antigen Processing and Presentation | 0.75 | 0.00 | 0.61 | 0.00 | 1.00 | 0.00 | 0.00 | 0.93 |
| Lamp1 | 4285.20 | Autophagy | 0.07 | 0.00 | 0.80 | 0.00 | 0.59 | 0.00 | 0.00 | 0.40 |
| Ctsd | 98930.23 | Microglia-DAM Stage 1 | 0.52 | 0.00 | 0.70 | 0.00 | 1.00 | 0.00 | 0.00 | 0.87 |
| Timp2 | 6859.80 | Microglia-DAM Stage 1 | 0.25 | 0.00 | 0.56 | 0.00 | 0.97 | 0.00 | 0.00 | 0.58 |
| Ctsb | 12402.21 | Microglia-DAM Stage 1 | 0.13 | 0.00 | 0.22 | 0.00 | 0.99 | 0.00 | 0.00 | 0.23 |
| Gla | 195.47 | NO Metabolism and Signaling | 0.35 | 0.00 | 0.21 | 0.00 | 0.99 | 0.00 | 0.00 | 0.40 |
| Fabp5 | 125.74 | Microglia-DAM Stage 1 | 0.91 | 0.00 | 0.93 | 0.00 | 1.00 | 0.00 | 0.00 | 1.00 |
| Cst7 | 1880.34 | Microglia-DAM Stage 2-MGnD | 0.63 | 0.00 | 0.61 | 0.00 | 1.00 | 0.00 | 0.00 | 0.88 |
| Spp1 | 6373.26 | Microglia-DAM Stage 2-MGnD | 0.92 | 0.00 | 0.88 | 0.00 | 1.00 | 0.00 | 0.00 | 1.00 |
| Cd68 | 7056.76 | Microglia-DAM Stage 2-MGnD | 0.84 | 0.00 | 0.38 | 0.00 | 0.85 | 0.00 | 0.00 | 0.95 |
| Itgax | 263.83 | Microglia-DAM Stage 2-MGnD | 0.68 | 0.00 | 0.75 | 0.00 | 1.00 | 0.00 | 0.00 | 0.95 |
| Lpl | 570.95 | Microglia-DAM Stage 2-MGnD | 0.60 | 0.00 | 0.60 | 0.00 | 1.00 | 0.00 | 0.00 | 0.87 |
| Apoe | 39490.38 | Microglia-DAM Stage 1 | 0.98 | 0.00 | 0.80 | 0.00 | 1.00 | 0.00 | 0.00 | 1.00 |
| Axl | 941.91 | Microglia-DAM Stage 2-MGnD | 0.90 | 0.00 | 0.78 | 0.00 | 1.00 | 0.00 | 0.00 | 0.99 |
| Clec7a | 2266.04 | Microglia-DAM Stage 2-MGnD | 0.83 | 0.00 | 0.61 | 0.00 | 1.00 | 0.00 | 0.00 | 0.95 |
| Lgals3 | 1566.33 | Microglia-DAM Stage 2-MGnD | 0.87 | 0.00 | 0.43 | 0.00 | 0.96 | 0.00 | 0.00 | 0.89 |
| Ptger4 | 197.48 | Neuroactive Ligands and Receptors | 0.50 | 0.00 | 0.95 | 0.00 | 0.95 | 0.00 | 0.00 | 0.97 |
| Igf1 | 85.18 | Microglia-M2 | 0.67 | 0.00 | 0.60 | 0.00 | 0.90 | 0.00 | 0.00 | 1.00 |
| Fth1 | 50481.79 | Microglia-DAM Stage 1 | 0.89 | 0.00 | 0.50 | 0.00 | 0.98 | 0.00 | 0.00 | 0.93 |
| Lyz2 | 7058.21 | Microglia-DAM Stage 1 | 0.82 | 0.00 | 0.39 | 0.00 | 0.96 | 0.00 | 0.00 | 0.85 |
| Cybb | 901.48 | Microglia-M1 | 0.87 | 0.00 | 0.34 | 0.00 | 0.84 | 0.00 | 0.00 | 0.93 |
| Atp6v1c1 | 1262.17 | Primed Microglia | 0.22 | 0.00 | 0.10 | 0.00 | 0.19 | 0.00 | 0.00 | 0.98 |
| Ass1 | 84.87 | NO Metabolism and Signaling | 0.24 | 0.00 | 0.82 | 0.00 | 0.88 | 0.00 | 0.00 | 0.71 |
| Ptprc | 1208.19 | Microglia-M1 | 0.55 | 0.00 | 0.90 | 0.00 | 0.98 | 0.00 | 0.00 | 0.95 |
| Got1 | 511.97 | Glutamatergic Synapse | 0.19 | 0.00 | 0.12 | 0.00 | 1.00 | 0.00 | 0.00 | 0.19 |
| Gpnmb | 2840.49 | Microglia-DAM Stage 1 | 0.40 | 0.00 | 0.43 | 0.00 | 1.00 | 0.00 | 0.00 | 0.63 |
| Msr1 | 84.55 | Microglia-DAM Stage 2-MGnD | 0.06 | 0.00 | 0.15 | 0.00 | 0.97 | 0.00 | 0.00 | 0.10 |
| C3 | 243.27 | Primed Microglia | 0.07 | 0.00 | 0.31 | 0.00 | 0.88 | 0.00 | 0.00 | 0.19 |
| Pgk1 | 1433.26 | Glucose Metabolism | 0.07 | 0.00 | 0.04 | 0.00 | 0.05 | 0.00 | 0.00 | 0.99 |
| Lilrb4a | 1285.56 | Microglia-DAM Stage 2-MGnD | 0.11 | 0.00 | 0.50 | 0.00 | 0.86 | 0.00 | 0.00 | 0.35 |
| Myo5a | 302.69 | Myelogenesis | 0.83 | 0.00 | 0.90 | 0.00 | 0.99 | 0.00 | 0.00 | 1.00 |
| Ptgs2 | 496.30 | Microglia-M1 | 0.14 | 0.00 | 0.88 | 0.00 | 0.61 | 0.00 | 0.00 | 0.73 |
| Aplp2 | 1314.62 | Microglia-DAM Stage 1 | 0.28 | 0.00 | 0.33 | 0.00 | 0.45 | 0.00 | 0.00 | 1.00 |
| Hspa9 | 927.18 | Primed Microglia | 0.22 | 0.00 | 0.81 | 0.00 | 0.69 | 0.00 | 0.00 | 0.87 |
| Vim | 943.56 | Macrophage Function | 0.31 | 0.00 | 0.33 | 0.00 | 0.48 | 0.00 | 0.00 | 1.00 |
| Bhlhe40 | 343.24 | Neuronal Markers | 0.62 | 0.00 | 0.31 | 0.00 | 0.68 | 0.00 | 0.00 | 0.98 |
| Ctnnb1 | 2100.69 | Wnt Signaling | 0.47 | 0.00 | 0.17 | 0.00 | 0.42 | 0.00 | 0.00 | 0.95 |
| Ifnar1 | 735.89 | Interferon Signaling | 0.96 | 0.00 | 1.00 | 0.00 | 1.00 | 0.00 | 0.00 | 1.00 |
| Ank | 452.29 | Primed Microglia | 0.89 | 0.00 | 0.72 | 0.00 | 0.98 | 0.00 | 0.00 | 1.00 |
| Atp6v1a | 634.69 | Primed Microglia | 0.64 | 0.00 | 0.51 | 0.00 | 0.84 | 0.00 | 0.00 | 1.00 |
| Gas7 | 107.64 | Primed Microglia | 0.66 | 0.00 | 0.53 | 0.00 | 0.86 | 0.00 | 0.00 | 1.00 |
| Txn1 | 1284.32 | Oxidative & Nitrosative Stress | 0.43 | 0.00 | 0.51 | 0.00 | 0.71 | 0.00 | 0.00 | 1.00 |
| Atp6v1b2 | 801.94 | Primed Microglia | 0.60 | 0.00 | 0.38 | 0.00 | 0.99 | 0.00 | 0.00 | 0.73 |
| Cspg4 | 111.95 | Neurogenesis | 0.55 | 0.00 | 0.89 | 0.00 | 0.99 | 0.00 | 0.00 | 0.94 |
| Atp6v0d1 | 893.27 | Phagocytosis | 0.86 | 0.00 | 0.77 | 0.00 | 0.99 | 0.00 | 0.00 | 1.00 |
| Galc | 320.65 | Primed Microglia | 0.32 | 0.00 | 0.20 | 0.00 | 1.00 | 0.00 | 0.00 | 0.37 |
| Psmb8 | 894.83 | Primed Microglia | 0.88 | 0.00 | 0.57 | 0.00 | 0.95 | 0.00 | 0.00 | 0.99 |
| Grb2 | 948.94 | T-cell signaling | 0.10 | 0.00 | 0.40 | 0.00 | 0.91 | 0.00 | 0.00 | 0.29 |
| Gnas | 1864.08 | Serotonergic Synapse | 0.64 | 0.00 | 0.62 | 0.01 | 1.00 | 0.00 | 0.00 | 0.89 |
| Hacd2 | 225.77 | Lipid Metabolism | 0.63 | 0.00 | 0.77 | 0.00 | 0.94 | 0.00 | 0.00 | 1.00 |
| Atp1a1 | 1181.52 | Primed Microglia | 0.94 | 0.00 | 0.19 | 0.00 | 0.78 | 0.02 | 0.01 | 0.72 |
| Meaf6 | 178.59 | Neurogenesis | 0.98 | 0.00 | 0.93 | 0.04 | 1.00 | 0.05 | 0.04 | 1.00 |
| Atp6v1d | 492.55 | Phagocytosis | 0.24 | 0.00 | 0.25 | 0.00 | 0.36 | 0.02 | 0.00 | 1.00 |
| Cd74 | 4450.02 | Microglia-M1 | 0.69 | 0.01 | 0.41 | 0.07 | 0.80 | 0.36 | 0.10 | 0.99 |
| Por | 249.47 | NO Metabolism and Signaling | 0.52 | 0.00 | 0.16 | 0.01 | 0.92 | 0.24 | 0.10 | 0.44 |
| Tomm20 | 849.69 | Autophagy | 0.88 | 0.00 | 0.97 | 0.04 | 1.00 | 0.04 | 0.03 | 1.00 |
| Pgam1 | 552.02 | Glucose Metabolism | 0.90 | 0.00 | 0.63 | 0.03 | 0.97 | 0.07 | 0.04 | 0.99 |
| Alas1 | 376.40 | Primed Microglia | 0.75 | 0.00 | 0.69 | 0.03 | 1.00 | 0.01 | 0.01 | 0.95 |
| Ccl6 | 612.15 | Microglia-DAM Stage 2-MGnD | 0.73 | 0.01 | 0.84 | 0.13 | 0.98 | 0.20 | 0.11 | 1.00 |
| Trem2 | 5474.90 | Microglia-DAM Stage 2-MGnD | 0.82 | 0.01 | 0.97 | 0.19 | 1.00 | 0.18 | 0.23 | 1.00 |
| C1qc | 28034.06 | Microglia-Homeostatic | 0.63 | 0.02 | 0.92 | 0.21 | 0.99 | 0.25 | 0.36 | 0.97 |
| Syngr1 | 954.10 | Neurogenesis | 0.53 | 0.02 | 0.88 | 0.19 | 0.94 | 0.25 | 0.11 | 0.98 |
| Cd9 | 6707.48 | Microglia-DAM Stage 2-MGnD | 0.03 | 0.00 | 0.87 | 0.00 | 0.27 | 0.00 | 0.04 | 0.37 |
| Tap1 | 424.16 | Primed Microglia | 0.49 | 0.02 | 0.77 | 0.31 | 0.89 | 0.18 | 0.44 | 0.99 |
| Gpr84 | 963.05 | Microglia-M1 | 0.86 | 0.04 | 0.42 | 0.66 | 0.96 | 0.16 | 0.30 | 0.88 |
| Pmp22 | 1740.30 | Myelogenesis | 0.53 | 0.02 | 0.57 | 0.47 | 1.00 | 0.16 | 0.15 | 0.81 |
| Rtn4 | 1824.24 | Neurogenesis | 0.68 | 0.01 | 0.99 | 0.11 | 0.99 | 0.11 | 0.07 | 0.99 |
| Csf3r | 1682.53 | Cytokines | 0.00 | 0.00 | 0.53 | 0.03 | 0.12 | 0.01 | 0.00 | 0.03 |
| Hmox1 | 902.84 | Primed Microglia | 0.04 | 0.00 | 0.78 | 0.07 | 0.48 | 0.04 | 0.01 | 0.30 |
| Hsp90ab1 | 3385.72 | Proteotoxic Stress | 0.02 | 0.00 | 0.12 | 0.01 | 0.05 | 0.27 | 0.00 | 0.83 |
| Stat2 | 139.11 | Interferon Signaling | 0.08 | 0.02 | 0.02 | 1.00 | 0.91 | 0.02 | 0.04 | 0.03 |
| Adam22 | 53.84 | Myelogenesis | 0.34 | 0.01 | 0.18 | 0.55 | 0.99 | 0.04 | 0.06 | 0.36 |
| Abi2 | 125.92 | Neurogenesis | 0.44 | 0.03 | 0.64 | 0.47 | 1.00 | 0.20 | 0.15 | 0.80 |
| Gsk3b | 479.12 | Wnt Signaling | 0.64 | 0.02 | 0.77 | 0.32 | 1.00 | 0.18 | 0.15 | 0.94 |
| Dnaja2 | 281.93 | Proteotoxic Stress | 0.25 | 0.02 | 0.51 | 0.43 | 0.98 | 0.12 | 0.07 | 0.55 |
| Tgfb1 | 2776.22 | Microglia-Homeostatic | 0.09 | 0.00 | 0.00 | 1.00 | 0.01 | 0.00 | 0.55 | 0.43 |
| Arpc2 | 2046.01 | Phagocytosis | 0.29 | 0.02 | 0.11 | 0.86 | 0.25 | 0.04 | 0.62 | 0.97 |
| Ywhag | 722.72 | Neurogenesis | 0.84 | 0.01 | 0.10 | 0.60 | 0.50 | 0.02 | 0.14 | 0.66 |

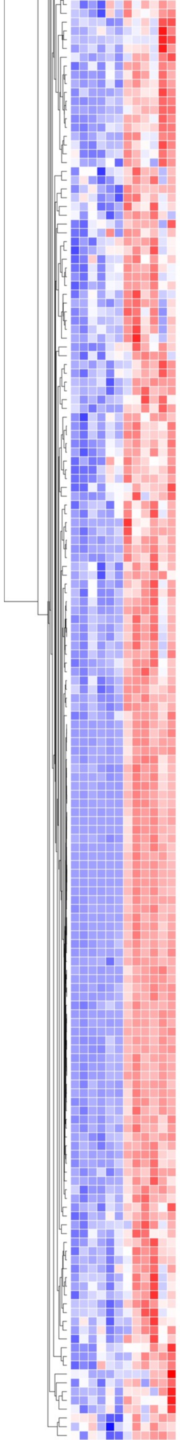

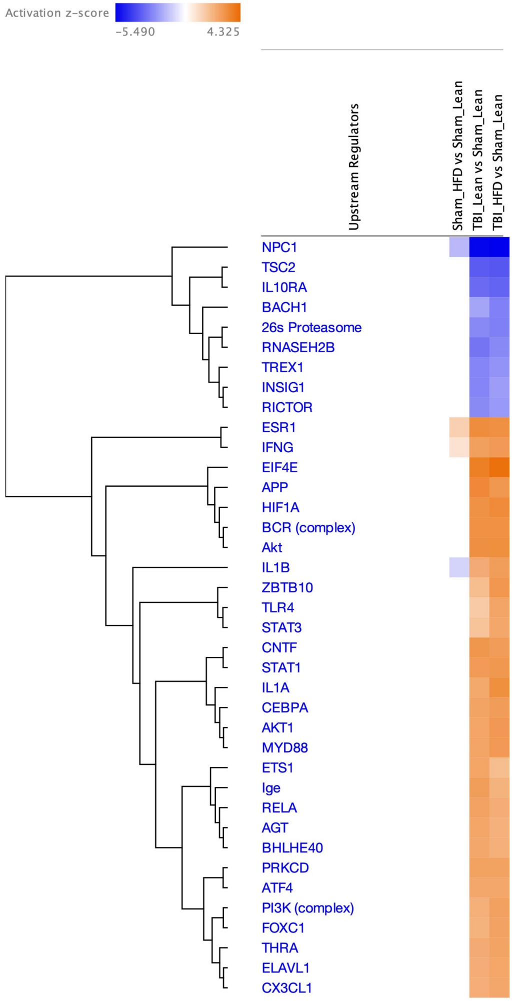

#### Upstream regulators – Genes (upregulated)

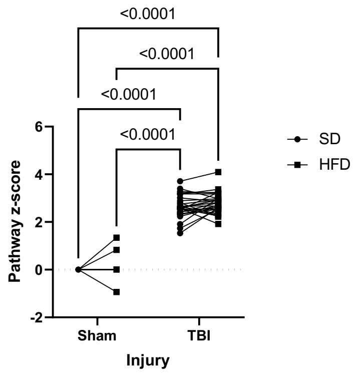

| | | | | | |
|---|---|---|---|---|---|
| Source of Variation | % of total variation | P value | P value summary | Significant? | |
| Injury | 93.23 | <0.0001 | **** | Yes | |
| Diet | 0.1260 | 0.0477 | * | Yes | |
| Injury x Diet | 0.04093 | 0.3049 | ns | No | |
| Pathways x Injury | 1.961 | | | | |
| Pathways x Diet | 0.8223 | | | | |
| Pathways | 2.771 | | | | |
| ANOVA table | SS | DF | MS | F (DFn, DFd) | P value |
| Injury | 210.5 | 1 | 210.5 | F (1, 28) = 1331 | P<0.0001 |
| Diet | 0.2844 | 1 | 0.2844 | F (1, 28) = 4.290 | P=0.0477 |
| Injury x Diet | 0.09241 | 1 | 0.09241 | F (1, 28) = 1.092 | P=0.3049 |

#### Upstream regulators – Genes (downregulated)

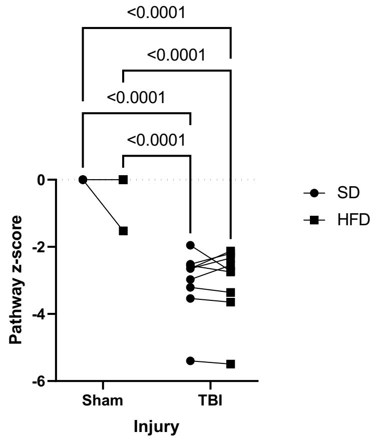

| | | | | | |
|---|---|---|---|---|---|
| Source of Variation | % of total variation | P value | P value summary | Significant? | |
| Injury | 80.49 | <0.0001 | **** | Yes | |
| Diet | 0.04470 | 0.5596 | ns | No | |
| Injury x Diet | 0.09182 | 0.3585 | ns | No | |
| Pathways x Injury | 5.046 | | | | |
| Pathways x Diet | 0.9651 | | | | |
| Pathways | 12.59 | | | | |
| ANOVA table | SS | DF | MS | F (DFn, DFd) | P value |
| Injury | 78.28 | 1 | 78.28 | F (1, 8) = 127.6 | P<0.0001 |
| Diet | 0.04347 | 1 | 0.04347 | F (1, 8) = 0.3705 | P=0.5596 |
| Injury x Diet | 0.08930 | 1 | 0.08930 | F (1, 8) = 0.9491 | P=0.3585 |

**Figure 2-Supplemental**

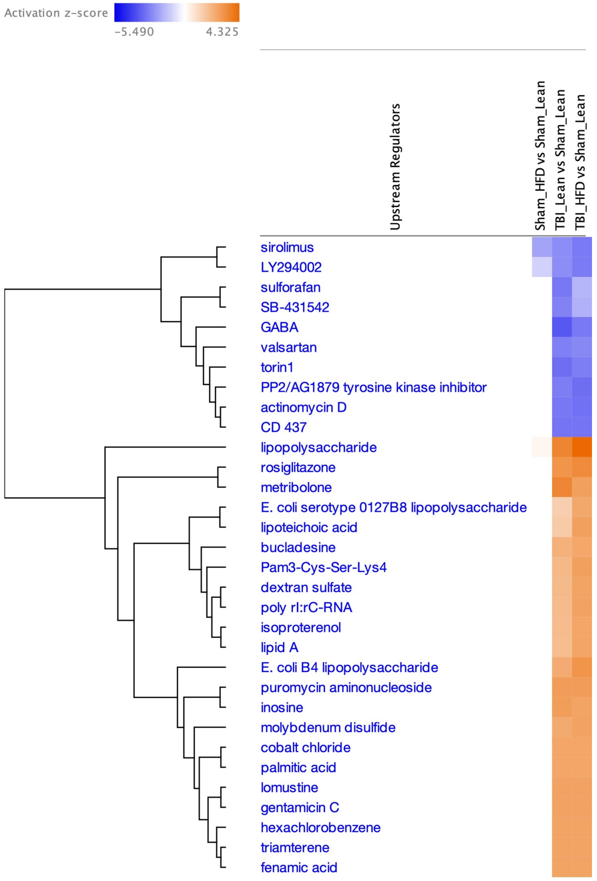

#### Upstream regulators – Drugs (upregulated)

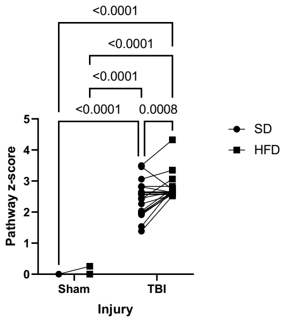

| | | | | | |
|---|---|---|---|---|---|
| Source of Variation | % of total variation | P value | P value summary | Significant? | |
| Injury | 93.38 | <0.0001 | **** | Yes | |
| Diet | 0.3788 | 0.0043 | ** | Yes | |
| Injury x Diet | 0.3277 | 0.0049 | ** | Yes | |
| Pathways x Injury | 2.013 | | | | |
| Pathways x Diet | 0.7758 | | | | |
| Pathways | 2.429 | | | | |
| ANOVA table | SS | DF | MS | F (DFn, DFd) | P value |
| Injury | 149.1 | 1 | 149.1 | F (1, 21) = 974.0 | P<0.0001 |
| Diet | 0.6051 | 1 | 0.6051 | F (1, 21) = 10.25 | P=0.0043 |
| Injury x Diet | 0.5234 | 1 | 0.5234 | F (1, 21) = 9.909 | P=0.0049 |

#### Upstream regulators – Drugs (downregulated)

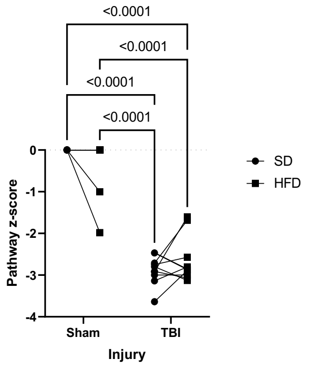

| | | | | | |
|---|---|---|---|---|---|
| Source of Variation | % of total variation | P value | P value summary | Significant? | |
| Diet | 88.90 | <0.0001 | **** | Yes | |
| Injury | 0.01425 | 0.8553 | ns | No | |
| Diet x Injury | 0.9141 | 0.0278 | * | Yes | |
| Pathway x Diet | 3.051 | | | | |
| Pathway x Injury | 3.643 | | | | |
| Pathway | 2.283 | | | | |
| ANOVA table | SS | DF | MS | F (DFn, DFd) | P value |
| Diet | 68.35 | 1 | 68.35 | F (1, 9) = 262.2 | P<0.0001 |
| Injury | 0.01096 | 1 | 0.01096 | F (1, 9) = 0.03521 | P=0.8553 |
| Diet x Injury | 0.7028 | 1 | 0.7028 | F (1, 9) = 6.860 | P=0.0278 |

**Figure 3-Supplemental**

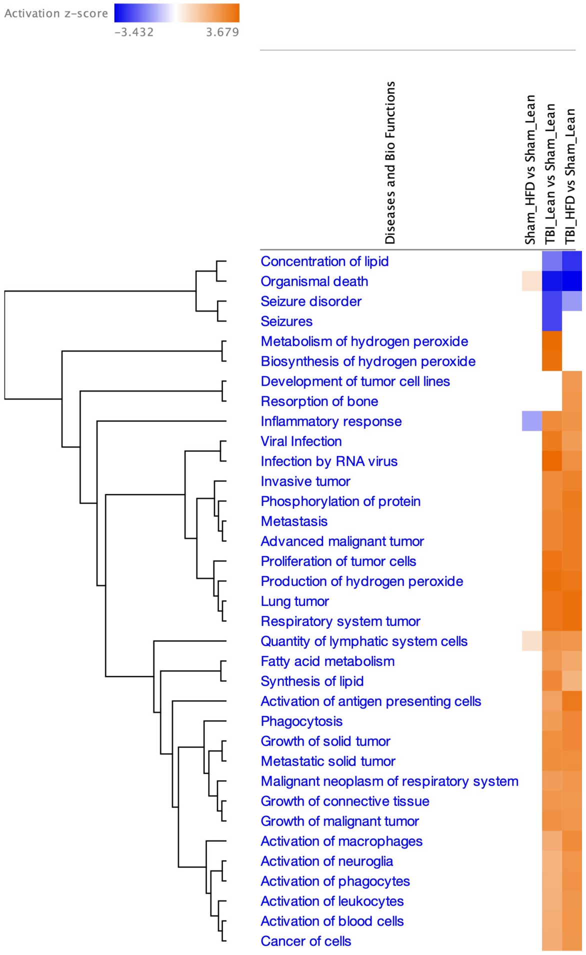

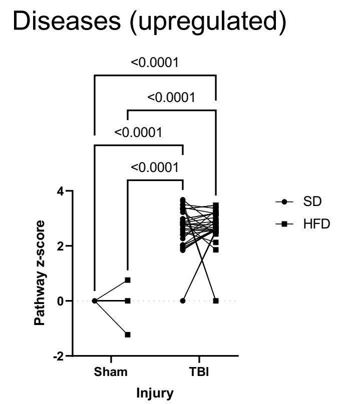

| | | | | | |
|---|---|---|---|---|---|
| Source of Variation | % of total variation | P value | P value summary | Significant? | |
| Injury | 82.76 | <0.0001 | **** | Yes | |
| Diet | 0.001747 | 0.9169 | ns | No | |
| Injury x Diet | 0.008928 | 0.8121 | ns | No | |
| Pathway x Injury | 3.984 | | | | |
| Pathway x Diet | 4.737 | | | | |
| Pathway | 3.853 | | | | |
| ANOVA table | SS | DF | MS | F (DFn, DFd) | P value |
| Injury | 209.7 | 1 | 209.7 | F (1, 30) = 623.2 | P<0.0001 |
| Diet | 0.004428 | 1 | 0.004428 | F (1, 30) = 0.01107 | P=0.9169 |
| Injury x Diet | 0.02263 | 1 | 0.02263 | F (1, 30) = 0.05752 | P=0.8121 |

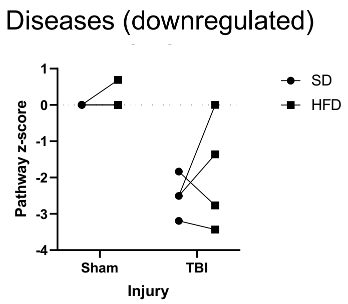

| | | | | | |
|---|---|---|---|---|---|
| Source of Variation | % of total variation | P value | P value summary | Significant? | |
| Diet | 2.092 | 0.3492 | ns | No | |
| Injury | 69.67 | 0.0203 | * | Yes | |
| Diet x Injury | 0.6655 | 0.6327 | ns | No | |
| Pathway x Diet | 5.123 | | | | |
| Pathway x Injury | 10.23 | | | | |
| Pathway | 5.118 | | | | |
| ANOVA table | SS | DF | MS | F (DFn, DFd) | P value |
| Diet | 0.6281 | 1 | 0.6281 | F (1, 3) = 1.225 | P=0.3492 |
| Injury | 20.92 | 1 | 20.92 | F (1, 3) = 20.42 | P=0.0203 |
| Diet x Injury | 0.1998 | 1 | 0.1998 | F (1, 3) = 0.2811 | P=0.6327 |

**Figure 4-Supplemental**

### Figure 5-Supplemental

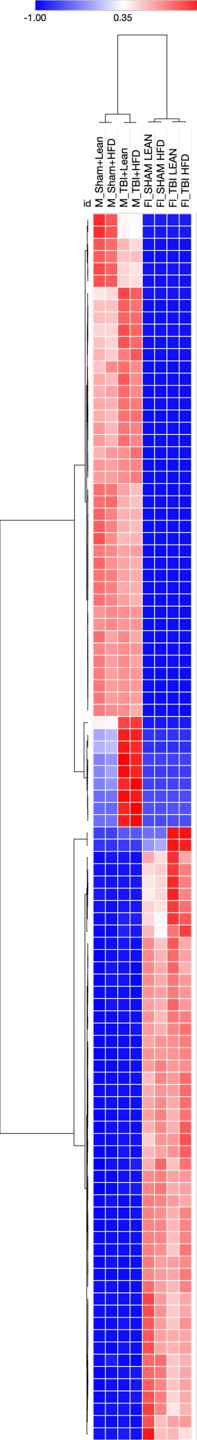

| Symbol | Average | Ratio M/FI | Probe.Annotation |
|---|---|---|---|
| P2ry12 | 9641.56 | 544.96 | Microglia-Homeostatic |
| Slc2a5 | 2233.87 | 340.13 | Microglia-Homeostatic |
| Gpr34 | 4831.00 | 1048.26 | Microglia-Homeostatic |
| Siglech | 3553.03 | 391.04 | Microglia-Homeostatic |
| P2ry13 | 1932.16 | 425.24 | Microglia-Homeostatic |
| Tmem119 | 656.11 | 99.59 | Microglia-Homeostatic |
| Tyrobp | 8770.31 | 354.31 | Microglia-DAM Stage 1 |
| C1qb | 13854.00 | 887.16 | Microglia-Homeostatic |
| Trem2 | 2742.37 | 556.39 | Microglia-DAM Stage 2-MGnD |
| C1qa | 18344.93 | 919.93 | Microglia-Homeostatic |
| Cd84 | 1487.97 | 207.87 | Primed Microglia |
| Gpr84 | 484.80 | 147.18 | Microglia-M1 |
| Ccl3 | 8333.95 | 366.56 | Chemokines |
| Ctss | 27304.59 | 1223.81 | Microglia-Homeostatic |
| Aif1 | 3214.04 | 459.98 | Microglia-M1 |
| C1qc | 14032.68 | 895.84 | Microglia-Homeostatic |
| C3ar1 | 1914.23 | 227.22 | Primed Microglia |
| Ccl12 | 1427.55 | 266.27 | Chemokines |
| Lag3 | 1182.71 | 188.01 | Adaptive Immune Response |
| Csf3r | 846.34 | 165.75 | Cytokines |
| Irf8 | 564.26 | 103.27 | Interferon Signaling |
| Tlr7 | 412.71 | 151.30 | NF-kappaB Signaling |
| Olfml3 | 3070.09 | 189.48 | Microglia-Homeostatic |
| Ccr5 | 1741.90 | 148.85 | Primed Microglia |
| Csf1r | 11666.05 | 565.35 | Microglia-Homeostatic |
| Cx3cr1 | 7056.94 | 857.64 | Microglia-Homeostatic |
| Abcc3 | 304.44 | 128.20 | Oxidative & Nitrosative Stress |
| Ncf1 | 1441.43 | 100.79 | Phagocytosis |
| Il10ra | 1706.53 | 216.99 | Cytokines |
| Fcrls | 5546.36 | 403.60 | Microglia-Homeostatic |
| Itgam | 2638.94 | 345.09 | Microglia-Homeostatic |
| Adora3 | 318.85 | 186.24 | Microglia-Homeostatic |
| Il1a | 2517.05 | 255.54 | Cytokines |
| Tnf | 981.80 | 178.46 | Microglia-M1 |
| Hexb | 38867.30 | 182.76 | Microglia-Homeostatic |
| Spi1 | 1234.10 | 166.60 | Microglia-Homeostatic |
| Fcgr1 | 1495.53 | 363.81 | Phagocytosis |
| Cd33 | 1010.15 | 186.86 | Microglia-Homeostatic |
| Hpgds | 1326.50 | 199.49 | Microglia Markers |
| Lair1 | 569.70 | 99.87 | T-cell signaling |
| Lrrc25 | 275.77 | 98.12 | Macrophage Function |
| Cd68 | 3549.35 | 168.26 | Microglia-DAM Stage 2-MGnD |
| Il1b | 3891.33 | 315.55 | Microglia-M1 |
| Cd14 | 3335.97 | 174.83 | A2 Astrocyte |
| Lyz2 | 3541.15 | 293.01 | Microglia-DAM Stage 1 |
| Cybb | 454.79 | 111.40 | Microglia-M1 |
| Lilrb4a | 648.90 | 105.09 | Microglia-DAM Stage 2-MGnD |
| Clec7a | 1137.57 | 248.75 | Microglia-DAM Stage 2-MGnD |
| Cst7 | 947.06 | 136.49 | Microglia-DAM Stage 2-MGnD |
| Itgax | 133.35 | 91.81 | Microglia-DAM Stage 2-MGnD |
| Gfap | 1068.60 | 0.07 | Astrocyte Markers |
| Klk6 | 432.20 | 0.04 | Myelogenesis |
| Cldn5 | 3668.95 | 0.05 | Blood Brain Barrier |
| Plekhb1 | 4801.19 | 0.02 | Oligodendrocyte Markers |
| Mbp | 4298.43 | 0.03 | Oligodendrocyte Markers |
| Bcas1 | 1282.24 | 0.03 | Oligodendrocyte Function |
| S100b | 707.17 | 0.04 | Astrocyte Markers |
| Aqp4 | 369.88 | 0.06 | Astrocyte Markers |
| Gdpd2 | 34.97 | 0.04 | Astrocyte Markers |
| Itm2a | 2904.69 | 0.05 | Blood Brain Barrier |
| Abcb1a | 1320.47 | 0.08 | Blood Brain Barrier |
| Esam | 750.16 | 0.08 | Blood Brain Barrier |
| Adgrl4 | 642.16 | 0.08 | Blood Brain Barrier |
| Dnm3 | 477.19 | 0.06 | Neurogenesis |
| Lsr | 534.28 | 0.09 | Blood Brain Barrier |
| Pllp | 418.53 | 0.03 | Myelogenesis |
| Gjb1 | 309.99 | 0.03 | Oligodendrocyte Function |
| Cnp | 9934.92 | 0.04 | Oligodendrocyte Function |
| Ermn | 1204.08 | 0.03 | Oligodendrocyte Markers |
| Mog | 1961.68 | 0.02 | Oligodendrocyte Markers |
| Mag | 1411.74 | 0.05 | Oligodendrocyte Differentiation/Maturation |
| Sox10 | 1155.60 | 0.02 | Oligodendrocyte Markers |
| Atp9a | 380.01 | 0.06 | Ion Transport |
| Gal3st1 | 164.38 | 0.07 | Myelogenesis |
| Erbb3 | 109.30 | 0.08 | Oligodendrocyte Function |
| Myrf | 527.23 | 0.04 | Oligodendrocyte Differentiation/Maturation |
| Mapt | 389.96 | 0.07 | Neurogenesis |
| Grm7 | 71.29 | 0.09 | Neuroactive Ligands and Receptors |
| Plp1 | 20801.03 | 0.02 | Oligodendrocyte Markers |
| Gnai1 | 457.76 | 0.08 | Serotonergic Synapse |
| Nckap1 | 634.68 | 0.08 | Neurogenesis |
| Ncam1 | 849.07 | 0.05 | Neuronal Markers |
| Ugt8a | 580.17 | 0.03 | Oligodendrocyte Markers |
| Fa2h | 325.91 | 0.04 | Oligodendrocyte Differentiation/Maturation |
| Gpr17 | 260.18 | 0.03 | Purinergic Signaling |
| Sox2 | 461.57 | 0.06 | Neurogenesis |
| Olig1 | 375.47 | 0.02 | Oligodendrocyte Differentiation/Maturation |
| Adcyap1r1 | 252.13 | 0.07 | Neurotrophin Signaling |
| Acsl3 | 884.51 | 0.05 | Lipid Metabolism |
| Slc6a1 | 636.18 | 0.03 | Astrocyte Markers |
| Pdgfra | 304.10 | 0.06 | MAPK & PI3K |
| Ptprz1 | 1014.24 | 0.03 | Oligodendrocyte Differentiation/Maturation |
| Nrxn1 | 155.16 | 0.08 | Neurogenesis |
| Gpm6a | 2111.07 | 0.06 | Neurogenesis |
| Glrb | 500.82 | 0.04 | Neuroactive Ligands and Receptors |
| Amph | 113.61 | 0.07 | Phagocytosis |
| Olig2 | 98.76 | 0.09 | Oligodendrocyte Differentiation/Maturation |
| Myorg | 168.51 | 0.07 | Astrocyte Markers |
| Ddah1 | 263.02 | 0.05 | NO Metabolism and Signaling |
| Npas2 | 67.30 | 0.07 | Neuronal Markers |

### Figure 6-Supplemental

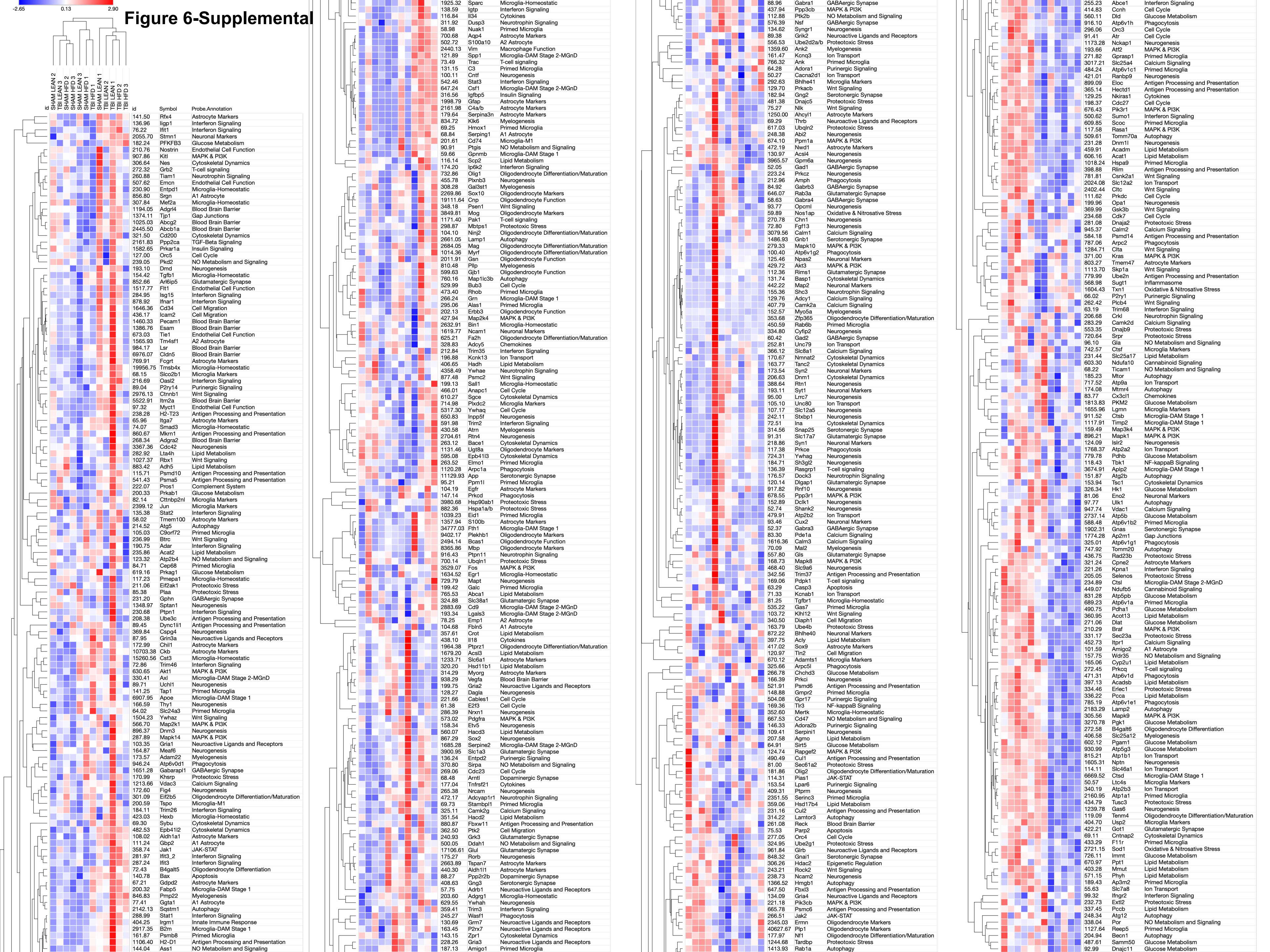

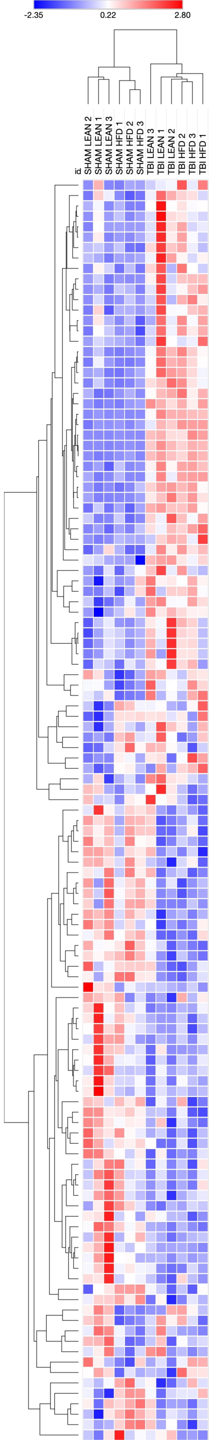

| Symbol | Average | Probe.Annotation | p_val_Diet | p_val_Condition | p_val_diet:condition | p_TBILean:ShamLean | p_TBIHFD:ShamLean | p_TBIHFD:ShamHFD | p_ShamHFD:ShamLean | p_TBIHFD:TBILean |
|---|---|---|---|---|---|---|---|---|---|---|
| Egfr | 104.19 | Astrocyte Markers | 0.46 | 0.04 | 0.09 | 0.97 | 0.17 | 0.06 | 0.85 | 0.30 |
| Aldh1a1 | 108.02 | Astrocyte Markers | 0.06 | 0.01 | 0.57 | 0.28 | 0.84 | 0.09 | 0.28 | 0.69 |
| H2-D1 | 1106.40 | Antigen Processing and Presentation | 0.23 | 0.02 | 0.74 | 0.16 | 0.62 | 0.30 | 0.91 | 0.67 |
| B2m | 2917.35 | Microglia-DAM Stage 1 | 0.25 | 0.04 | 0.46 | 0.20 | 0.85 | 0.68 | 0.99 | 0.52 |
| Irgm1 | 404.25 | Innate Immune Response | 0.38 | 0.03 | 0.68 | 0.20 | 0.61 | 0.42 | 0.98 | 0.77 |
| Stat1 | 288.99 | Interferon Signaling | 0.39 | 0.03 | 0.81 | 0.26 | 0.66 | 0.41 | 0.96 | 0.84 |
| Igtp | 138.59 | Interferon Signaling | 0.10 | 0.00 | 0.48 | 0.05 | 0.54 | 0.20 | 0.85 | 0.32 |
| Sparc | 1925.32 | Microglia-Homeostatic | 0.18 | 0.00 | 0.17 | 0.02 | 0.39 | 0.40 | 1.00 | 0.23 |
| Jak1 | 358.74 | JAK-STAT | 0.45 | 0.04 | 0.66 | 0.56 | 0.69 | 0.26 | 0.81 | 0.99 |
| Ggta1 | 77.41 | A1 Astrocyte | 0.94 | 0.00 | 0.71 | 0.09 | 0.07 | 0.04 | 0.99 | 1.00 |
| Sqstm1 | 2142.13 | Autophagy | 0.66 | 0.01 | 0.86 | 0.11 | 0.08 | 0.15 | 0.97 | 1.00 |
| Pmp22 | 846.83 | Myelogenesis | 0.53 | 0.02 | 0.57 | 0.36 | 0.11 | 0.12 | 1.00 | 0.82 |
| B4galt5 | 72.43 | Oligodendrocyte Differentiation | 0.97 | 0.04 | 0.95 | 0.39 | 0.35 | 0.35 | 1.00 | 1.00 |
| Gdpd2 | 67.21 | Astrocyte Markers | 0.93 | 0.03 | 0.25 | 0.74 | 0.32 | 0.09 | 0.78 | 0.84 |
| Fabp5 | 200.32 | Microglia-DAM Stage 1 | 0.75 | 0.01 | 0.24 | 0.41 | 0.17 | 0.04 | 0.68 | 0.91 |
| Tap1 | 141.25 | Primed Microglia | 0.60 | 0.03 | 0.85 | 0.37 | 0.49 | 0.27 | 0.95 | 0.99 |
| S100a10 | 502.72 | A2 Astrocyte | 0.68 | 0.00 | 0.82 | 0.02 | 0.02 | 0.01 | 0.97 | 1.00 |
| Aqp4 | 700.68 | Astrocyte Markers | 0.25 | 0.00 | 0.87 | 0.02 | 0.07 | 0.02 | 0.76 | 0.87 |
| Vim | 2440.13 | Macrophage Function | 0.36 | 0.00 | 0.48 | 0.01 | 0.03 | 0.02 | 1.00 | 0.64 |
| Spp1 | 121.89 | Microglia-DAM Stage 2-MGnD | 0.36 | 0.00 | 0.75 | 0.01 | 0.03 | 0.02 | 0.97 | 0.80 |
| Serping1 | 68.84 | A1 Astrocyte | 0.41 | 0.00 | 0.91 | 0.01 | 0.00 | 0.00 | 0.95 | 0.90 |
| Hmox1 | 69.25 | Primed Microglia | 0.98 | 0.00 | 0.56 | 0.00 | 0.00 | 0.00 | 0.97 | 0.98 |
| Gfap | 1998.79 | Astrocyte Markers | 0.99 | 0.00 | 0.96 | 0.00 | 0.00 | 0.00 | 1.00 | 1.00 |
| Igfbp5 | 316.56 | Insulin Signaling | 0.04 | 0.00 | 0.12 | 0.00 | 0.00 | 0.00 | 0.96 | 0.07 |
| C4a/b | 2161.98 | Astrocyte Markers | 0.38 | 0.00 | 0.50 | 0.00 | 0.00 | 0.00 | 1.00 | 0.67 |
| Serpina3n | 179.64 | Astrocyte Markers | 0.63 | 0.00 | 0.39 | 0.00 | 0.00 | 0.00 | 0.99 | 0.75 |
| Klk6 | 834.72 | Myelogenesis | 0.72 | 0.00 | 0.39 | 0.00 | 0.00 | 0.00 | 0.80 | 0.98 |
| Stat3 | 542.46 | Interferon Signaling | 0.21 | 0.00 | 0.94 | 0.00 | 0.00 | 0.00 | 0.80 | 0.74 |
| Csf1 | 647.24 | Microglia-DAM Stage 2-MGnD | 0.18 | 0.00 | 0.30 | 0.02 | 0.00 | 0.00 | 0.99 | 0.33 |
| Cntf | 100.11 | Neurogenesis | 0.77 | 0.00 | 0.45 | 0.00 | 0.00 | 0.00 | 0.98 | 0.86 |
| C3 | 131.15 | Primed Microglia | 0.90 | 0.00 | 0.75 | 0.00 | 0.00 | 0.00 | 1.00 | 0.99 |
| Trac | 73.49 | T-cell signaling | 0.55 | 0.00 | 0.59 | 0.00 | 0.01 | 0.00 | 1.00 | 0.84 |
| Gpnmb | 59.66 | Microglia-DAM Stage 1 | 0.76 | 0.00 | 0.90 | 0.05 | 0.07 | 0.06 | 1.00 | 0.99 |
| Cd74 | 201.61 | Microglia-M1 | 0.06 | 0.00 | 0.43 | 0.01 | 0.00 | 0.00 | 0.79 | 0.23 |
| Ptgis | 90.91 | NO Metabolism and Signaling | 0.24 | 0.00 | 0.80 | 0.10 | 0.02 | 0.06 | 0.89 | 0.71 |
| Nuak1 | 58.98 | Primed Microglia | 0.74 | 0.01 | 0.31 | 0.29 | 0.14 | 0.03 | 0.75 | 0.95 |
| Dusp3 | 311.92 | Neurotrophin Signaling | 0.02 | 0.00 | 0.70 | 0.04 | 0.40 | 0.02 | 0.19 | 0.39 |
| Scp2 | 116.14 | Lipid Metabolism | 0.59 | 0.04 | 0.32 | 0.14 | 0.58 | 0.77 | 0.98 | 0.68 |
| Slc38a1 | 324.88 | Glutamatergic Synapse | 0.57 | 0.02 | 0.27 | 0.07 | 0.13 | 0.61 | 0.61 | 0.97 |
| Abca1 | 765.53 | Lipid Metabolism | 0.75 | 0.02 | 0.69 | 0.16 | 0.18 | 0.35 | 0.95 | 1.00 |
| Cd9 | 2883.69 | Microglia-DAM Stage 2-MGnD | 0.66 | 0.00 | 0.22 | 0.01 | 0.03 | 0.19 | 0.60 | 0.92 |
| Lgals3 | 193.34 | Microglia-DAM Stage 2-MGnD | 0.31 | 0.00 | 0.03 | 0.01 | 0.02 | 0.69 | 0.11 | 0.69 |
| S100b | 1357.94 | Astrocyte Markers | 0.47 | 0.02 | 0.70 | 0.14 | 0.41 | 0.31 | 0.99 | 0.84 |
| Fth1 | 34777.03 | Microglia-DAM Stage 1 | 0.55 | 0.02 | 0.24 | 0.06 | 0.38 | 0.62 | 0.97 | 0.56 |
| Plekhb1 | 9402.17 | Oligodendrocyte Markers | 0.48 | 0.02 | 0.33 | 0.08 | 0.44 | 0.54 | 1.00 | 0.61 |
| Mbp | 8365.86 | Oligodendrocyte Markers | 0.55 | 0.05 | 0.42 | 0.19 | 0.64 | 0.73 | 1.00 | 0.73 |
| Eid1 | 1039.23 | Primed Microglia | 0.48 | 0.03 | 0.47 | 0.17 | 0.59 | 0.60 | 1.00 | 0.72 |
| Jun | 2399.12 | Microglia Markers | 0.03 | 0.04 | 0.24 | 0.85 | 1.00 | 0.12 | 0.10 | 0.79 |
| Fos | 3529.07 | MAPK & PI3K | 0.23 | 0.01 | 0.22 | 0.56 | 0.56 | 0.05 | 0.32 | 1.00 |
| Egr1 | 1634.52 | Microglia-Homeostatic | 0.37 | 0.04 | 0.13 | 0.93 | 0.69 | 0.07 | 0.32 | 0.95 |
| Lgmn | 1655.96 | Microglia Markers | 0.04 | 0.42 | 0.25 | 0.49 | 0.18 | 0.99 | 0.12 | 0.85 |
| Ctsb | 911.52 | Microglia-DAM Stage 1 | 0.05 | 0.11 | 0.06 | 0.09 | 0.07 | 0.99 | 0.05 | 1.00 |
| Hexb | 423.03 | Microglia-Homeostatic | 0.85 | 0.04 | 0.12 | 0.07 | 0.30 | 0.95 | 0.55 | 0.71 |
| Eif2b5 | 301.09 | Oligodendrocyte Differentiation/Maturation | 0.61 | 0.56 | 0.02 | 0.15 | 0.85 | 0.44 | 0.16 | 0.41 |
| Clta | 1284.71 | Wnt Signaling | 0.55 | 0.82 | 0.00 | 0.06 | 0.93 | 0.10 | 0.04 | 0.15 |
| Myrf | 1014.36 | Oligodendrocyte Differentiation/Maturation | 0.03 | 0.04 | 0.78 | 0.46 | 0.03 | 0.28 | 0.43 | 0.27 |
| Gsn | 2011.91 | Oligodendrocyte Function | 0.02 | 0.04 | 0.84 | 0.46 | 0.02 | 0.33 | 0.27 | 0.18 |
| Psma5 | 541.43 | Antigen Processing and Presentation | 0.16 | 0.03 | 0.15 | 0.07 | 0.89 | 0.90 | 1.00 | 0.20 |
| Ctnnb1 | 2976.13 | Wnt Signaling | 0.02 | 0.58 | 0.34 | 0.69 | 0.39 | 0.99 | 0.54 | 0.09 |
| Plcb4 | 262.42 | Wnt Signaling | 0.04 | 0.91 | 0.10 | 0.62 | 0.35 | 0.53 | 0.98 | 0.07 |
| Orc4 | 277.05 | Cell Cycle | 0.36 | 0.05 | 0.95 | 0.38 | 0.17 | 0.42 | 0.88 | 0.91 |
| Hectd1 | 365.14 | Antigen Processing and Presentation | 0.61 | 0.02 | 0.65 | 0.18 | 0.17 | 0.42 | 0.89 | 1.00 |
| Nkiras1 | 129.25 | Cytokines | 0.83 | 0.05 | 0.82 | 0.34 | 0.34 | 0.50 | 0.99 | 1.00 |
| Cdc27 | 198.37 | Cell Cycle | 0.63 | 0.01 | 0.64 | 0.10 | 0.10 | 0.25 | 0.90 | 1.00 |
| Ranbp9 | 421.01 | Neurogenesis | 0.07 | 0.02 | 0.97 | 0.28 | 0.03 | 0.26 | 0.49 | 0.46 |
| Glrb | 961.84 | Neuroactive Ligands and Receptors | 0.72 | 0.01 | 0.32 | 0.06 | 0.24 | 0.44 | 0.96 | 0.75 |
| Slc25a4 | 3017.21 | Calcium Signaling | 0.47 | 0.05 | 0.68 | 0.57 | 0.21 | 0.29 | 1.00 | 0.84 |
| Dlat | 271.06 | Glucose Metabolism | 0.41 | 0.03 | 0.58 | 0.50 | 0.14 | 0.19 | 1.00 | 0.75 |
| Acot13 | 360.95 | Lipid Metabolism | 0.34 | 0.01 | 0.68 | 0.22 | 0.05 | 0.10 | 0.97 | 0.74 |
| Braf | 210.29 | MAPK & PI3K | 0.98 | 0.04 | 0.59 | 0.57 | 0.36 | 0.22 | 0.98 | 0.97 |
| Nwd1 | 472.19 | Astrocyte Markers | 0.51 | 0.00 | 0.22 | 0.01 | 0.02 | 0.13 | 0.51 | 0.97 |
| Acsl4 | 130.97 | Neurogenesis | 0.25 | 0.02 | 0.13 | 0.04 | 0.06 | 0.78 | 0.23 | 0.99 |
| Gpm6a | 3965.57 | Neurogenesis | 0.75 | 0.01 | 0.63 | 0.11 | 0.24 | 0.28 | 1.00 | 0.93 |
| Pdhb | 779.78 | Glucose Metabolism | 0.94 | 0.00 | 0.16 | 0.29 | 0.06 | 0.01 | 0.73 | 0.67 |
| Atp2a2 | 1768.37 | Ion Transport | 0.36 | 0.00 | 0.36 | 0.06 | 0.06 | 0.01 | 0.55 | 1.00 |
| Kpna1 | 221.26 | Interferon Signaling | 0.91 | 0.04 | 0.67 | 0.51 | 0.33 | 0.25 | 1.00 | 0.98 |
| Prkcq | 272.45 | T-cell signaling | 0.23 | 0.01 | 0.12 | 0.76 | 0.58 | 0.04 | 0.22 | 0.99 |
| Rapgef2 | 124.74 | MAPK & PI3K | 0.05 | 0.04 | 0.12 | 0.07 | 0.04 | 0.96 | 0.08 | 0.98 |
| Acly | 397.75 | Lipid Metabolism | 0.48 | 0.08 | 0.02 | 0.04 | 0.80 | 0.94 | 0.50 | 0.13 |
| Myorg | 314.29 | Astrocyte Markers | 0.76 | 0.01 | 0.08 | 0.02 | 0.10 | 0.71 | 0.42 | 0.66 |
| Vegfa | 938.29 | Blood Brain Barrier | 0.56 | 0.01 | 0.47 | 0.08 | 0.10 | 0.36 | 0.77 | 1.00 |
| Gria2 | 199.75 | Neuroactive Ligands and Receptors | 0.46 | 0.01 | 0.59 | 0.09 | 0.07 | 0.26 | 0.78 | 1.00 |
| Dagla | 128.27 | Neurogenesis | 0.59 | 0.04 | 0.23 | 0.10 | 0.20 | 0.82 | 0.58 | 0.95 |
| Hsd11b1 | 320.20 | Lipid Metabolism | 0.21 | 0.02 | 0.32 | 0.08 | 0.06 | 0.55 | 0.38 | 1.00 |
| Slc6a1 | 1233.71 | Astrocyte Markers | 0.47 | 0.01 | 0.05 | 0.02 | 0.08 | 0.90 | 0.21 | 0.69 |
| Rorb | 175.27 | Neurogenesis | 0.15 | 0.05 | 0.05 | 0.05 | 0.10 | 1.00 | 0.10 | 0.96 |
| Acsl3 | 1679.20 | Lipid Metabolism | 0.37 | 0.03 | 0.07 | 0.04 | 0.12 | 0.97 | 0.21 | 0.84 |
| Ptprz1 | 1964.38 | Oligodendrocyte Differentiation/Maturation | 0.12 | 0.04 | 0.08 | 0.06 | 0.07 | 0.99 | 0.12 | 1.00 |
| Sirt5 | 64.91 | Glucose Metabolism | 0.71 | 0.04 | 0.81 | 0.43 | 0.48 | 0.28 | 0.97 | 1.00 |
| Mertk | 352.60 | Microglia-Homeostatic | 0.20 | 0.00 | 0.84 | 0.03 | 0.01 | 0.05 | 0.69 | 0.84 |
| Cd47 | 667.53 | NO Metabolism and Signaling | 0.33 | 0.02 | 0.58 | 0.13 | 0.09 | 0.39 | 0.68 | 0.99 |
| Serpini1 | 109.41 | Neurogenesis | 0.45 | 0.00 | 0.62 | 0.04 | 0.03 | 0.11 | 0.79 | 1.00 |
| Adora2b | 146.33 | Purinergic Signaling | 0.69 | 0.01 | 0.61 | 0.10 | 0.12 | 0.29 | 0.91 | 1.00 |
| Tlr3 | 169.36 | NF-kappaB Signaling | 0.21 | 0.01 | 0.13 | 0.02 | 0.03 | 0.49 | 0.22 | 1.00 |
| Gabra4 | 58.63 | GABAergic Synapse | 0.71 | 0.04 | 0.82 | 0.31 | 0.52 | 0.46 | 1.00 | 0.97 |
| Gabrb3 | 84.92 | GABAergic Synapse | 0.53 | 0.03 | 0.36 | 0.12 | 0.16 | 0.62 | 0.67 | 1.00 |
| Rab3a | 646.07 | Glutamatergic Synapse | 0.82 | 0.04 | 0.32 | 0.14 | 0.30 | 0.77 | 0.79 | 0.93 |
| Amph | 212.96 | Phagocytosis | 0.92 | 0.02 | 0.14 | 0.06 | 0.25 | 0.84 | 0.63 | 0.71 |
| Opcml | 93.77 | Neurogenesis | 0.59 | 0.03 | 0.65 | 0.19 | 0.48 | 0.45 | 1.00 | 0.88 |
| Mapk8 | 168.73 | MAPK & PI3K | 0.02 | 0.06 | 0.63 | 0.30 | 0.03 | 0.65 | 0.16 | 0.39 |
| Cables1 | 221.66 | Cell Cycle | 0.22 | 0.00 | 0.31 | 0.01 | 0.01 | 0.05 | 0.38 | 1.00 |
| Ahcyl1 | 1250.00 | Astrocyte Markers | 0.54 | 0.03 | 0.34 | 0.13 | 0.18 | 0.68 | 0.66 | 0.99 |
| Dnajc5 | 481.38 | Proteotoxic Stress | 0.04 | 0.00 | 0.11 | 0.01 | 0.01 | 0.44 | 0.07 | 0.95 |
| Gng2 | 182.94 | Serotonergic Synapse | 0.14 | 0.01 | 0.12 | 0.03 | 0.03 | 0.70 | 0.15 | 1.00 |
| Nlk | 75.27 | Wnt Signaling | 0.32 | 0.01 | 0.29 | 0.06 | 0.06 | 0.49 | 0.46 | 1.00 |
| Abi2 | 248.38 | Neurogenesis | 0.65 | 0.02 | 0.06 | 0.03 | 0.18 | 0.98 | 0.29 | 0.62 |
| Mapk1 | 896.21 | MAPK & PI3K | 0.13 | 0.05 | 0.13 | 0.97 | 0.97 | 0.09 | 0.16 | 1.00 |
| Islr2 | 124.09 | Neurogenesis | 0.31 | 0.01 | 0.93 | 0.14 | 0.35 | 0.11 | 0.83 | 0.89 |
| Rfx4 | 141.50 | Astrocyte Markers | 0.12 | 0.72 | 0.02 | 0.31 | 0.77 | 0.15 | 0.04 | 0.80 |
| Iigp1 | 136.96 | Interferon Signaling | 0.01 | 0.44 | 0.12 | 0.92 | 0.38 | 0.34 | 0.03 | 0.72 |
| Ptk2 | 362.50 | Cell Migration | 0.01 | 0.28 | 0.63 | 0.65 | 0.06 | 0.96 | 0.12 | 0.31 |
| Gad1 | 52.05 | GABAergic Synapse | 0.06 | 0.36 | 0.03 | 0.12 | 0.20 | 0.65 | 0.04 | 0.98 |
| Plaa | 85.38 | Proteotoxic Stress | 0.03 | 0.99 | 0.95 | 1.00 | 0.33 | 1.00 | 0.35 | 0.31 |
| Bhlhe40 | 872.22 | Neuronal Markers | 0.86 | 0.83 | 0.04 | 0.43 | 0.99 | 0.29 | 0.41 | 0.30 |
| Adora1 | 64.28 | Purinergic Signaling | 0.52 | 1.00 | 0.00 | 0.07 | 0.96 | 0.07 | 0.13 | 0.03 |
| Atg2b | 151.87 | Autophagy | 0.05 | 0.12 | 0.94 | 0.59 | 0.98 | 0.66 | 0.44 | 0.39 |
| Gla | 96.10 | NO Metabolism and Signaling | 0.01 | 0.75 | 0.14 | 0.80 | 0.21 | 0.55 | 0.03 | 0.60 |
| B4galt6 | 272.58 | Oligodendrocyte Differentiation | 0.44 | 0.05 | 0.17 | 0.93 | 0.70 | 0.09 | 0.41 | 0.96 |
| Ap3m2 | 189.43 | Primed Microglia | 0.12 | 0.29 | 0.04 | 0.81 | 0.97 | 0.14 | 0.08 | 0.96 |
| F11r | 433.29 | Primed Microglia | 0.44 | 0.05 | 0.00 | 0.66 | 0.73 | 0.01 | 0.04 | 0.20 |
| Klhl12 | 103.72 | Wnt Signaling | 0.90 | 0.37 | 0.04 | 0.75 | 0.86 | 0.16 | 0.44 | 0.35 |

### Figure 7-Supplemental

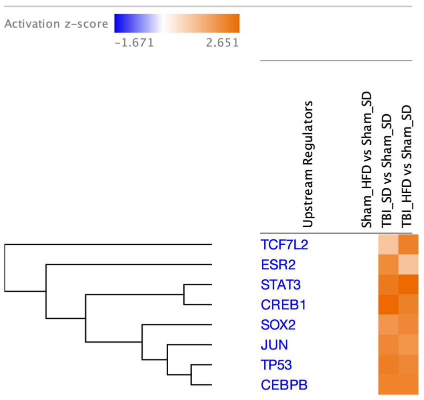

Upstream regulators – Genes (upregulated)

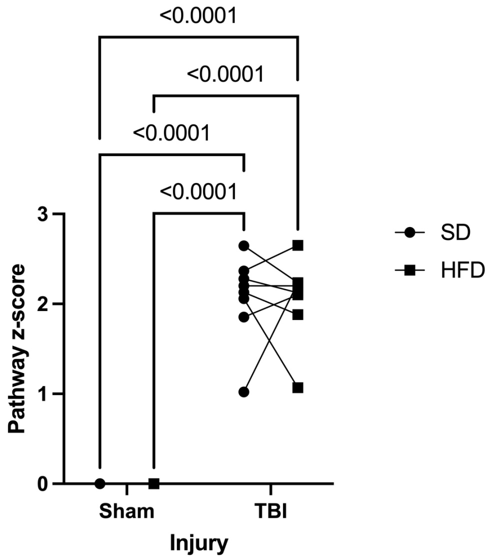

| | | | | | |
|---|---|---|---|---|---|
| Source of Variation | % of total variation | P value | P value summary | Significant? | |
| Injury | 91.72 | <0.0001 | **** | Yes | |
| Diet | 0.0002727 | 0.9757 | ns | No | |
| Injury x Diet | 0.0002727 | 0.9757 | ns | No | |
| Subject x Injury | 2.226 | | | | |
| Subject x Diet | 1.913 | | | | |
| Subject | 2.226 | | | | |
| ANOVA table | SS | DF | MS | F (DFn, DFd) | P value |
| Injury | 34.15 | 1 | 34.15 | F (1, 7) = 288.5 | P<0.0001 |
| Diet | 0.0001015 | 1 | 0.0001015 | F (1, 7) = 0.0009978 | P=0.9757 |
| Injury x Diet | 0.0001015 | 1 | 0.0001015 | F (1, 7) = 0.0009978 | P=0.9757 |

**Figure 8-Supplemental**

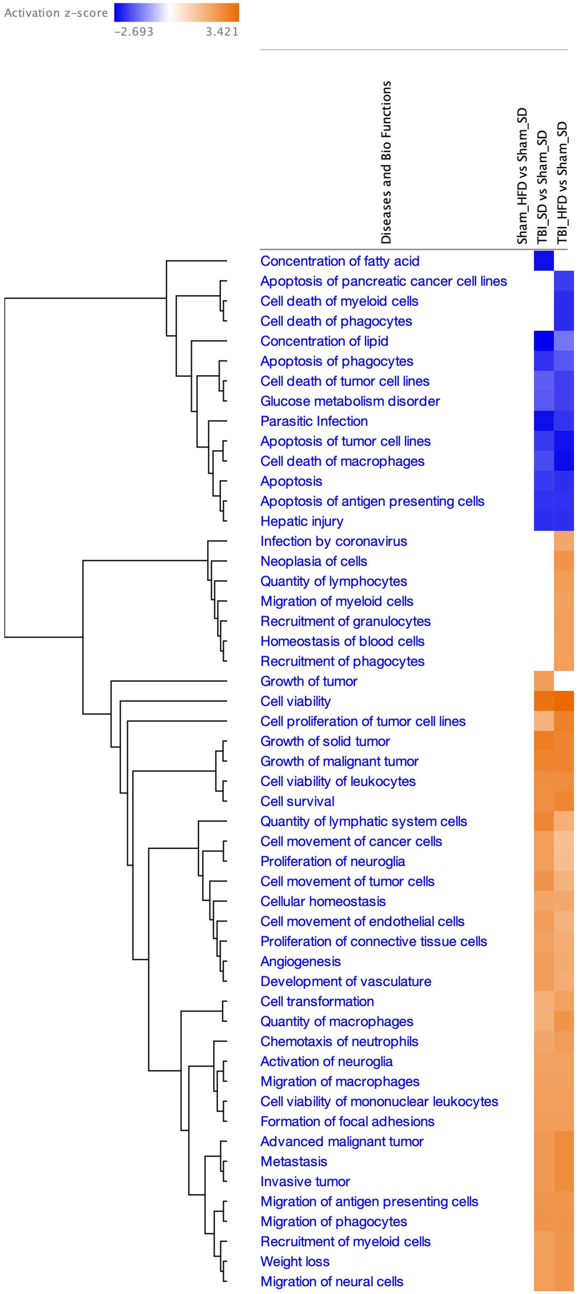

#### Diseases (upregulated)

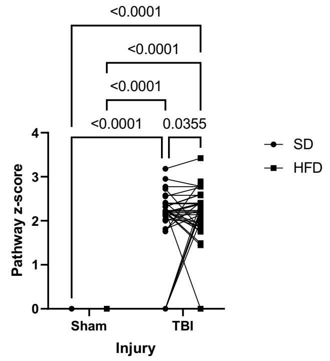

| | | | | | |
|---|---|---|---|---|---|
| Source of Variation | % of total variation | P value | P value summary | Significant? | |
| Injury | 77.53 | <0.0001 | **** | Yes | |
| Diet | 0.5334 | 0.0523 | ns | No | |
| Injury x Diet | 0.5334 | 0.0523 | ns | No | |
| Pathway x Injury | 5.792 | | | | |
| Pathway x Diet | 4.909 | | | | |
| Pathway | 5.792 | | | | |
| ANOVA table | SS | DF | MS | F (DFn, DFd) | P value |
| Injury | 157.9 | 1 | 157.9 | F (1, 37) = 495.3 | P<0.0001 |
| Diet | 1.087 | 1 | 1.087 | F (1, 37) = 4.020 | P=0.0523 |
| Injury x Diet | 1.087 | 1 | 1.087 | F (1, 37) = 4.020 | P=0.0523 |

#### Diseases (downregulated)

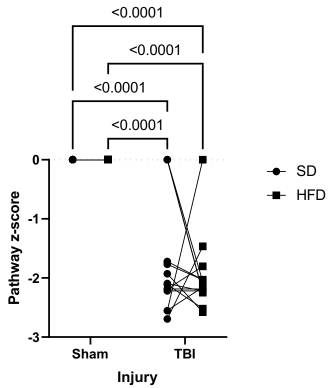

| | | | | | |
|---|---|---|---|---|---|
| Source of Variation | % of total variation | P value | P value summary | Significant? | |
| Injury | 72.75 | <0.0001 | **** | Yes | |
| Diet | 0.3818 | 0.4590 | ns | No | |
| Injury x Diet | 0.3818 | 0.4590 | ns | No | |
| Pathway x Injury | 4.722 | | | | |
| Pathway x Diet | 8.522 | | | | |
| Pathway | 4.722 | | | | |
| ANOVA table | SS | DF | MS | F (DFn, DFd) | P value |
| Injury | 47.87 | 1 | 47.87 | F (1, 13) = 200.3 | P<0.0001 |
| Diet | 0.2513 | 1 | 0.2513 | F (1, 13) = 0.5825 | P=0.4590 |
| Injury x Diet | 0.2513 | 1 | 0.2513 | F (1, 13) = 0.5825 | P=0.4590 |

**Figure 9-Supplemental**

### Figure 10-Supplemental

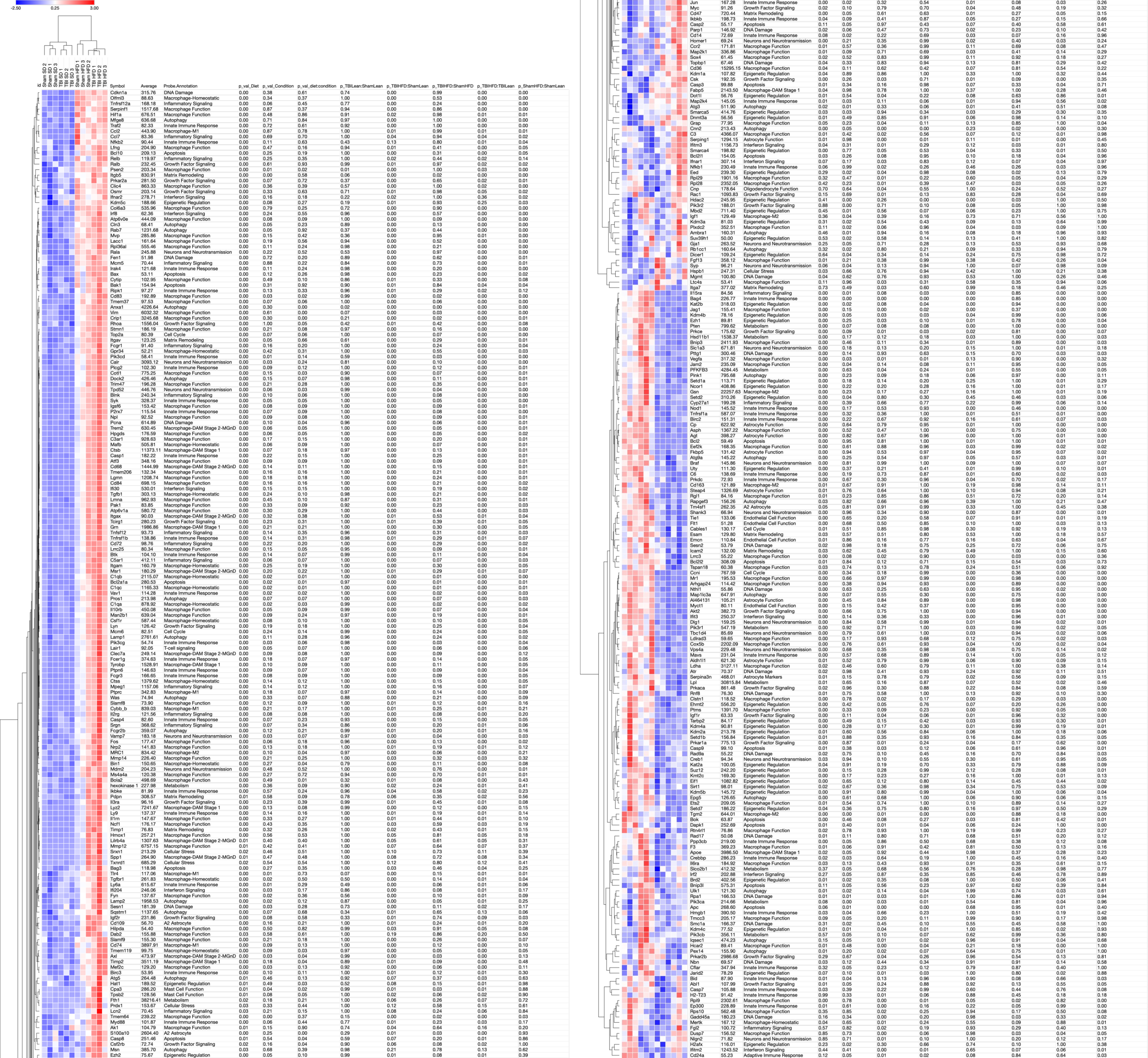

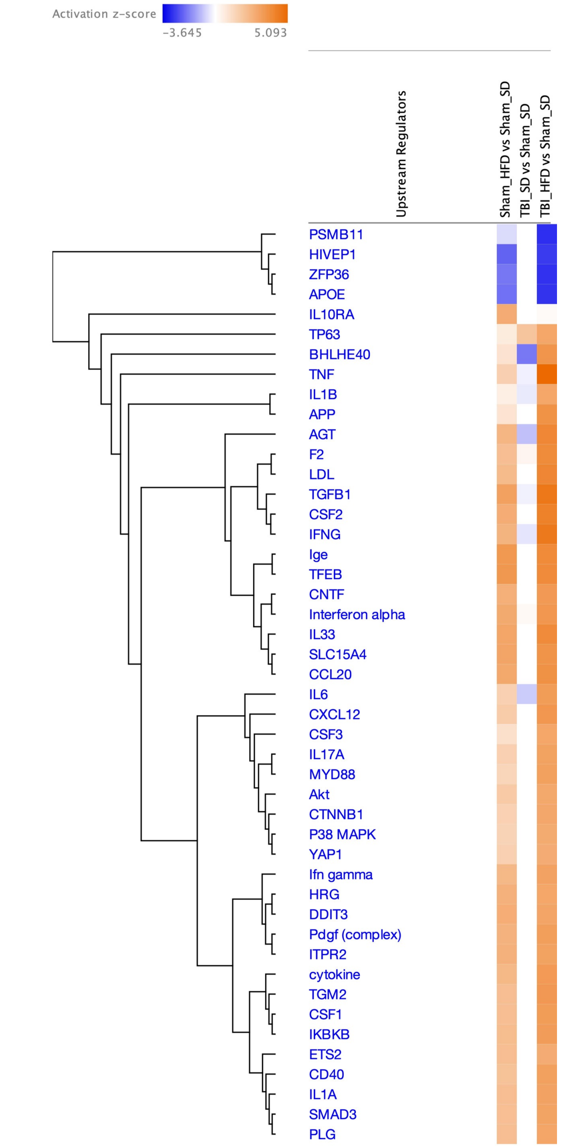

#### Upstream regulators – Genes (upregulated)

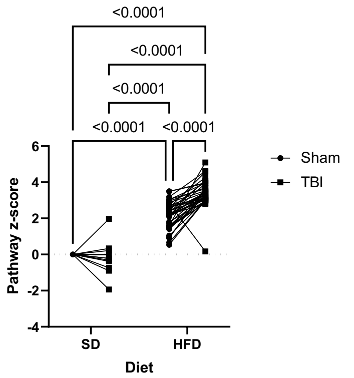

| | | | | | |
|---|---|---|---|---|---|
| Source of Variation | % of total variation | P value | P value summary | Significant? | |
| Diet | 79.66 | <0.0001 | **** | Yes | |
| Injury | 3.446 | <0.0001 | **** | Yes | |
| Diet x Injury | 4.076 | <0.0001 | **** | Yes | |
| Pathway x Diet | 3.955 | | | | |
| Pathway x Injury | 2.456 | | | | |
| Pathway | 3.364 | | | | |
| ANOVA table | SS | DF | MS | F (DFn, DFd) | P value |
| Diet | 331.5 | 1 | 331.5 | F (1, 41) = 825.9 | P<0.0001 |
| Injury | 14.34 | 1 | 14.34 | F (1, 41) = 57.52 | P<0.0001 |
| Diet x Injury | 16.96 | 1 | 16.96 | F (1, 41) = 54.95 | P<0.0001 |

#### Upstream regulators – Genes (downregulated)

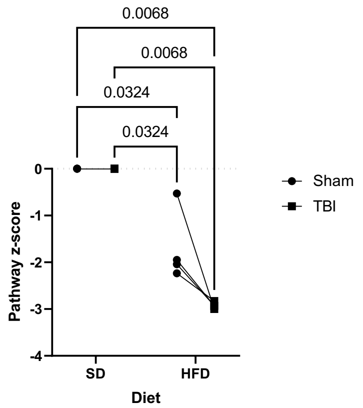

| | | | | | |
|---|---|---|---|---|---|
| Source of Variation | % of total variation | P value | P value summary | Significant? | |
| Diet | 81.28 | 0.0011 | ** | Yes | |
| Injury | 5.814 | 0.0611 | ns | No | |
| Diet x Injury | 5.814 | 0.0611 | ns | No | |
| Pathway x Diet | 1.511 | | | | |
| Pathway x Injury | 2.035 | | | | |
| Pathway | 1.511 | | | | |
| ANOVA table | SS | DF | MS | F (DFn, DFd) | P value |
| Diet | 21.26 | 1 | 21.26 | F (1, 3) = 161.4 | P=0.0011 |
| Injury | 1.521 | 1 | 1.521 | F (1, 3) = 8.572 | P=0.0611 |
| Diet x Injury | 1.521 | 1 | 1.521 | F (1, 3) = 8.572 | P=0.0611 |

**Figure 11-Supplemental**

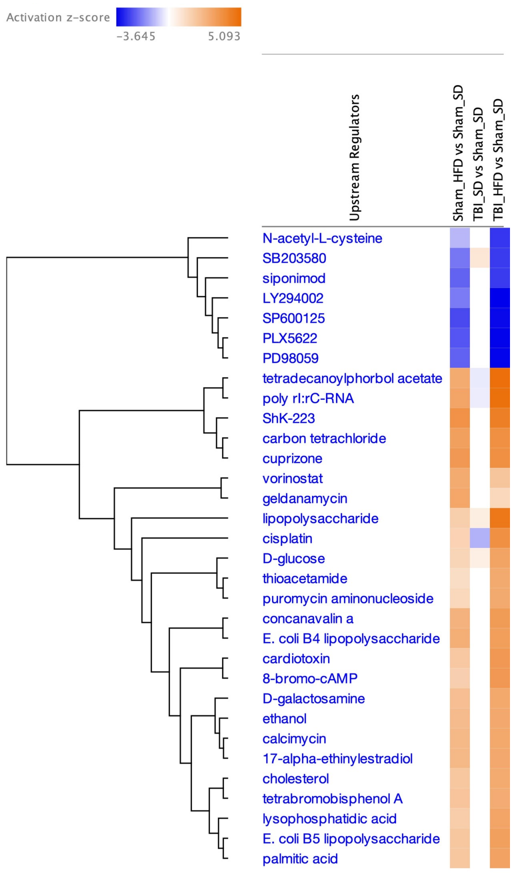

#### Upstream regulators – Drugs (upregulated)

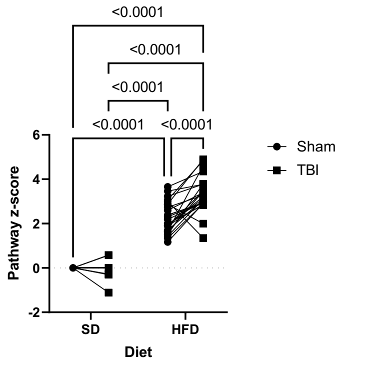

| | | | | | |
|---|---|---|---|---|---|
| Source of Variation | % of total variation | P value | P value summary | Significant? | |
| Diet | 82.22 | <0.0001 | **** | Yes | |
| Injury | 2.685 | <0.0001 | **** | Yes | |
| Diet x Injury | 2.918 | <0.0001 | **** | Yes | |
| Pathway x Diet | 3.833 | | | | |
| Pathway x Injury | 2.423 | | | | |
| Pathway | 3.328 | | | | |
| ANOVA table | SS | DF | MS | F (DFn, DFd) | P value |
| Diet | 196.3 | 1 | 196.3 | F (1, 24) = 514.8 | P<0.0001 |
| Injury | 6.408 | 1 | 6.408 | F (1, 24) = 26.59 | P<0.0001 |
| Diet x Injury | 6.965 | 1 | 6.965 | F (1, 24) = 27.00 | P<0.0001 |

#### Upstream regulators – Drugs (downregulated)

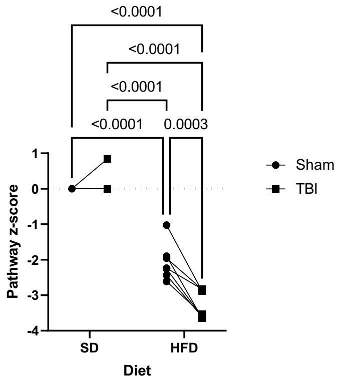

| | | | | | |
|---|---|---|---|---|---|
| Source of Variation | % of total variation | P value | P value summary | Significant? | |
| Diet | 86.08 | <0.0001 | **** | Yes | |
| Injury | 3.468 | 0.0041 | ** | Yes | |
| Diet x Injury | 5.159 | 0.0003 | *** | Yes | |
| Pathway x Diet | 1.478 | | | | |
| Pathway x Injury | 1.028 | | | | |
| Pathway | 2.240 | | | | |
| ANOVA table | SS | DF | MS | F (DFn, DFd) | P value |
| Diet | 52.09 | 1 | 52.09 | F (1, 6) = 349.5 | P<0.0001 |
| Injury | 2.099 | 1 | 2.099 | F (1, 6) = 20.23 | P=0.0041 |
| Diet x Injury | 3.122 | 1 | 3.122 | F (1, 6) = 56.21 | P=0.0003 |

**Figure 12-Supplemental**

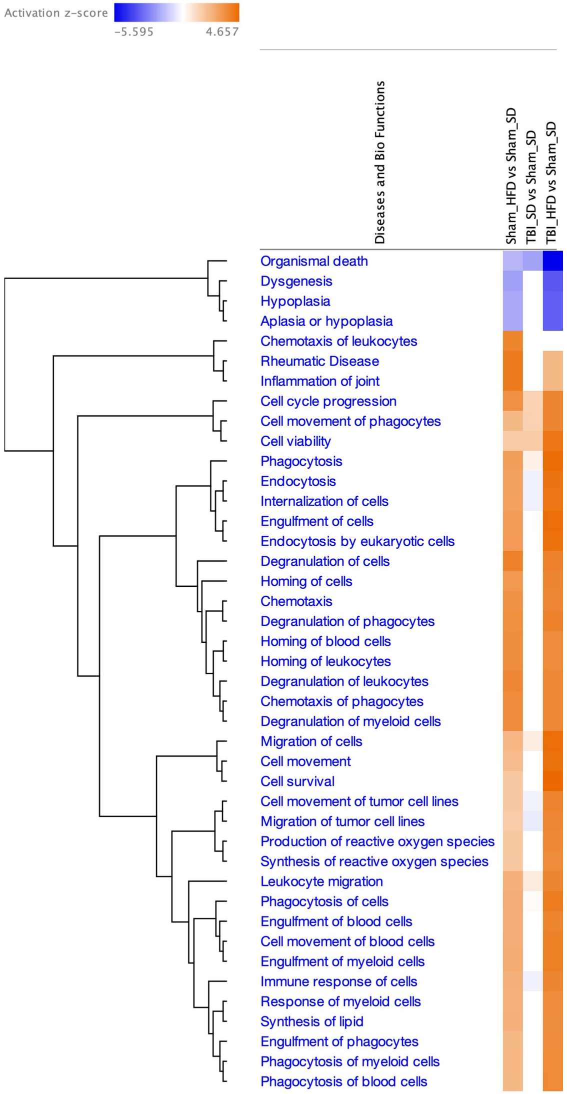

#### Diseases (upregulated)

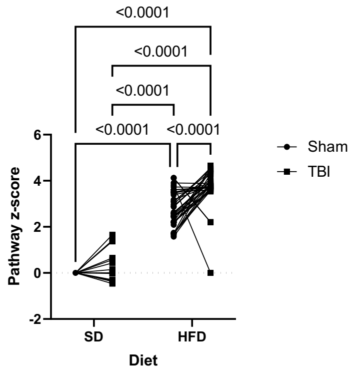

| | | | | | |
|---|---|---|---|---|---|
| Source of Variation | % of total variation | P value | P value summary | Significant? | |
| Diet | 84.80 | <0.0001 | **** | Yes | |
| Injury | 2.318 | <0.0001 | **** | Yes | |
| Diet x Injury | 1.414 | 0.0005 | *** | Yes | |
| Pathway x Diet | 1.821 | | | | |
| Pathway x Injury | 4.205 | | | | |
| Pathway | 1.856 | | | | |
| ANOVA table | SS | DF | MS | F (DFn, DFd) | P value |
| Diet | 388.5 | 1 | 388.5 | F (1, 37) = 1723 | P<0.0001 |
| Injury | 10.62 | 1 | 10.62 | F (1, 37) = 20.39 | P<0.0001 |
| Diet x Injury | 6.479 | 1 | 6.479 | F (1, 37) = 14.59 | P=0.0005 |

#### Diseases (downregulated)

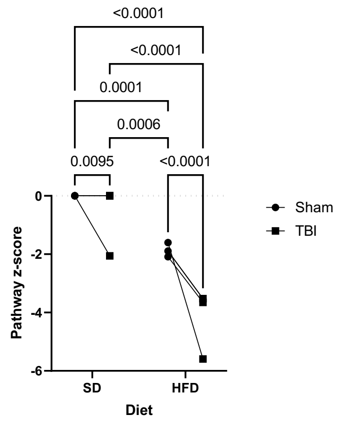

| | | | | | |
|---|---|---|---|---|---|
| Source of Variation | % of total variation | P value | P value summary | Significant? | |
| Diet | 63.91 | <0.0001 | **** | Yes | |
| Injury | 16.03 | 0.0920 | ns | No | |
| Diet x Injury | 6.186 | 0.0003 | *** | Yes | |
| Pathway x Diet | 0.09986 | | | | |
| Pathway x Injury | 8.034 | | | | |
| Pathway | 5.696 | | | | |
| ANOVA table | SS | DF | MS | F (DFn, DFd) | P value |
| Diet | 29.47 | 1 | 29.47 | F (1, 3) = 1920 | P<0.0001 |
| Injury | 7.390 | 1 | 7.390 | F (1, 3) = 5.984 | P=0.0920 |
| Diet x Injury | 2.853 | 1 | 2.853 | F (1, 3) = 413.2 | P=0.0003 |

**Figure 13-Supplemental**

### A Microglia – Fig. 6 & 7

#### Homeostatic Associated Microglia

| | | | | | |
|---|---|---|---|---|---|
| Two-way ANOVA | Ordinary | | | | |
| Alpha | 0.05 | | | | |
| Source of Variation | % of total variation | P value | P value summary | Significant? | |
| Interaction | 1.591 | 0.3838 | ns | No | |
| Diet | 9.135 | 0.0583 | ns | No | |
| Injury | 74.28 | 0.0002 | *** | Yes | |
| ANOVA table | SS | DF | MS | F (DFn, DFd) | P value |
| Interaction | 76.58 | 1 | 76.58 | F (1, 8) = 0.8490 | P=0.3838 |
| Diet | 439.6 | 1 | 439.6 | F (1, 8) = 4.874 | P=0.0583 |
| Injury | 3574 | 1 | 3574 | F (1, 8) = 39.63 | P=0.0002 |
| Residual | 721.6 | 8 | 90.20 | | |

#### Autophagy Associated Microglia

| | | | | | |
|---|---|---|---|---|---|
| Two-way ANOVA | Ordinary | | | | |
| Alpha | 0.05 | | | | |
| Source of Variation | % of total variation | P value | P value summary | Significant? | |
| Interaction | 0.001490 | 0.9722 | ns | No | |
| Diet | 0.08478 | 0.7932 | ns | No | |
| Injury | 90.68 | <0.0001 | **** | Yes | |
| ANOVA table | SS | DF | MS | F (DFn, DFd) | P value |
| Interaction | 0.01219 | 1 | 0.01219 | F (1, 8) = 0.001292 | P=0.9722 |
| Diet | 0.6933 | 1 | 0.6933 | F (1, 8) = 0.07348 | P=0.7932 |
| Injury | 741.6 | 1 | 741.6 | F (1, 8) = 78.60 | P<0.0001 |
| Residual | 75.48 | 8 | 9.435 | | |

#### Interferon Response Microglia

| | | | | | |
|---|---|---|---|---|---|
| Two-way ANOVA | Ordinary | | | | |
| Alpha | 0.05 | | | | |
| Source of Variation | % of total variation | P value | P value summary | Significant? | |
| Interaction | 8.438 | 0.2130 | ns | No | |
| Diet | 0.02917 | 0.9386 | ns | No | |
| Injury | 54.66 | 0.0088 | ** | Yes | |
| ANOVA table | SS | DF | MS | F (DFn, DFd) | P value |
| Interaction | 40.67 | 1 | 40.67 | F (1, 8) = 1.831 | P=0.2130 |
| Diet | 0.1406 | 1 | 0.1406 | F (1, 8) = 0.006327 | P=0.9386 |
| Injury | 263.5 | 1 | 263.5 | F (1, 8) = 11.86 | P=0.0088 |
| Residual | 177.7 | 8 | 22.22 | | |

#### Disease Associated Microglia

| | | | | | |
|---|---|---|---|---|---|
| Two-way ANOVA | Ordinary | | | | |
| Alpha | 0.05 | | | | |
| Source of Variation | % of total variation | P value | P value summary | Significant? | |
| Interaction | 0.01600 | 0.8514 | ns | No | |
| Diet | 0.2044 | 0.5088 | ns | No | |
| Injury | 96.36 | <0.0001 | **** | Yes | |
| ANOVA table | SS | DF | MS | F (DFn, DFd) | P value |
| Interaction | 1.279 | 1 | 1.279 | F (1, 8) = 0.03743 | P=0.8514 |
| Diet | 16.34 | 1 | 16.34 | F (1, 8) = 0.4783 | P=0.5088 |
| Injury | 7702 | 1 | 7702 | F (1, 8) = 225.4 | P<0.0001 |
| Residual | 273.3 | 8 | 34.17 | | |

#### Pro-inflammatory-like Microglia

| | | | | | |
|---|---|---|---|---|---|
| Two-way ANOVA | Ordinary | | | | |
| Alpha | 0.05 | | | | |
| Source of Variation | % of total variation | P value | P value summary | Significant? | |
| Interaction | 2.816 | 0.3363 | ns | No | |
| Diet | 0.5815 | 0.6544 | ns | No | |
| Injury | 75.07 | 0.0007 | *** | Yes | |
| ANOVA table | SS | DF | MS | F (DFn, DFd) | P value |
| Interaction | 13.64 | 1 | 13.64 | F (1, 8) = 1.046 | P=0.3363 |
| Diet | 2.818 | 1 | 2.818 | F (1, 8) = 0.2161 | P=0.6544 |
| Injury | 363.7 | 1 | 363.7 | F (1, 8) = 27.90 | P=0.0007 |
| Residual | 104.3 | 8 | 13.04 | | |

#### Microglia Markers

| | | | | | |
|---|---|---|---|---|---|
| Two-way ANOVA | Ordinary | | | | |
| Alpha | 0.05 | | | | |
| Source of Variation | % of total variation | P value | P value summary | Significant? | |
| Interaction | 2.343 | 0.2989 | ns | No | |
| Diet | 17.07 | 0.0171 | * | Yes | |
| Injury | 65.39 | 0.0004 | *** | Yes | |
| ANOVA table | SS | DF | MS | F (DFn, DFd) | P value |
| Interaction | 6.990 | 1 | 6.990 | F (1, 8) = 1.234 | P=0.2989 |
| Diet | 50.92 | 1 | 50.92 | F (1, 8) = 8.987 | P=0.0171 |
| Injury | 195.0 | 1 | 195.0 | F (1, 8) = 34.43 | P=0.0004 |
| Residual | 45.32 | 8 | 5.665 | | |

#### Canonical genes (upregulated)

| | | | | |
|---|---|---|---|---|
| Source of Variation | % of total variation | P value | P value summary | Significant? |
| Injury | 73.34 | <0.0001 | **** | Yes |
| Diet | 2.919 | 0.0055 | ** | Yes |
| Injury x Diet | 2.919 | 0.0055 | ** | Yes |
| Pathway x Injury | 5.874 | | | |
| Pathway x Diet | 4.540 | | | |
| Pathway | 5.874 | | | |
| ANOVA table | SS | DF | MS | F (DFn, DFd) |
| Injury | 57.09 | 1 | 57.09 | F (1, 16) = 199.8 |
| Diet | 2.272 | 1 | 2.272 | F (1, 16) = 10.29 |
| Injury x Diet | 2.272 | 1 | 2.272 | F (1, 16) = 10.29 |
| Pathway x Injury | 4.573 | 16 | 0.2858 | |
| Pathway x Diet | 3.534 | 16 | 0.2209 | |
| Pathway | 4.573 | 16 | 0.2858 | |
| Residual | 3.534 | 16 | 0.2209 | |

#### Canonical genes (downregulated)

| | | | | | |
|---|---|---|---|---|---|
| Source of Variation | % of total variation | P value | P value summary | Significant? | |
| Injury | 97.29 | <0.0001 | **** | Yes | |
| Diet | 0.1164 | 0.3369 | ns | No | |
| Injury x Diet | 0.1164 | 0.3369 | ns | No | |
| Pathway x Injury | 0.8481 | | | | |
| Pathway x Diet | 0.3920 | | | | |
| Pathway | 0.8481 | | | | |
| ANOVA table | SS | DF | MS | F (DFn, DFd) | P value |
| Injury | 24.19 | 1 | 24.19 | F (1, 4) = 458.9 | P<0.0001 |
| Diet | 0.02896 | 1 | 0.02896 | F (1, 4) = 1.188 | P=0.3369 |
| Injury x Diet | 0.02896 | 1 | 0.02896 | F (1, 4) = 1.188 | P=0.3369 |
| Pathway x Injury | 0.2109 | 4 | 0.05272 | | |
| Pathway x Diet | 0.09747 | 4 | 0.02437 | | |
| Pathway | 0.2109 | 4 | 0.05272 | | |
| Residual | 0.09747 | 4 | 0.02437 | | |

### B Flow Through – Fig. 8

#### Flowthrough Neurons

| | | | | | |
|---|---|---|---|---|---|
| Two-way ANOVA | Ordinary | | | | |
| Alpha | 0.05 | | | | |
| Source of Variation | % of total variation | P value | P value summary | Significant? | |
| Interaction | 17.35 | 0.0181 | * | Yes | |
| Diet | 5.702 | 0.1278 | ns | No | |
| Injury | 61.15 | 0.0005 | *** | Yes | |
| ANOVA table | SS | DF | MS | F (DFn, DFd) | P value |
| Interaction | 194.4 | 1 | 194.4 | F (1, 8) = 8.781 | P=0.0181 |
| Diet | 63.88 | 1 | 63.88 | F (1, 8) = 2.886 | P=0.1278 |
| Injury | 685.1 | 1 | 685.1 | F (1, 8) = 30.95 | P=0.0005 |
| Residual | 177.1 | 8 | 22.13 | | |

#### Flowthrough Oligodendrocytes

| | | | | | |
|---|---|---|---|---|---|
| Two-way ANOVA | Ordinary | | | | |
| Alpha | 0.05 | | | | |
| Source of Variation | % of total variation | P value | P value summary | Significant? | |
| Interaction | 13.26 | 0.0485 | * | Yes | |
| Diet | 9.273 | 0.0877 | ns | No | |
| Injury | 57.85 | 0.0013 | ** | Yes | |
| ANOVA table | SS | DF | MS | F (DFn, DFd) | P value |
| Interaction | 10.47 | 1 | 10.47 | F (1, 8) = 5.410 | P=0.0485 |
| Diet | 7.322 | 1 | 7.322 | F (1, 8) = 3.782 | P=0.0877 |
| Injury | 45.68 | 1 | 45.68 | F (1, 8) = 23.60 | P=0.0013 |
| Residual | 15.49 | 8 | 1.936 | | |

#### Flowthrough Astrocytes

| | | | | | |
|---|---|---|---|---|---|
| Two-way ANOVA | Ordinary | | | | |
| Alpha | 0.05 | | | | |
| Source of Variation | % of total variation | P value | P value summary | Significant? | |
| Interaction | 12.83 | 0.0777 | ns | No | |
| Diet | 3.150 | 0.3455 | ns | No | |
| Injury | 58.93 | 0.0025 | ** | Yes | |
| ANOVA table | SS | DF | MS | F (DFn, DFd) | P value |
| Interaction | 95.47 | 1 | 95.47 | F (1, 8) = 4.091 | P=0.0777 |
| Diet | 23.44 | 1 | 23.44 | F (1, 8) = 1.005 | P=0.3455 |
| Injury | 438.6 | 1 | 438.6 | F (1, 8) = 18.79 | P=0.0025 |
| Residual | 186.7 | 8 | 23.34 | | |

### C Adipose Tissue – Fig. 10 &11

#### Homeostatic Associated Macrophage

| | | | | | |
|---|---|---|---|---|---|
| Two-way ANOVA | Ordinary | | | | |
| Alpha | 0.05 | | | | |
| Source of Variation | % of total variation | P value | P value summary | Significant? | |
| Interaction | 9.361 | 0.0284 | * | Yes | |
| Diet | 73.80 | <0.0001 | **** | Yes | |
| Injury | 6.324 | 0.0596 | ns | No | |
| ANOVA table | SS | DF | MS | F (DFn, DFd) | P value |
| Interaction | 182.0 | 1 | 182.0 | F (1, 8) = 7.123 | P=0.0284 |
| Diet | 1435 | 1 | 1435 | F (1, 8) = 56.16 | P<0.0001 |
| Injury | 123.0 | 1 | 123.0 | F (1, 8) = 4.813 | P=0.0596 |
| Residual | 204.5 | 8 | 25.56 | | |

#### Disease Associated Macrophage

| | | | | | |
|---|---|---|---|---|---|
| Two-way ANOVA | Ordinary | | | | |
| Alpha | 0.05 | | | | |
| Source of Variation | % of total variation | P value | P value summary | Significant? | |
| Interaction | 5.465 | 0.1190 | ns | No | |
| Diet | 70.91 | 0.0002 | *** | Yes | |
| Injury | 9.276 | 0.0525 | ns | No | |
| ANOVA table | SS | DF | MS | F (DFn, DFd) | P value |
| Interaction | 105.7 | 1 | 105.7 | F (1, 8) = 3.048 | P=0.1190 |
| Diet | 1372 | 1 | 1372 | F (1, 8) = 39.55 | P=0.0002 |
| Injury | 179.4 | 1 | 179.4 | F (1, 8) = 5.173 | P=0.0525 |
| Residual | 277.5 | 8 | 34.69 | | |

#### Macrophage Function

| | | | | | |
|---|---|---|---|---|---|
| Two-way ANOVA | Ordinary | | | | |
| Alpha | 0.05 | | | | |
| Source of Variation | % of total variation | P value | P value summary | Significant? | |
| Interaction | 7.680 | 0.0656 | ns | No | |
| Diet | 59.71 | 0.0003 | *** | Yes | |
| Injury | 19.09 | 0.0099 | ** | Yes | |
| ANOVA table | SS | DF | MS | F (DFn, DFd) | P value |
| Interaction | 1227 | 1 | 1227 | F (1, 8) = 4.545 | P=0.0656 |
| Diet | 9538 | 1 | 9538 | F (1, 8) = 35.34 | P=0.0003 |
| Injury | 3050 | 1 | 3050 | F (1, 8) = 11.30 | P=0.0099 |
| Residual | 2159 | 8 | 269.9 | | |

#### Inflammatory Signaling

| | | | | | |
|---|---|---|---|---|---|
| Two-way ANOVA | Ordinary | | | | |
| Alpha | 0.05 | | | | |
| Source of Variation | % of total variation | P value | P value summary | Significant? | |
| Interaction | 3.029 | 0.3567 | ns | No | |
| Diet | 64.53 | 0.0020 | ** | Yes | |
| Injury | 7.116 | 0.1722 | ns | No | |
| ANOVA table | SS | DF | MS | F (DFn, DFd) | P value |
| Interaction | 40.46 | 1 | 40.46 | F (1, 8) = 0.9567 | P=0.3567 |
| Diet | 861.9 | 1 | 861.9 | F (1, 8) = 20.38 | P=0.0020 |
| Injury | 95.05 | 1 | 95.05 | F (1, 8) = 2.248 | P=0.1722 |
| Residual | 338.3 | 8 | 42.29 | | |

#### Innate Immunity

| | | | | | |
|---|---|---|---|---|---|
| Two-way ANOVA | Ordinary | | | | |
| Alpha | 0.05 | | | | |
| Source of Variation | % of total variation | P value | P value summary | Significant? | |
| Interaction | 17.87 | 0.0425 | * | Yes | |
| Diet | 12.69 | 0.0768 | ns | No | |
| Injury | 44.83 | 0.0051 | ** | Yes | |
| ANOVA table | SS | DF | MS | F (DFn, DFd) | P value |
| Interaction | 807.4 | 1 | 807.4 | F (1, 8) = 5.806 | P=0.0425 |
| Diet | 573.5 | 1 | 573.5 | F (1, 8) = 4.124 | P=0.0768 |
| Injury | 2026 | 1 | 2026 | F (1, 8) = 14.57 | P=0.0051 |
| Residual | 1113 | 8 | 139.1 | | |

#### Pro-inflammatory

| | | | | | |
|---|---|---|---|---|---|
| Two-way ANOVA | Ordinary | | | | |
| Alpha | 0.05 | | | | |
| Source of Variation | % of total variation | P value | P value summary | Significant? | |
| Interaction | 3.056 | 0.1129 | ns | No | |
| Diet | 82.03 | <0.0001 | **** | Yes | |
| Injury | 7.194 | 0.0258 | * | Yes | |
| ANOVA table | SS | DF | MS | F (DFn, DFd) | P value |
| Interaction | 7.134 | 1 | 7.134 | F (1, 8) = 3.168 | P=0.1129 |
| Diet | 191.5 | 1 | 191.5 | F (1, 8) = 85.04 | P<0.0001 |
| Injury | 16.79 | 1 | 16.79 | F (1, 8) = 7.458 | P=0.0258 |
| Residual | 18.01 | 8 | 2.252 | | |

#### Canonical genes (upregulated)

| | | | | | |
|---|---|---|---|---|---|
| Source of Variation | % of total variation | P value | P value summary | Significant? | |
| Diet | 86.09 | <0.0001 | **** | Yes | |
| Injury | 2.593 | <0.0001 | **** | Yes | |
| Diet x Injury | 1.890 | <0.0001 | **** | Yes | |
| Pathway x Diet | 3.463 | | | | |
| Pathway x Injury | 1.131 | | | | |
| Pathway | 3.742 | | | | |
| ANOVA table | SS | DF | MS | F (DFn, DFd) | P value |
| Diet | 348.6 | 1 | 348.6 | F (1, 52) = 1293 | P<0.0001 |
| Injury | 10.50 | 1 | 10.50 | F (1, 52) = 119.2 | P<0.0001 |
| Diet x Injury | 7.655 | 1 | 7.655 | F (1, 52) = 90.09 | P<0.0001 |
| Pathway x Diet | 14.02 | 52 | 0.2696 | | |
| Pathway x Injury | 4.578 | 52 | 0.08805 | | |
| Pathway | 15.15 | 52 | 0.2914 | | |
| Residual | 4.418 | 52 | 0.08497 | | |

**Figure 14-Supplemental**
